# Identification of the 216kbp gene cluster and structure elucidation of gargantulides B and C, new complex 52-membered macrolides from *Amycolatopsis* sp

**DOI:** 10.1101/2021.08.11.455926

**Authors:** Daniel Carretero-Molina, Francisco Javier Ortiz-López, Tetiana Gren, Daniel Oves-Costales, Jesús Martín, Fernando Román-Hurtado, Tue Sparholt Jørgensen, Mercedes de la Cruz, Caridad Díaz, Francisca Vicente, Kai Blin, Fernando Reyes, Tilmann Weber, Olga Genilloud

## Abstract

Gargantulides B and C, two new and highly complex 52-membered glycosylated macrolactones, were isolated from *Amycolatopsis* sp. strain CA-230715 during an antibacterial screening campaign. The structures of these giant macrolides were elucidated by 2D NMR spectroscopy and shown to be related to gargantulide A, although containing additional *β*-glucopyranose and/or *α*-arabinofuranose monosaccharides separately attached to their backbones. Genome sequencing allowed the identification of a strikingly large 216 kbp biosynthetic gene cluster, among the largest type I PKS clusters described so far, and the proposal of a biosynthetic pathway for gargantulides A-C. Additionally, genes putatively responsible for the biosynthesis of the amino sugar *β*-3,6-deoxy-3-methylamino glucose, reported exclusively in gargantulide macrolides, were also found in the cluster and described in this work. The absolute configurations of gargantulides B and C were assigned based on a combination of NMR and bioinformatics analysis of ketoreductase and enoylreductase domains within the multimodular type I PKS. Furthermore, the absolute stereochemistry of the related macrolide gargantulide A has now been revised and completed. Gargantulides B and C display potent antibacterial activity against a set of drug-resistant Gram-positive bacteria and moderate activity against the clinically relevant Gram-negative pathogen *Acinetobacter baumannii*.

## Introduction

Natural products remain a source of privileged chemical scaffolds with high structural diversity and prominent biological activities which can serve to address urgent clinical needs (1). Among them, macrolides are a particularly large and widespread family of secondary metabolites mainly produced by actinomycetes (2, 3), and typically encoded by multimodular type I polyketide synthase (T1PKS) biosynthetic gene clusters (BGC). The best known macrolide is erythromycin, not only relevant as an antibacterial agent, but also as a model to study the biosynthesis of macrolide antibiotics (4). Other clinically relevant members of this family are, for example, amphotericin B, avermectin or rapamycin, which cover from antimicrobial or insecticide to immunosuppressant activities (5, 6). In the last years, some of the largest and more complex macrocyclic polyketides ever isolated from actinomycetes have been reported, including quinolidomicin A_1_ (60-membered macrolactone) (7), and stambomycins A-D (51-membered) (8). Establishing the structures of such complex and stereochemically rich large polyol macrolides has usually been extremely complicated and implied tedious chemical approaches (9–12). Fortunately, the advances on genome mining tools and the combination of NMR and bioinformatics gene cluster analysis of the T1PKS genes have arisen as a powerful approach to assign the stereochemistry of these complex polyol macrolides (13–15), greatly facilitating this task.

Recently, the discovery of gargantulide A, a complex 52-membered macrolide with antibacterial activity against Gram-positive pathogens isolated from a *Streptomyces* sp. strain was reported (16). However, no genetic information about the biosynthesis of this macrolactone has been provided to date. Based only on NMR analysis and given its structural complexity, the stereochemistry assignment of gargantulide A could not be fully achieved in the original work. During a screening program targeting a panel of Gram-negative pathogens, culture extracts of the strain *Amycolatopsis* sp. CA-230715 displayed selective antimicrobial activity against *Acinetobacter baumannii*. A bioassay-guided purification process led to the isolation of gargantulides B (**1**) and C (**2**) (Figure 1), two new giant glycosylated macrolactones structurally related to their formerly isolated congener gargantulide A (**3**). The planar structures of the new bioactive macrolides were fully elucidated by 2D NMR spectroscopy and were shown to contain the same 52-membered polyol ring and 22-membered side chain as that reported for gargantulide A, although being glycosylated with additional glucose and/or arabinofuranose sugars at their backbones. The BGC putatively responsible for the biosynthesis of gargantulides B and C (*gar*) was identified in the genome of the producing strain and it was reasoned to be responsible for the production of gargantulide A. The gene cluster codes for a giant T1PKS comprising 10 subunits, harboring 35 modules and PKS-genes spanning approximately 180 kbp. Thus, it represents the largest macrolide-encoding BGC discovered so far and, with an approximate size of 216 kbp, the gene cluster is placed among the largest BGCs described to date. Interestingly, the gene cluster also encodes the genes presumably responsible for the biosynthesis of the rare amino sugar *β*-3,6-deoxy-3-methylamino glucose (maG), which is present exclusively in gargantulides A-C.

**Figure 1.**
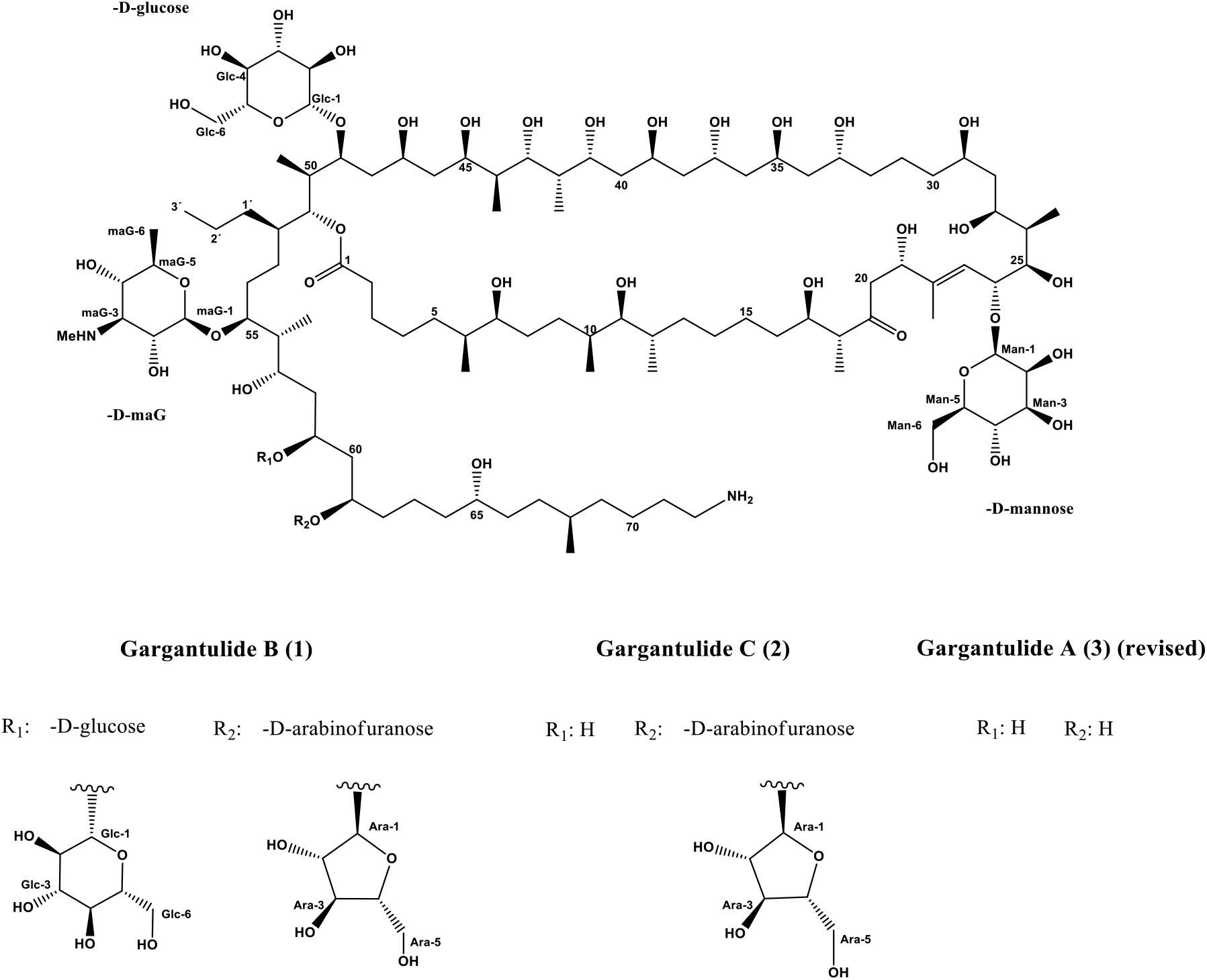
Structures of gargantulide B (**1**) and C (**2**) isolated from culture broths of *Amycolatopsis sp*. CA-230715 and that of gargantulide A (revised in this work)

Integration of genomic data from bioinformatics analysis of the T1PKS enzymes and detailed NMR-based analysis allowed us to confidently assign the stereochemistry of the polyketide chain of gargantulides B and C and to further revise and complete that of gargantulide A. Additionally, we determined the absolute configuration of the additional monosaccharide residues in gargantulides B and C, thus completing the full stereostructure of these giant macrolides (Fig. 1).

## Results and Discussion

### Isolation and planar structure of gargantulides B and C

An acetone extract of a fermentation broth of the strain *Amycolatopsis sp*. CA-230715 showed selective activity against *Acinetobacter baumannii* MB5973. Bioactivity-guided fractionation (*SI Appendix*, Fig. S1) and subsequent HRESIMS analyses of the active fractions traced this activity to gargantulides B (**1**) and C (**2**), as two peaks with [M+H]^+^ ions at *m/z* 2392.4804 and 2230.4286 (*SI Appendix*, Figs. S2, S3), which were indicative of the previously unreported molecular formulae C_116_H_218_N_2_O_47_ (Δ -0.13 ppm) and C_110_H_208_N_2_O_42_ (Δ +0.22 ppm) respectively. These formulae assignments were further supported by the presence of [M+2H]^2+^ and [M+3H]^3+^ ions for each compound (*SI Appendix*, Figs. S2, S3) and advanced the close relationship between them. Searches in natural products databases failed to identify known compounds with the observed exact masses for **1** and **2**, suggesting that they were new secondary metabolites. The structures of gargantulides B (**1**) and C (**2**) were therefore fully elucidated by combining extensive NMR spectroscopy, genome-based bioinformatics and chemical approaches. The planar structure of **1** was determined by 1D (^1^H and ^13^C) and 2D NMR (COSY, TOCSY, HSQC, HSQC-TOCSY and HMBC) (*SI Appendix*, Table S1, *SI Appendix*, Fig. S4). Detailed analyses of the ^13^C NMR and HSQC spectra revealed the presence of 3 quaternary carbons (including one ketone at δ_C_ 214.6 and one ester at δ_C_ 175.2), 1 olefinic methine (δ_C_ 123.1), 46 oxygenated methines (including five anomeric carbons at δ_C_ 108.2, 104.4, 103.9, 102.4 and 97.0 ppm, representing five sugar moieties) 12 aliphatic methines, 40 methylenes (one of them being attached to an amino group, at δ_C_ 41.0) and 14 methyl carbons (including a *N*HMe group at δ_C_ 31.1). From 9 double-bond equivalents deduced from the molecular formula, one was assigned to an olefinic double bond and two to carbonyl moieties, suggesting that **1** contained six rings in its structure. All these data jointly advanced the glycosylated polyketide macrolactone nature of **1**. Furthermore, many of these NMR features and the high molecular weight of **1** resembled those of the previously reported macrolide gargantulide A (**3**) (16). Indeed, a comparison of the HSQC and ^13^C NMR spectra of **1** (*SI Appendix*, Fig. S4) with those reported for **3** clearly evidenced the close relationship between both compounds. From the analysis of COSY and TOCSY spectra (*SI Appendix*, Fig. S4) three spin systems comprising C-2 to C-18, C-20 to C-21 and C-23 to C-72, could be constructed (Fig. 2). For the assignment of contiguous methylenes within each segment, we relied on key HSQC-TOCSY and HMBC correlations (*SI Appendix*, Fig. S4). Further ^1^H–^13^C-HMBC correlations observed from methylene protons H-2 and H-3 to carbonyl C-1 (δ_C_ 175.2) and from H-17/H-18 and H-20/H-21 to carbonyl ketone C-19 (δ_C_ 214.6), enabled the extension of the linear polyketide chain from C-1 to C-21 (Fig. 2). Additionally, intense HMBC cross-peaks from H-21 to olefinic carbons C-22/C-23 (*SI Appendix*, Fig. S4) linked the last independent spin systems and allowed the construction of the linear polyketide chain from C1 to C-72 (Fig. 2). Finally, one long-range correlation from H-51 to C-1 (*SI Appendix*, Fig. S4) alongside the downfield-shifted resonance frequency of the oxymethine proton H-51 (δ_H_ 4.88) formally assembled a 52-membered macrolide ring system (Fig. 2).

**Figure 2.**
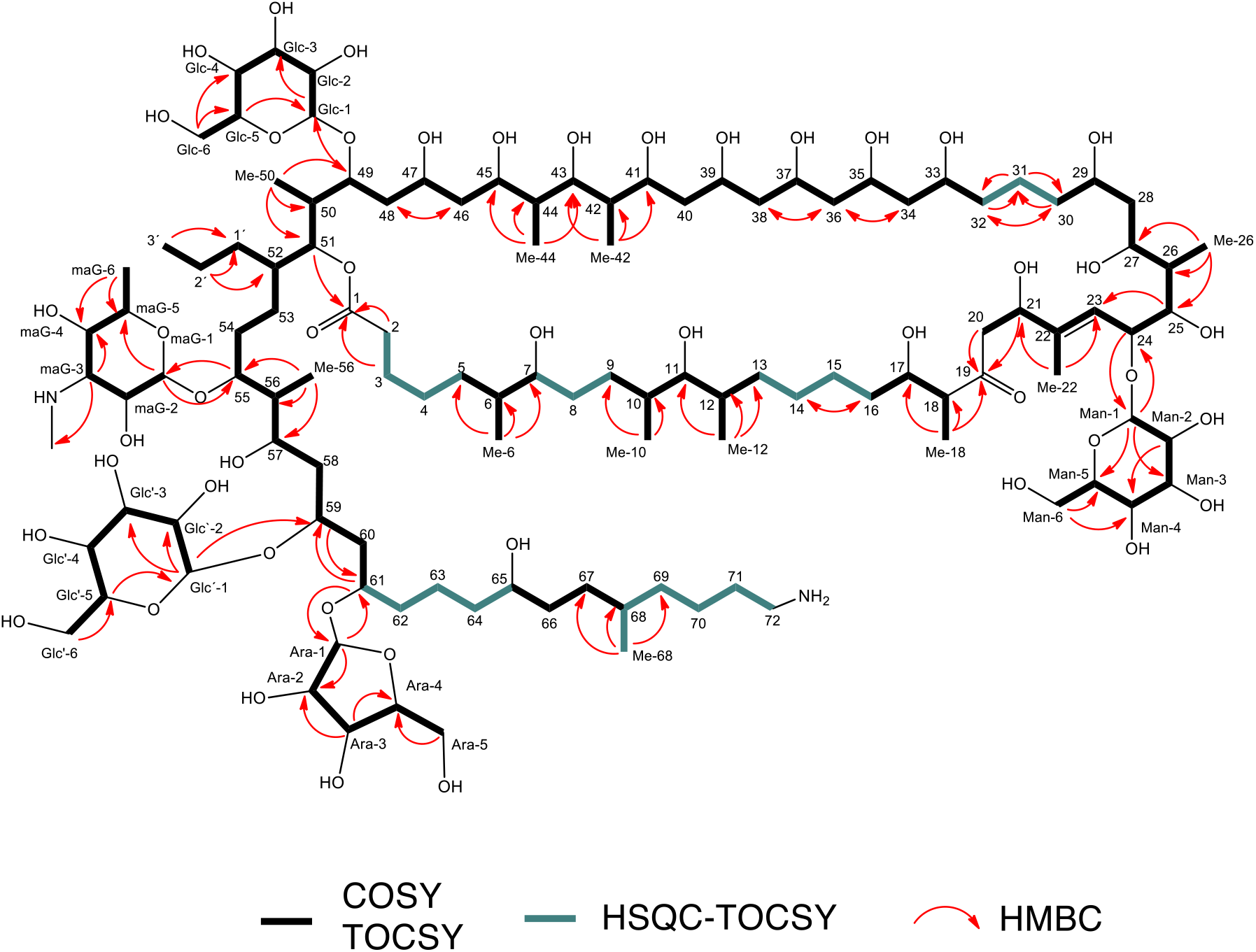
Key COSY, TOCSY, HSQC-TOCSY and HMBC correlations observed for gargantulide B (**1**)

Gargantulide B (**1**) (Fig. 1) has only one double bond between C22/C-23 whose *E* geometry was determined based on NOESY correlations between H-21/H-23 and Me-22/H-24.

The presence of five monosaccharide residues in **1** was revealed by the analysis of the remaining 2D NMR data (*SI Appendix*, Fig. S4). Starting from anomeric protons at δ_H_ 4.58, 4.20 and 4.46, three hexose units were constructed by analysis of COSY and TOCSY spectra. Based on ^3^*J*_H,H_ coupling constants and NOESY correlations, we identified the three pyranose sugar components as *β*-mannose, *β*-glucose and the rare *β*-3,6-deoxy-3-methylamino glucose (maG), already reported as components of gargantulide A. Key HMBC correlations between their anomeric protons and the corresponding positions in the backbone of **1** at C-24, C-49 and C-55, respectively, allowed us to establish the independendent *O*-glycosidic linkages of these residues at positions equivalent to those described for **3**. An additional hexose residue was found in **1** and determined to be an extra *β*-glucose unit based on the ^3^*J*_H,H_ coupling constants and *nOe* cross-peaks. HMBC correlations indicated that this sugar residue was *O*-attached to C-59 in the backbone of **1**. Finally, the remaining 2D NMR data (*SI Appendix*, Fig. S4) accounted for the presence of a pentofuranose moiety in **1**. The *α*-arabinofuranosyl configuration was assigned from ^13^C-NMR fingerprint chemical shift values (17, 18), as well as from small coupling constants ^3^*J*_1,2_ (1.6 Hz) and ^3^*J*_2,3_ (3.9 Hz) (*SI Appendix*,Table S1). Based on HMBC correlations, this arabinose unit was found to be *O*-linked to C-61 in gargantulide B (**1**).

As advanced above, gargantulide C (**2**) (Fig. 1) was assigned the molecular formula C_110_H_208_N_2_O_42_ by HRESI(+)-TOF MS ([M+H]^+^ at m/z 2230.4281, calculated for [C_110_H_209_N_2_O_42_]^+^ 2230.4274) (*SI Appendix*, Fig. S3), accounting for eight degrees of unsaturation, one less than gargantulide B (**1**). The ^1^H, ^13^C and HSQC NMR data of **2** (*SI Appendix*, Table S1, Fig. S5) showed high similarity to those of **1**, except for the absence of the signals corresponding to the *β*-glucose moiety *O*-attached to C-59. Further analysis of COSY and HSQC-TOCSY spectra confirmed the lack of this glucose residue in **2**, also reflected in the upfield-shifted resonance of C-59 compared to **1** (*SI Appendix*, Table S1) and consistent with the relationship between molecular formulae of both compounds. Connectivity of the polyketide skeleton of **2** was established based on in depth analysis of 2D NMR correlations (*SI Appendix*, Fig. S5, S6) and was identical to that of **1**. Finally, the four sugar moieties present in **2** were as well identified as *β*-mannose, *β*-glucose, 3,6-deoxy-3-methylamino glucose (maG) and *α*-arabinofuranose (analogously to **1**), and their respective glycosidic linkages positions to the backbone were found to be identical to those of **1** (C-24, C-49, C-55 and C-61, respectively).

Interestingly, although the bioactivity-guided isolation process described herein led to the isolation of gargantulides B (**1**) and C (**2**), a species with an exact mass and UV spectrum identical to those of the formerly reported congener gargantulide A was co-detected together with **1** and **2** in a parallel small-scale fermentation (*SI Appendix*, Fig. S7). Unfortunately, the low production titers for this compound prevented its isolation and confirmation as gargantulide A. Nevertheless, this finding, along with the high similarity between the three macrolides, strongly supports a common biosynthetic origin for all of them. Since no genetic or biochemical information on this class of macrolides has been reported to date, the genome of the producing strain Amycolatopsis sp. CA-230715 was fully sequenced sequenced (GenBank CP059997.1, CP059998.1) and the putative T1PKS gene cluster was identified. This allowed us to gain insight into the biosynthetic machinery of these complex macrolactones and complete the determination of their absolute configurations, as discussed below.

### Identification and in Silico Analysis of the 216 Kb Type I PKS Gene Cluster in the genome of *Amycolatopsis sp*. CA-230715

In order to identify the BGC of gargantulides B (**1**) and C (**2**) in the genome of the producing strain, we used the bacterial version of the antiSMASH software (v. 6.0.0) with the default ‘ relaxed strictness’ settings (19). In total, 45 biosynthetic regions were predicted, all situated on the circular chromosome (*SI Appendix*, Table S2). The structure of the gargantulides suggests that it is a T1PKS product. One of the predicted T1PKS regions, (region 11) (*SI Appendix*, Table S2) fits well with the size and structure of gargantulides. The cluster, further referred to as the *gar* cluster, contains 10 T1PKS genes encoding 35 modules and spanning over approximately 180 kbp. The overall size of the cluster is approximately 216 kbp, placing it among the largest uninterrupted BGCs ever discovered, and simultaneously - to the best of our knowledge - as the largest T1PKS cluster by the number of encoded modules. The putative functions of all genes in this cluster were determined by sequence comparisons and are listed in *SI Appendix*, Table S3 and depicted in Figure 3.

**Figure 3.**
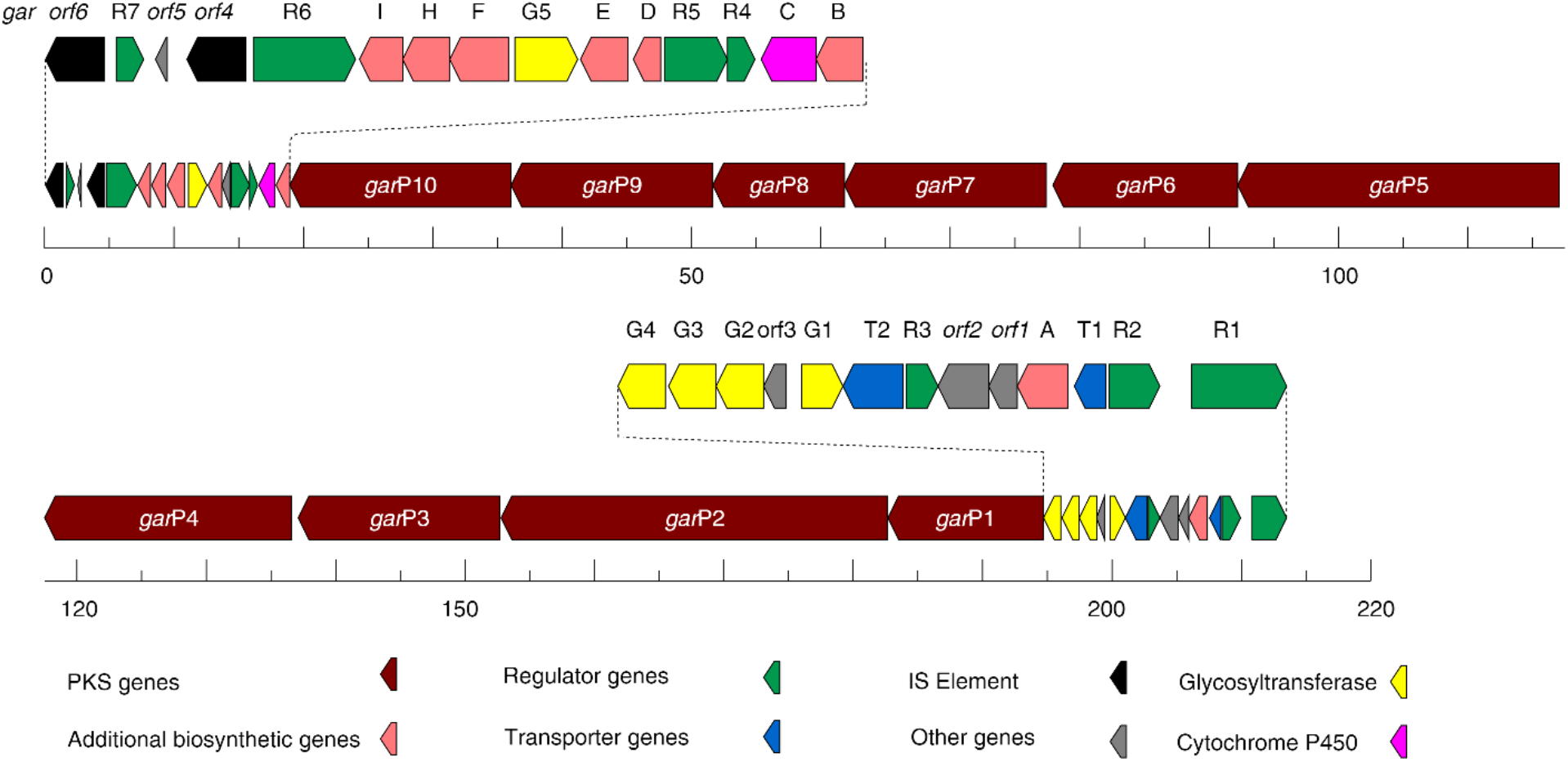
Scheme of the *gar* biosynthetic gene cluster with putative functions of all genes (see table S3).

To confirm that gargantulides are indeed derived from the *gar* cluster, we attempted to genetically engineer the strain. However, initial tests indicated intrinsic resistance to all commonly used antibiotics for selection in Actinobacteria genetics. This unfortunately at the moment prevents direct genetic engineering of the strain. To identify other bacteria that could contain *gar*-like BGCs, we used Cluster BLAST and BIG-SCAPE to search a database of complete Actinobacterial genomes. However, no further strains were found that contain closely related BGCs. The genome of *Streptomyces* sp. A42983, the producer of gargantulide A (16) is not publicly available, which prevented the direct comparison. In the proximity of the putative gar BGC, we identified 2 genes, *orf4* and *orf6*, which are highly similar to IS4 and IS256 mobile genetic elements. The presence of such IS-like elements might hint at potential horizontal transfer events in this region of the genome. In the *gar* gene cluster there are 10 core genes that code for a modular T1PKS named *garP1* to *garP10* (Figure 3; *SI Appendix*, Table S3). Two putative ABC transporters *garT1* and *garT2* are situated in close proximity to the core genes and may play a role in transport and/or resistance of/against the produced macrolides. Of note, both transporters are predicted to have nucleotide-binding domains, while only one of them, GarT2, possesses the predicted 6 transmembrane helical domains (*SI Appendix*, Table S3) typically found in ABC transporters that function as exporters (20). The transporter GarT2 is situated right next to the putative TetR-family regulator GarR3. The gar BGC-containing region contains 7 putative regulators that were named *garR1* to *garR7* (*SI Appendix*, Table S3). Such a high number of regulators in the *gar* BGC indicates the need for tight and effective regulation of the expression of the gene cluster.

The structure of the gargantulides aglycon is in very good agreement with the product that can be predicted based on the PKS domain organization (*SI Appendix*, Table S4). Based on NMR data, the starter unit of GarP1 is proposed to be 4-aminobutanoyl-CoA, which might be derived from ornithine via putative oxidative deamination and decarboxylation reactions (21) and introduced as a starter unit through the function of the putative CoA-ligase encoded by *garA*.

To predict the substrates of all AT domains, activity and stereochemistry of KR, DH and ER domains, sequence alignments of the individual domains were performed and key residues involved in substrate selection or stereochemistry of the reaction products were extracted and compared with references (22–25) (*SI Appendix*, Table S4 and Fig. S8). In case of AT domains, all but one (module 9), have amino acid signatures indicating malonyl-CoA (modules 2-6, 8, 11, 12, 15-21, 23, 25, 27, 28, 31, 33, 34) or methylmalonyl-CoA (modules 1, 7, 10, 13, 14, 22, 24, 26, 29, 30, 32) specificity, which are common extender units for the biosynthesis of polyketides (25). AT domain in module 9 is clearly different and is proposed to be responsible for the introduction of the unusual branched extender unit propylmalonyl-CoA (*SI Appendix*, Fig. S8-A), which was confirmed by NMR. Consistently, the *gar* BGC includes a putative crotonyl-CoA carboxylase/reductase coding gene *garF* and a putative FabH-coding gene *garH*, products of which are well known to be involved in the biosynthesis of unusual branched extender units (25). The cluster contains one inactive KR domain (type C1) at module 26, which is consistent with the presence of a keto function instead of a hydroxy group at C-19 in the structures. The remaining KR domains belong to the A1, B1 and B2 types (*SI Appendix*, Table S4). NMR data supports the presence of a single double bond between C-22/C-23, introduced by the DH domain at module 24, suggesting that the DH domains at modules 8 and 11 are inactive. Close examination of the DH domain at module 11 shows the absence of a key catalytic histidine and aspartic acid (*SI Appendix*, Fig. S8-D). In the case of the DH domain at module 8, although the key residues in the catalytic domain are conserved, several supporting active site residues are missing (*SI Appendix*, Fig. S8-D). Taking these findings together and assuming the collinearity rule, we can propose a biosynthetic pathway for the common backbone of gargantulides A-C (Fig. 4).

**Figure 4.**
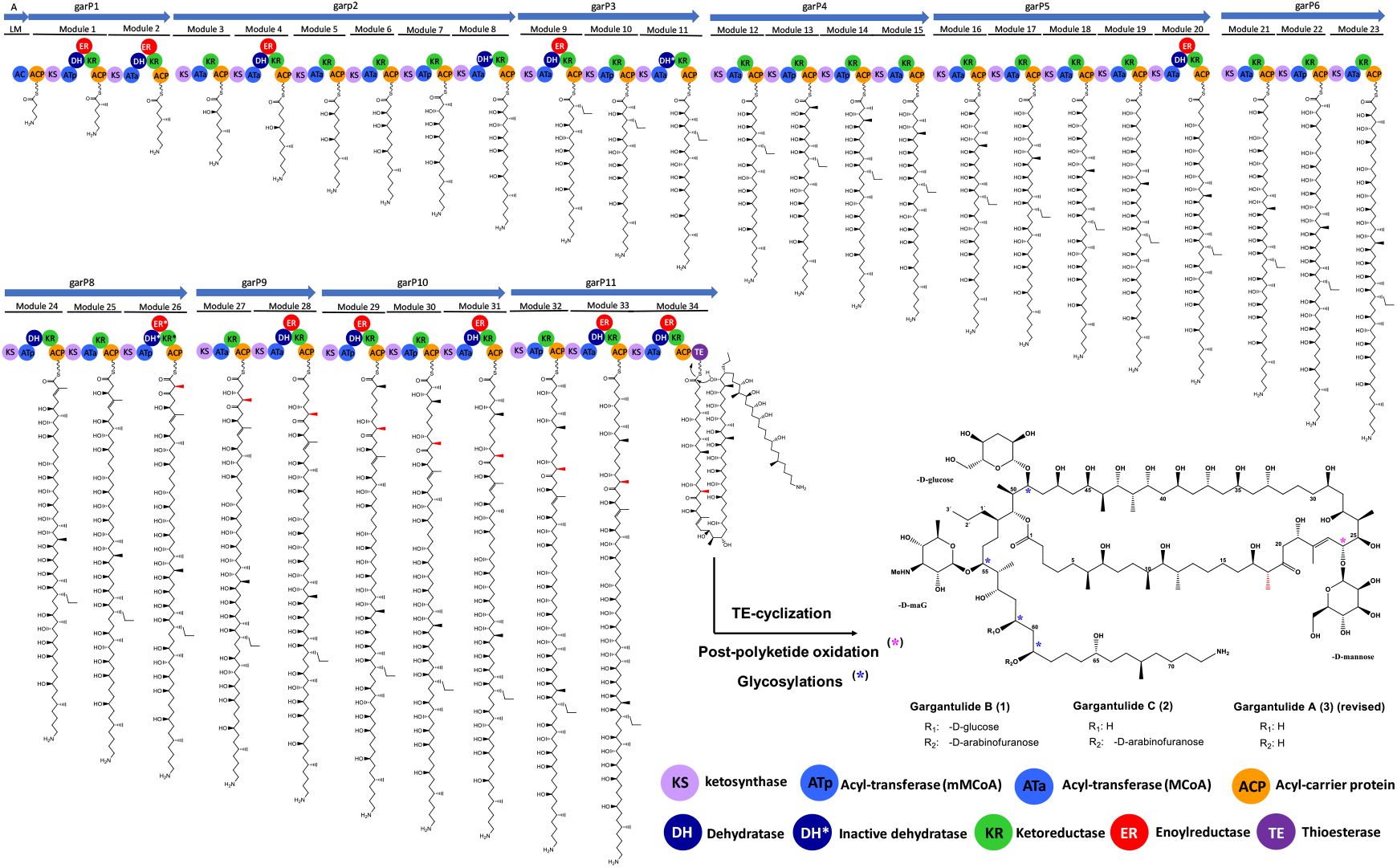
Proposed biosynthesis of the polyketide core of gargantulides A-C. Asterisks indicate post-PKS modifications. Methyl-epimerization at C-18 (redox-inactive KR domains in module 26) is marked in red.

After the polyketide chain is released from the PKS complex and the TE-domain mediated cyclization, hydroxylation at C-24 and several different glycosylations take place. Despite the high degree of glycosylation of gargantulides A-C, only few genes encoding for proteins putatively involved in sugar biosynthesis are found in the *gar* BGC. All three macrolides contain the unusual amino sugar 3,6-deoxy-3-methylamino glucose (maG) attached at C-55, which, to the best of our knowledge, is present exclusively in this family of compounds. We hypothesized that this amino sugar could share a common biosynthetic origin with mycaminose (3,6-deoxy-3-dimethylamino-D-glucose), an amino sugar found in other glycosylated macrolides (e.g. tylosin), whose biosynthesis has been extensively studied (26–28). Within the *gar* BGC, *garD* codes for a protein with high level of similarity (51% identity, 64% similarity) to Tyl1a from the mycaminose biosynthetic pathway in the tylosin BGC (29), suggesting it could function as a TDP-4-keto-6-deoxy-D-glucose 3,4-isomerase. *garE* encodes for a PLP-dependent aminotransferase with high homology (62% identity, 73% similarity) to the biochemically characterized aminotransferase TylB from mycaminose biosynthesis (30). Thus, GarE is proposed to catalyze a C-3 transamination to afford TDP-3-amino-3,6-dideoxy-D-glucose. Surprisingly, the *gar* cluster lacks a gene encoding for a putative *N*-methyltransferase to convert TDP-3-amino-3,6-dideoxy-D-glucose into *N*-demethyl-mycaminose (i.e., maG). To look for a candidate gene responsible for the *N*-methylation, we employed the amino acid sequences of some known methyltransferases found in the mycaminose, desosamine, L-ossamine and D-forosamine pathways from different microorganisms. The best candidate was HUW46_03188, located just outside of the boundaries of the BGC 15, which encodes for a putative S-adenosylmethionine-dependent methyltransferase with high similarity (51% / 65% amino acid identity / similarity) to DesVI, the biochemically characterized *N,N*-dimehyltransferase from desosamine biosynthesis (31) (*SI Appendix*, Table S5). Interestingly, other DesVI homologous proteins, such as TylM1 (tylosin gene cluster) or SpnS (spinosyns BGC), have been shown to catalyze *N*-methylation in a stepwise manner, involving the release of the monomethylated product from the active site (31, 32).This could explain the formation of the monomethylated amino sugar maG in gargantulides A-C. Additionally, genes encoding for a putative glucose-1-phosphate thymidyltransferase and dTDP-glucose 4,6-dehydratase, required in the first two steps to afford the intermediate TDP-4-keto-2,6-dideoxy-D-glucose, are also found elsewhere in the genome (*SI Appendix*, Table S6). Based on these findings, a biosynthetic pathway to maG from glucose-1-phosphate can be proposed (*SI Appendix*, Fig. S9).

Five genes encoding for glycosyltransferases, *garG1*-*garG5*, are found in the *gar* BGC and assumed to be responsible for the introduction of the different monosaccharide residues. GarG1-GarG4 are glycosyltransferases of the GT-1 inverting family, whereas GarG5 belongs to the GT-39 inverting family (33). Unlike described for the mycaminose attachment to the tylosin aglycon (34), no glycosyltransferase requiring a P450-like activator is found in the *gar* BGC. Moreover, GarC, the only P450 enzyme encoded in the BGC, contains the conserved cysteine within the Cys-pocket found in canonical P450 proteins, which is reported to be missing in the P450-like activators of these glycosyltransferases (35). Thus, GarC is likely to function as a canonical P450 enzyme catalyzing hydroxylation at position C-24.

### Correlation of bioinformatics assignments with NMR-based analysis. Proposal of absolute stereochemistry for gargantulides A-C

Gargantulides A (**3**), B (**1**) and C (**2**) contain 34 stereocenters in the shared common polyketide backbone, along with other 15, 24 or 19 additional chiral centers respectively, from their monosaccharide residues. Despite the complexity of these huge and stereochemically rich polyol macrolides, the relative configurations for most of the chiral carbons within the polyketide chain of gargantulide A were originally established by a combination of NOESY NMR, *J*-based configuration analysis and Kishi’s universal NMR database method (16). Absolute configurations for these centers were also proposed by using the stereochemically defined sugar residues as internal probes, employing the corresponding *nOe* correlations between the monosaccharide units and the aglycon (16). However, the configurations of eight stereocenters (C-6, C-7, C-10, C-11, C-12, C-17, C-18 and C-68) remained unassigned.

The likely common biosynthetic origin of gargantulides A-C (**1**-**3**) ensures the same absolute configuration for the three congeners. Therefore, in order to establish or complete the stereochemical assignments of gargantulides A-C, the bioinformatic-derived assignments for each stereocenter within the polyketide backbone (except that of C-24) were compared and validated with those obtained from NMR data.

Unexpectedly, some discrepancies were found between the *in silico* assigned stereochemistry for gargantulides B and C and that determined experimentally for gargantulide A (16). In terms of relative configuration, the NMR-based determinations for gargantulide A are in excellent agreement with the bioinformatics prediction for all the stereocenters reported, except for C-55. Strikingly, bioinformatics analysis of the large central segment comprising C-25 to C-52, predicted the same relative stereochemistry but opposite absolute configuration to that reported for gargantulide A. On the other hand, configurations of stereocenters within the segment C-6 to C-18, as well as that of C-68, were not assigned for gargantulide A but they have now been predicted *in silico* (*SI Appendix*, Fig. S10). To make a definitive proposal of the absolute stereochemistry for gargantulides A-C, we then set out to clarify disagreements between the *in silico* predictions and the experimental determinations for gargantulide A, as well as to experimentally verify the predicted absolute configurations for those stereocenters not previously determined. For this purpose, we employed a combination of Kishi’s universal NMR database (36, 37), ^1^H-^1^H coupling constant and NOE analyses and qualitative ^3^*J*_C,H_-based configuration analysis (38–40).

Focusing on **1**, we first confirmed that the relative stereochemistry of gargantulide B (and thus also that of C) was consistent with the prediction and that it was essentially the same as that partially reported for gargantulide A. Of note, a single comparison of the nearly identical ^1^H and ^13^C chemical shift values (except for the differentially substituted positions) as well as the ^1^H-^1^H coupling constants (*SI Appendix*, Table S1) for gargantulides A, B and C, provided a clear indication that we would obtain the same results. Thus, similar to what was reported previously (16), relative configurations within segments C-27 to C-49 and C-57–C61 were assigned by applying the Kishi’s NMR databases corresponding to the 1,3-diol (C-27–C-29), 1,3,5,7-pentol (C-33–C-41), 1,3,5-triol (C-45–C-49 and C-57–C61) and the hydroxy/methyl/hydroxy/methyl (C-41–C-45) sets (39, 41– 43). Additionally, the Matsunaga empirical rule for 1,5-diols (44) allowed us to establish *anti* and *syn* configurations for the pairs C-29/C-33 and C-61/C-65, respectively. In parallel, relative configurations within C-24–C-27 and C-49-C-52 segments were also confirmed to be consistent with those reported for gargantulide A based on NOESY, ^3^*J*_H,H_ and ^3^*J*_C,H_-based configuration analyses.

The absolute configuration of C-55 in gargantulide A was defined as *R*, and the C-55/C-56/C-57 substereocluster was assigned a *syn/syn* relative configuration (16). However, the *in silico* analysis predicted a 55*S* configuration and the *anti/syn* relationship for those centers (*SI Appendix*, Table S4 and Fig. S10). From the NMR-based analysis, we were able to validate this bioinformatics assignment based on the following observations: First, even in spite of the glycosylation at C-55, the application of the Kishi’s data set I (37) (*SI Appendix*, Fig. S11-a) to Me-56 (δ_C_ 10.7) strongly suggested an *anti/syn* configuration for C-55/C-56/C-57, which was further supported by NOESY correlations between Me-56 and H-55, H-54 (one of the methylene protons at δ_H_ 1.89) and H-58 (*SI Appendix*, Figs S11-c and d). Second, although ^3^*J*_H,H_ constants could not be accurately measured for H-55/H-56/H-57, the higher multiplicity of H-55 compared to that of H-57 as well as the appearance of H-56 as a “dqd” with only one large coupling constant (^3^*J*_H-55,H-56_ = ca. 9 Hz; ^3^*J*_H-56,Me-56_ = 6.9 Hz; ^3^*J*_H-56,H-57_ = ca. 2 Hz), jointly suggested the *anti/syn* configuration as well (*SI Appendix*, Fig. S11-b). Finally, the small estimated values (based on the low intensity of their HMBC cross-peaks) for the coupling constants ^3^*J*_Me-56–H55_, ^3^*J*_C-54–H56_, ^3^*J*_C-58–H56_ and ^3^*J*_C-55–H57_ and the large one for ^3^*J*_Me-56–H57_, assigned Me-56/H-55, C-54/H-56 and Me-56/57-OH as *gauche* and Me-56/H-57 as *anti* (*SI Appendix*, Fig. S11-c and d), definitely consistent with the bioinformatics assignment.

In view of these results, we might conclude that disagreements between the absolute configurations determined within the segment C25-C52 and for the center C-55 in gargantulide A (**3**) and those predicted respectively from the gene cluster analysis, may have arisen from misinterpretations of some *nOe* correlations between the sugar units and the polyketide backbone of **3** in the previous work (16).

We then focused on chiral centers whose stereochemistry had not previously been determined for **3**. The *syn* configuration for C-6/C-7 was deduced from the Kishi’s NMR database for the 4-methylnonan-5-ol isomers (45) (*SI Appendix*, Fig. S12), based on the chemical shift of the branched methyl Me-6 at δ_C_14.5 ppm. This configuration assignment matched that of the bioinformatics prediction.

Regarding the stereocluster C-10–C-12, although ^1^H-^1^H coupling constants of H-10 or H-12 could not be measured due to signal overlapping, the appearance of H-11 as double-doublet (dd) with one large (8.2 Hz) and one small (3.2 Hz) ^3^*J*_H,H_ coupling constant indicated either a *syn/anti* or *anti/syn* configuration for C10/C-11/C-12. The clearly different chemical shifts for the methyl carbons Me-10 (δ_C_ 13.4 ppm) and Me-12 (δ_C_ 16.7 ppm) (*SI Appendix*, Table S1) also referred to one of these two configurations. The greater intensity of the COSY cross-peak for H-11/H-12 compared to that of H-10/H-11 (^3^*J*_H10-H11_ < ^3^*J*_H11-H12_) definitely suggested a *syn*/*anti* configuration (*SI Appendix*, Fig. S13-a), which was further supported by NOESY correlations between H-10/ H-11 and H-11/ H-9 (*SI Appendix*, Fig. S13-c). The large (estimated) value for the heteronuclear coupling constant ^3^*J*_Me-10–H11_ while the small one for ^3^*J*_Me-12–H11_ were consistent with this configuration assignment (*SI Appendix*, Figs. S13-b and d), which in turn, was in agreement with the *in silico* analysis.

Moving to the C-17/C-18 stereocluster, the absolute configuration 17*R*, 18*S* (*syn* relative configuration) was predicted from the gene cluster analysis (*SI Appendix*, Fig. S10 and Table S4). However, the NMR analysis revealed a large coupling constant ^3^*J*_H17-H18_ (8.6 Hz) (*SI Appendix*, Table S1 and Fig. S14). Furthermore, key NOESY correlations Me-18/H-17 and Me-18/H-16 were also consistent with an *anti* relative configuration (*SI Appendix*, Fig. S14). Although this result apparently did not match the bioinformatics prediction, redox-inactive KR domains such as the one in the corresponding module 26 (which ultimately gives rise to a keto function instead of a hydroxyl group at C-19) have been shown to catalyze α-methyl epimerizations (46–48), in agreement with the stereochemistry determined for C-18. Thus, the 17*R*, 18*R* absolute configuration was eventually assigned.

Finally, since C-68 is an isolated stereocenter, it was not possible to relate it to any other stereocluster and we assumed the 68*R* configuration deduced from the *in silico* analysis.

To complete the stereochemical assignments of gargantulides B and C, absolute configurations of the non-amino sugars present in **1** and **2** were determined according to a previously reported procedure (49) (*SI Appendix*, Supplementary methods and Fig. S15). This way, the β-glucose units (two in **1**, only one in **2**) as well as the β-mannose and α-arabinose residues were all determined as D sugars, thus matching the previous determinations for gargantulide A and being consistent with the inverting behavior of all GTs found in the *gar* BGC. Regarding the rare amino sugar maG, although its absolute configuration in gargantulides B and C could not be determined experimentally, we propose a D configuration assuming a common biosynthetic origin with mycaminose, as discussed above. Absolute configuration of maG was originally proposed as L in gargantulide A based on NOESY correlations between the monosaccharide and the aglycon (16). However, it should be noted that maG is *O*-glycosidically bound to C-55 in all three macrolides and that this stereocenter has been assigned in this work as the opposite absolute configuration to that originally reported for gargantulide A (**3**). Therefore, the original misassignment of the C-55 stereochemistry in gargantulide A may have led to the previous proposal of maG as L sugar.

### Bioactivity of gargantulides B and C

Compounds **1** and **2** were evaluated for their antimicrobial properties against a panel of different pathogens, including Gram-positive and Gram-negative bacteria and fungi. Thus, gargantulides B (**1**) and C (**2**) showed low activity against the fungi *Aspergillus fumigatus*_ATCC46645 (MIC 32-64 μg/mL) and were inactive against *Candida albicans*_ATCC64124 (MIC > 128 μg/mL). In contrast, and similar to gargantulide A, **1** and **2** showed potent activity against the Gram-positive bacteria methicillin-resistant and susceptible *Staphylococcus aureus* (MRSA, MSSA) and vancomycin-resistant *Enterococcus* (VRE), with MIC values in the range from 1 to 8 μg/mL. Remarkably and consistent with the bioassay-guided isolation of gargantulides B and C, both compounds showed moderate and selective activity against a clinical isolate of *Acinetobacter baumannii* (MIC values of 16-32 μg/mL) although no activity against any other Gram-negative bacteria tested (*SI Appendix*, Table S9). To our knowledge, the activity of large macrolactones against Gram-negative bacteria such as *A. baumannii* has been rarely reported, indicating that gargantulides may represent and alternative in the fight against the increasing spread of this clinically relevant pathogen.

## Conclusions

Two new glycosylated 52-membered macrolactones with 22-membered linear side-chains, gargantulides B (**1**) and C (**2**), were isolated and structurally characterized from the *Amycolatopsis sp*. CA-230715 strain. The extraordinarily large 216 kbp biosynthetic gene cluster (*gar*) encoding a 35 modules T1PKS was identified and characterized, allowing us to establish the first genetic evidence for the biosynthesis of this interesting class of compounds. Additionally, the *gar* gene cluster has been shown to harbor a set of genes putatively responsible for the biosynthesis of the rare sugar 3,6-deoxy-3-methyl amino glucose (maG), which, as far as we know, is present exclusively in this family of macrolides. The combination of NMR spectroscopy and bioinformatic analysis of the gene cluster allowed the assignments of the absolute configuration for all chiral centers in gargantulides B (**1**) and C (**2**) and the revision of those previously proposed for gargantulide A. The new macrolactones were evaluated for their antibiotic properties, showing remarkable antibacterial bioactivities against Gram-positive bacteria, including MRSA and VRE, and selective activity against the Gram-negative pathogen *A. baumannii*. This work is yet another example of the importance of the bioactivity-guided screen in the discovery of new bioactive secondary metabolites and further highlights the potential and reliability of the combination of NMR and bioinformatic analyses for the structural elucidation of complex natural products.

## Materials and Methods

All materials and methods in this study are detailed in SI Appendix, Materials and Methods.

## Acknowledgments

The work of the authors is funded by grants of the Novo Nordisk Foundation, Denmark [NNF20CC0035580, NNF16OC0021746]. The published results are part of the doctoral thesis of D.C.-M. (Doctoral Programme in Pharmacy (B15.56.1), Doctoral School in Health Sciences, University of Granada, 52005 Granada, Spain). The authors would like to thank Omkar S. Mohite for the help with BIG-SCAPE analysis.

