## Supporting Information for "Identification of the 216kbp gene cluster and structure elucidation of gargantulides B and C, new complex 52-membered macrolides from *Amycolatopsis* sp"

[a] Fundación MEDINA, Centro de Excelencia en Investigación de Medicamentos Innovadores en Andalucía, Avda. del Conocimiento 34, 18016 Armilla (Granada), Spain

[b]The Novo Nordisk Foundation Center for Biosustainability, Technical University of Denmark, Kemitorvet building 220, 2800 Kgs. Lyngby, Denmark

**Supplementary methods**

General chemical analysis procedures
Strain isolation, identification and fermentation conditions
Isolation of gargantulides B and C
Characterization data
Determination of the absolute configurations of the non-amino sugars present in gargantulides B and C
Multiple sequence alignments
Testing sensitivity of CA-230715 towards antibiotics
Antimicrobial and antifungal sensitivity testing

**Supplementary data:**

**Figure S1.** Bioassay-guided isolation of gargantulides B (**1**) and C (**2**)
**Figure S2.** UV (DAD) and ESI-TOF spectrum for pure gargantulide B (**1**)
**Figure S3.** UV (DAD) and ESI-TOF spectrum for pure gargantulide C (**2**)
**Table S1.** $^1$H NMR ($CD_3OD$, 500 MHz) and 13C NMR ($CD_3OD$, 125 MHz) data of gargantulides B (**1**) and C (**2**)
**Figure S4.** NMR spectra of gargantulide B (**1**)
**Figure S5.** NMR spectra of gargantulide C (**2**)
**Figure S6.** Key COSY, TOCSY, HSQC-TOCSY and HMBC correlations observed for gargantulide C (**2**)
**Figure S7.** LC-UV-HRMS chromatogram of the acetone crude extract
**Table S2.** antiSMASH results
**Table S3.** Putative functions of genes in *gar* BGC
**Table S4.** Prediction of activity and stereochemistry of AT, KR, ER and DH domains from bioinformatics analysis. Stereochemical outcomes for gargantulides B and C
**Figure S8.** Extracted amino acid sequence alignments of AT, KR, DH and ER domains
**Table S5.** Levels of identity and similarity of the putative *N*-Methyltransferase HUW46_03188 from CA-230715 with other known *N*-Methyltransferases
**Table S6.** Identified genes encoding for putative glucose-1-phosphate thymidyltransferase and dTDP-glucose 4,6-dehydratase in the genome of CA-230715

**Figure S9.** Proposed biosynthetic pathway for the unusual amino sugar 3,6-deoxy-3-methylamino glucose (maG)
**Figure S10.** Bioinformatics prediction of the absolute configurations for the gargantulides polyketide aglycon (a). Comparison with the absolute configurations previously reported for gargantulide A (b)
**Figure S11.** Determination of the relative configuration of the C-55 to C-57 stereocluster for gargantulide B (**1**)
**Figure S12.** Determination of the relative configuration of the C-6–C-7 stereocluster for gagantulide B (**1**)
**Figure S13.** Determination of the relative configuration of the C-10–C-12 stereocluster for gargantulide B (**1**)
**Figure S14.** Determination of the relative configuration of the C-17(R)–C-18(R) stereocluster for gargantulide B (**1**)
**Figure S15.** Determination of the absolute configuration of the non-amino sugars present in gargantulides B (**1**) and C (**2**)
**Table S9.** Antibacterial and antifungal activities of compounds **1** and **2**

### Supplementary methods.

**General chemical analysis procedures:** Optical rotations were measured on a Jasco P-2000 polarimeter. IR spectra were recorded with a JASCO FT/IR-4100 spectrometer equipped with a PIKE MIRacle single reflection ATR accessory. NMR spectra were recorded on a Bruker Avance III spectrometer (500 and 125 MHz for $^{1}H$ and $^{13}C$ NMR, respectively) equipped with a 1.7 mm TCI MicroCryoProbe. Chemical shifts were reported in ppm using the signals of the residual solvents as internal reference ($\delta_H$ 3.31 and $\delta_C$ 49.1 for $CD_3OD$). LC-UV-LRMS analysis were performed on an Agilent 1100 single quadrupole LC-MS system as previously described (1). ESI-TOF and MS/MS spectra were acquired using a Bruker maXis QTOF mass spectrometer coupled to an Agilent Rapid Resolution 1200 LC. The mass spectrometer was operated in positive ESI mode. The instrumental parameters were 4 kV capillary voltage, drying gas flow of 11 L $min^{-1}$ at 200 °C, and nebulizer pressure of 2.8 bar. TFA-Na cluster ions were used for mass calibration of the instrument prior to sample injection. Pre-run calibration was done by infusion with the same TFA-Na calibrant. Semi-preparative HPLC separation was performed on Gilson GX-281 322H2 with a semi-preparative reversed-phase column (Waters XBridge Phenyl, 150 x 10 mm, 5 µm). Acetone used for extraction was analytical grade. Solvents employed for isolation were all HPLC grade. Chemical reagents and standards were purchased from Sigma-Aldrich.

**Strain isolation, identification and fermentation conditions:** The strain CA-230715 was isolated from a soil sample collected at a waterlogged forest in Central African Republic. Initial similarity-based search with the 16S rDNA sequence (1318nt) against the EzBioCloud database indicated that the strain is closely related to *Amycolatopsis nigrescens* (82.0% identity, 91.0 % completeness).
A 250 mL fermentation of the producing microorganism was obtained as follows: a seed culture of the strain was obtained by inoculating one 150 x 25 mm tube containing 14 mL of seed-medium (soluble starch 20 g/L, glucose 10 g/L, NZ Amine Type E 5 g/L, meat extract 3 g/L, peptone 5 g/L, yeast extract 5 g/L, calcium carbonate 1 g/L, pH 7) with 0.7 mL of freshly thawed inoculum stock of CA-230715. The tube was incubated at 28 ºC, 70% relative humidity and 220 rpm for 5 days. The fresh inoculum thus generated was employed to inoculate two 250 mL conical flasks each containing 125 mL of APM9 medium (glucose 50 g/L, soluble starch 12 g/L, soy flour 30 g/L, CoCl2·6H2O 2 mg/L, calcium carbonate 7 g/L, pH 7). The flasks were incubated at 28 ºC, 70% relative humidity and 220 rpm during 13 days before harvesting.

**Isolation of gargantulides B and C:** The 250 mL culture broth was extracted with an equal volume of acetone under continuous shaking at 220 rpm for 1h. The mycelial debris was discarded by centrifugation at 9000 rpm and the filtered supernatant/acetone mixture (ca. 0.5L) was concentrated to 0.25 L under a nitrogen stream. The aqueous crude extract was loaded in a Sepabeads® SP207ss resin column and eluted using acetone/methanol (1:1). The resulting organic extract was chromatographed by semipreparative reverse-phase HPLC (Waters XBridge Phenyl, 150 x 10 mm, 5 µm; 3.6 mL/min, UV detection at 210 and 280 nm) with a linear gradient of $CH_3CN/H_2O$/0.1% Trifluoroacetic acid, from 15 to 28% $CH_3CN$ (0.1% TFA) over 15 min followed by an isocratic step of 28% CH3CN(0.1% trifluoroacetic acid) over 19 min , to yield gargantulides B (**1**, 2.1 mg, $t_R$ 22 min) and C (**2**, 1.7 mg, $t_R$ 24.5min) as white amorphous powders.

**Characterization data:**

*Gargantulide B* (**1**): $[\alpha]^{25}_D$ -28.3º (*c* 0.33, MeOH); IR (ATR) $cm^{-1}$: 3343, 2934, 1674, 1456, 1428, 1379, 1201, 1134, 1068, 1023; (+)-ESI-TOFMS *m/z* 2392.4804 $[M+H]^+$ (calcd. for $C_{116}H_{219}N_2O_{47}^+$, 2392.4808), 1197.2455 $[M+2H]^{2+}$ (calcd. for $C_{116}H_{220}N_2O_{47}^{2+}$, 1197.2455), 798.4995 $[M+3H]^{3+}$ (calcd. for $C_{116}H_{221}N_2O_{47}^{3+}$, 798.4994); $^{1}H$ and $^{13}C$ NMR data see *SI Appendix*, Table S1.

*Gargantulide C* (**2**): $[\alpha]^{25}_D$ -20.5º (*c* 0.33, MeOH); IR (ATR) $cm^{-1}$: 3342, 2935, 1674, 1456, 1425, 1379, 1201, 1135, 1067, 1024; (+)-ESI-TOFMS *m/z* 2230.4286 $[M+H]^+$ (calcd. for $C_{110}H_{209}N_2O_{42}^+$,

2230.4280), 1116.2170 $[M+2H]^{2+}$ (calcd. for $C_{110}H_{210}N_2O_{42}^{2+}$, 1116.2191), 744.4808 $[M+3H]^{3+}$ (calcd. for $C_{110}H_{211}N_2O_{42}^{3+}$, 744.4818); $^{1}H$ and $^{13}C$ NMR data see *SI Appendix*, Table. S1.

**Determination of the absolute configurations of the non-amino sugars present in gargantulides B and C:** Compounds **1** and **2** (400 µg each one) were separately dissolved in 0.1 mL of 1 N HCl and heated at 85 ºC for 3.5 h in a sealed vial. The crude hydrolysates were evaporated to dryness under a nitrogen stream and each residue was dissolved in 100 µL of pyridine containing 0.5 mg of L-cysteine methyl ester hydrochloride and heated at 60 ºC for 1 h. A 100 µL solution of *o*-tolyl isothiocyanate (0.5 mg) in pyridine was added to the mixture, which was heated at 60 ºC for 1 h. Standard monosaccharides (D-glucose, D-mannose and D-arabinose) were treated in the same manner with both L- and D- forms of cysteine methyl ester hydrochloride separately. The reaction mixtures (20 µL) were diluted with 50 µL of methanol and analyzed by ESI LC/MS on an Agilent 1100 single Quadrupole LC/MS. Separations were carried out on a Waters XBridge C18 column (4.6 × 150 mm, 5µm), maintained at 40 ºC. A mixture of two solvents, A (10% $CH_3CN$, 90% $H_2O$) and B (90% $CH_3CN$, 10% $H_2O$), both containing 1.3 mM trifluoroacetic acid and 1.3 mM ammonium formate, was used as the mobile phase under a linear gradient elution mode (isocratic 15% B for 25 min, 15-100% B in 0.1 min and then isocratic 100% B for 5 min) at a flow rate of 1.0 mL/min. Retention times (min) for the derivatized standard monosaccharides (with both L- and D- forms of cysteine methyl ester hydrochloride) present in **1** and **2** were: D-glucose + L-Cys/*o*-TolylNCS: 17.05, D-glucose + D-Cys/*o*-TolylNCS: 15.70, D-mannose + L-Cys/*o*-TolylNCS: 10.14, D-mannose + D-Cys/*o*-TolylNCS: 17.22, D-arabinose + L-Cys/*o*-TolylNCS: 20.41, D-arabinose + D-Cys/*o*-TolylNCS: 18.99. Retention times for the observed peaks in the HPLC trace of the L-Cys/*o*-TolylNCS derivatized hydrolysis product of **1** were: D-glucose + L-Cys/*o*-TolylNCS: 17.11, D-mannose + L-Cys/*o*-TolylNCS: 10.10, D-arabinose + L-Cys/*o*-TolylNCS: 20.48. Retention times for the observed peaks in the HPLC trace of the L-Cys/*o*-TolylNCS derivatized hydrolysis product of **2** were: D-glucose + L-Cys/*o*-TolylNCS: 17.10, D-mannose + L-Cys/*o*-TolylNCS: 10.10, D-arabinose + L-Cys/*o*-TolylNCS: 20.56.

**Multiple sequence alignments:** The sequences of AT, KR, DH and ER domains were manually extracted from the genome sequence of CA-230715 and analyzed using Geneious 9.1.8 software platform (2). The catalytic regions and regions valuable for the prediction of stereochemistry were identified as described in previous works (3–6). Multiple sequence alignments were run using MUSCLE Alignment function (7) of Geneious, set to 8 maximum iterations. As a comparison, domains manually extracted from the erythromycin cluster of *S. erythraea* NRRL2338 were used.

**Testing sensitivity of CA-230715 towards antibiotics:** The producing strain CA-230715 was grown on MS solid media plates for six days (8) until visual signs of sporulation were observed. The spore suspension was washed from the plate using 4 ml of sterile water and 100 µL of spore suspension was directly used for plating. The spore suspensions were plated with 100 µL on MS media and MS media supplemented with apramycin (50 µg/mL), hygromycin (50 µg/mL), spectinomycin (50 µg/mL), streptomycin (50 µg/mL), thiostrepton (50 mg/mL), erythromycin (50 µg/ml), the most commonly used antibiotics for selection in Actinobacteria genetics. The plates were incubated for six days at 28°C. All the plates, both MS supplemented and not supplemented with antibiotics were evenly covered with a lawn of mycelia, indicating the high resistance of CA-230715 against all tested antibiotics.

**Antimicrobial and antifungal sensitivity testing:** Compounds **1** and **2** were tested in antimicrobial assays against the growth of the gram-negative bacteria *E. coli* ATCC 25922, *A. baumannii* MB5973, *K. pneumoniae* ATCC 700603 and *P. aeruginosa* MB5919 and against the gram-positive bacteria methicillin-resistant *S. aureus* (MRSA) MB5393, methicillin-susceptible *S. aureus* (MSSA) and Van-A vancomycin-resistant *Enterococcus* (VRE) following previously described methodologies (9–11).

Compounds **1** and **2** were also evaluated for their antifungal activity and spectrum against filamentous fungi *Aspergillus fumigatus* ATCC46645 following previously described methodologies (9) and yeast *Candida albicans* ATCC64124 as follows: thawed stock inoculum suspension from cryovials was streaked onto Sabouraud dextrose agar (SDA) plates and incubated at 37ºC

overnight to obtain isolated colonies. The single colonies were inoculated into 20 ml RPMI 1640 (Sigma) in 250 ml Erlenmeyer flasks and were adjusted at 0.25-0.28 (OD612nm) and then diluted to 1:10 in order to obtain the inoculum for assay.
The variation of the growth was quantified by measuring absorbance (OD612nm) using the plate reader EnVision® Multilabel (Perkin Elmer) with two readings, one at initial time and other after that the plate was incubated at 37º C for 20 hours.
Briefly, each compound was serially diluted in DMSO with a dilution factor of 2 to provide 10 concentrations starting at 128 µg/mL for all the assays. The MIC was defined as the lowest concentration of compound that inhibited ≥ 95% of the growth of a microorganism after overnight incubation.
The Genedata Screener software (Genedata, Inc., Basel, Switzerland) was used to process and analyze the data and also to calculate the RZ' factor, which predicts the robustness of an assay (12). In all experiments performed in this work the RZ' factor obtained was between 0.76 and 0.93

**Supplementary data**.

**Figure S1.** Bioassay-guided isolation of gargantulides B (**1**) and C (**2**). Semipreparative HPLC-UV chromatogram (blue trace: 210 nm; (orange trace: 280 nm)

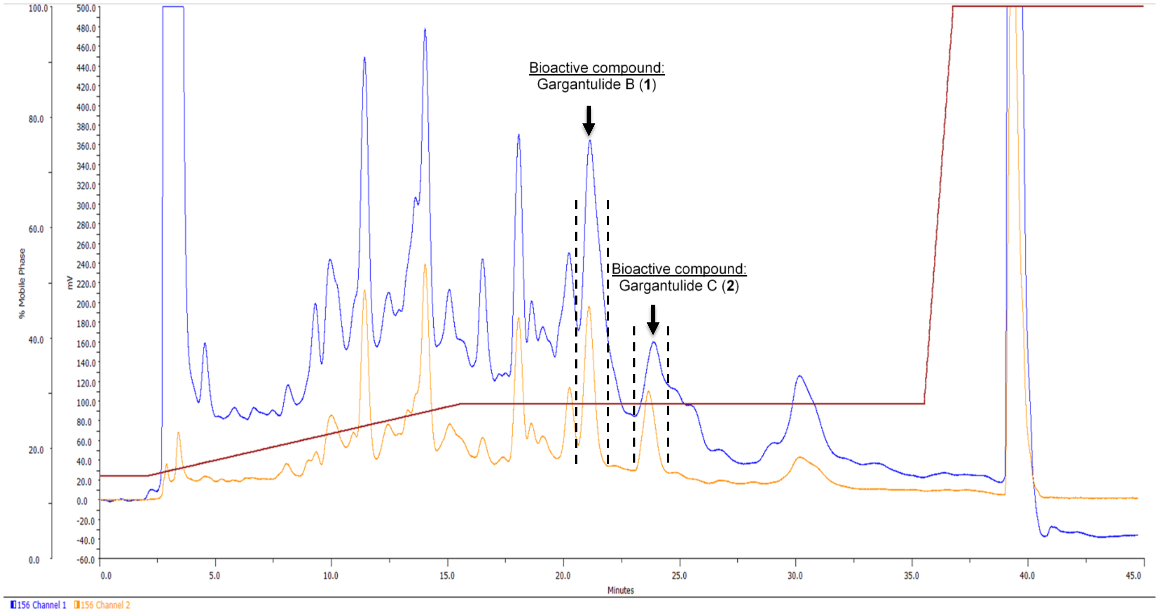

**Figure S2.** UV (DAD) and ESI(+)-TOF spectrum for pure gargantulide B (**1**)

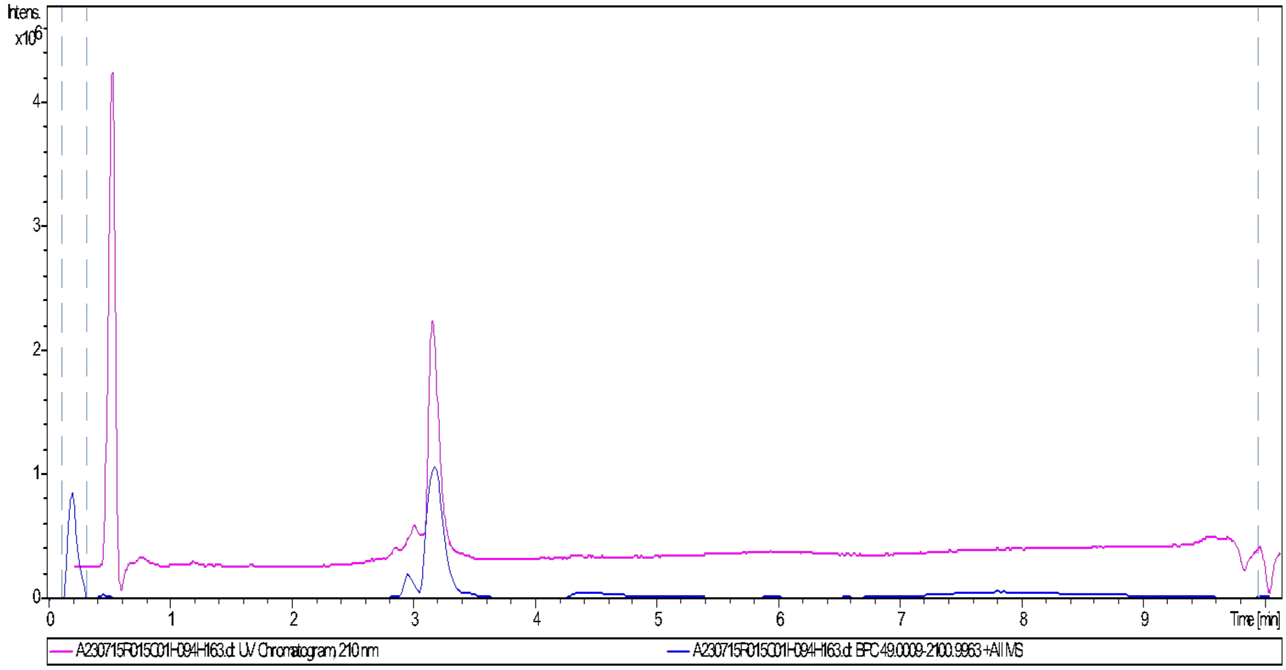


a) LC-UV-HRMS chromatogram (UV 210 nm: pink trace; MS+: blue trace) for **1**

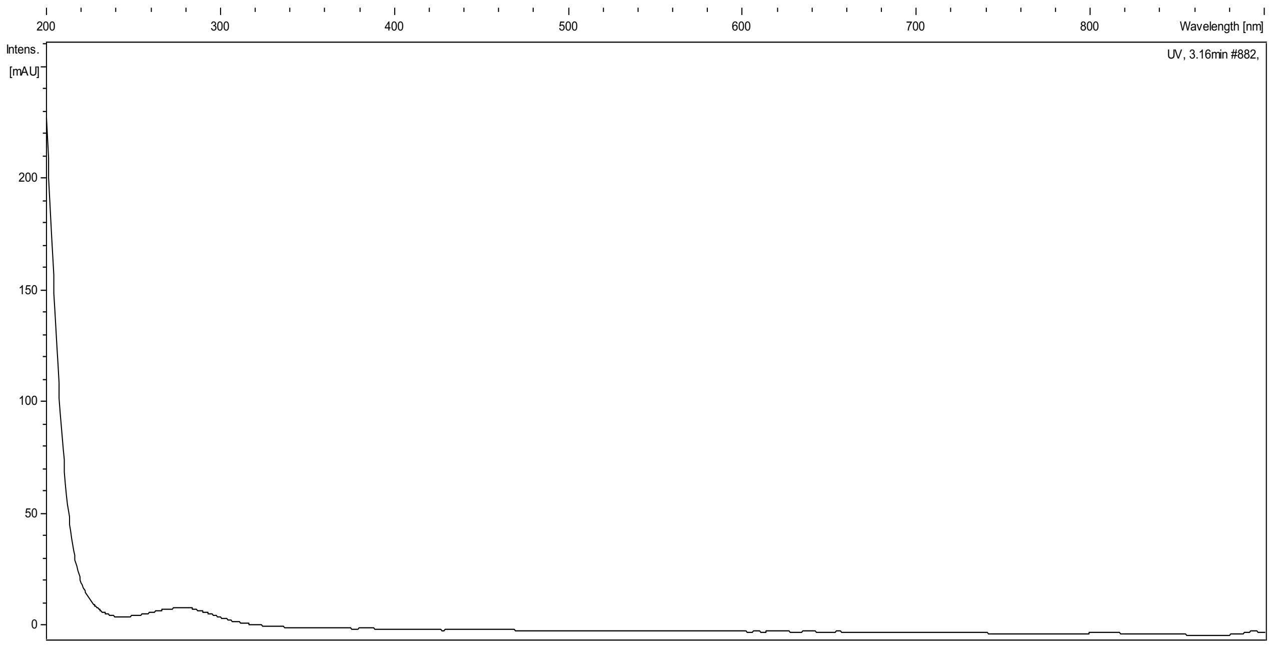


b) UV spectrum of **1**

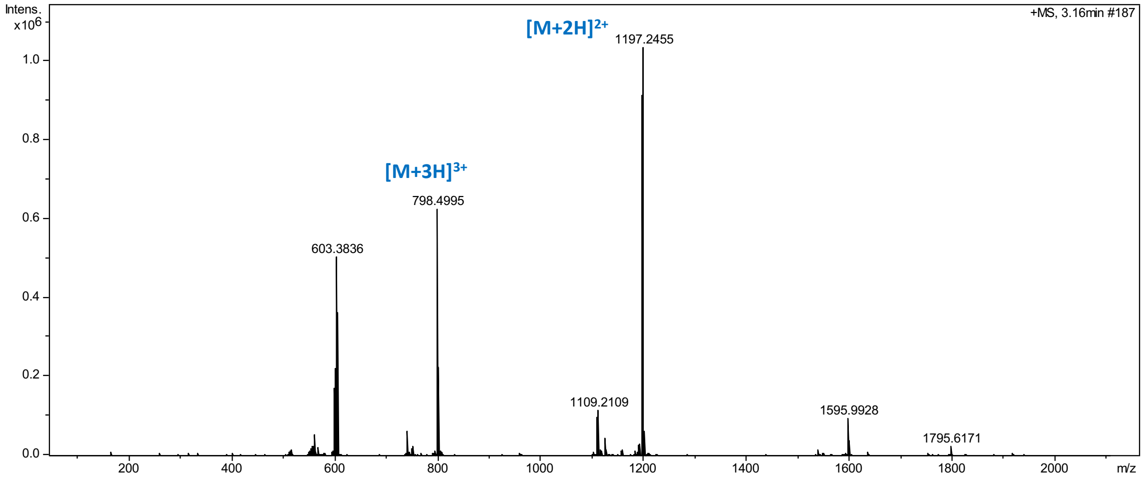


c) HRESIMS(+)-TOF spectra (ISCID 0 eV) of **1** (overview)

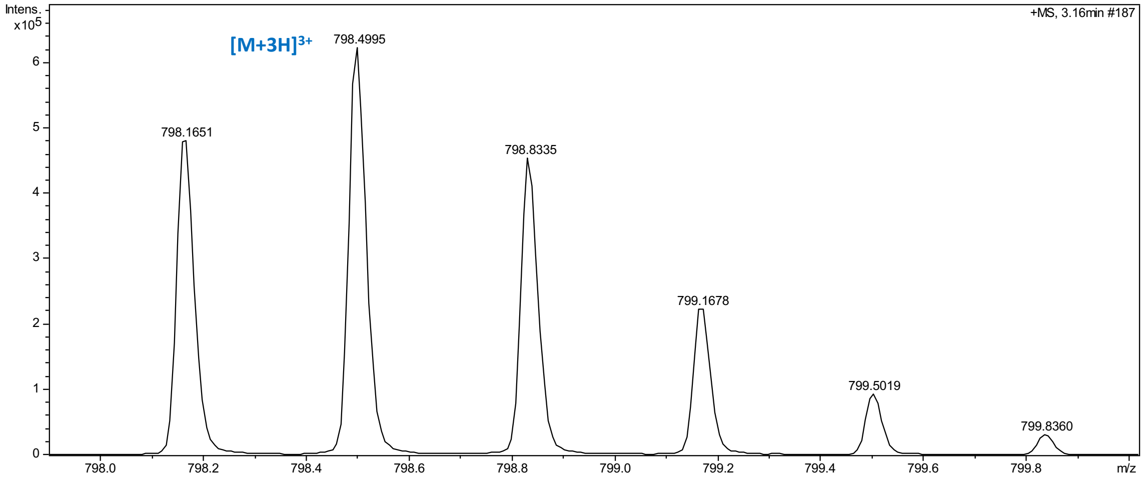


d) HRESIMS(+)-TOF spectra (ISCID 0 eV) of **1**. Triply-charged adducts (zoom in region)

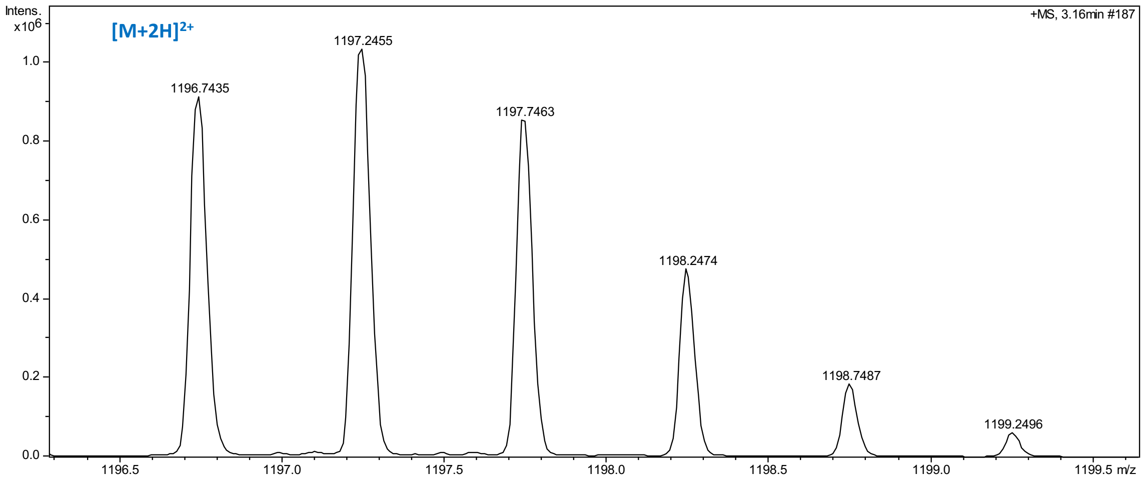


e) HRESIMS(+)-TOF spectra (ISCID 0 eV) of **1**. Doubly-charged adducts (zoom in region)

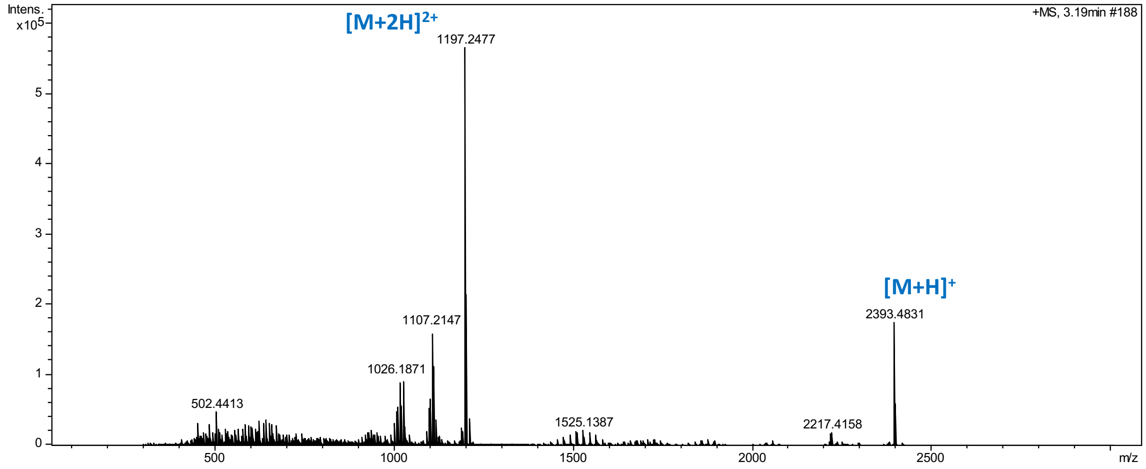


f) . HRESIMS(+)-TOF spectra (ISCID 75 eV) of **1** (overview)

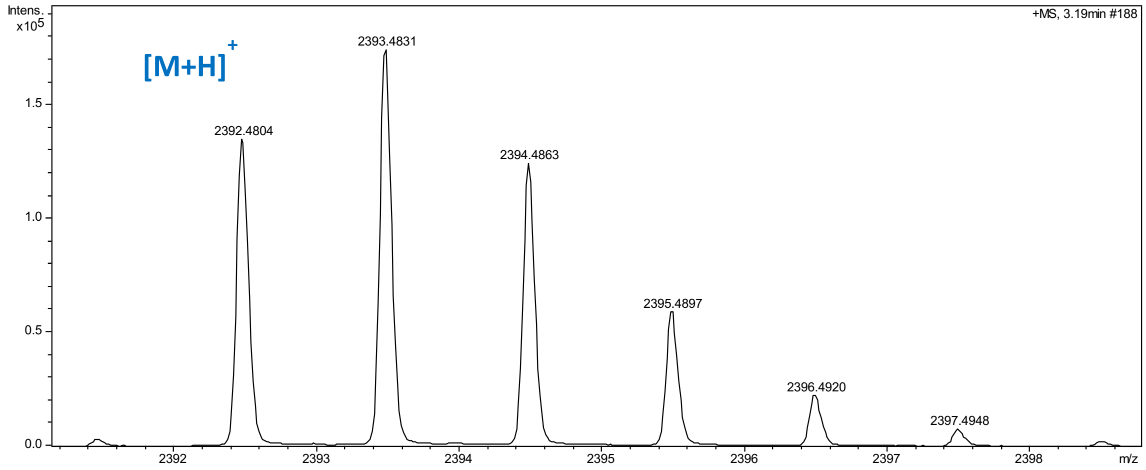


g) HRESIMS(+)-TOF spectra (ISCID 75 eV) of **1**. $[M+H]^+$ adduct (zoom in region)

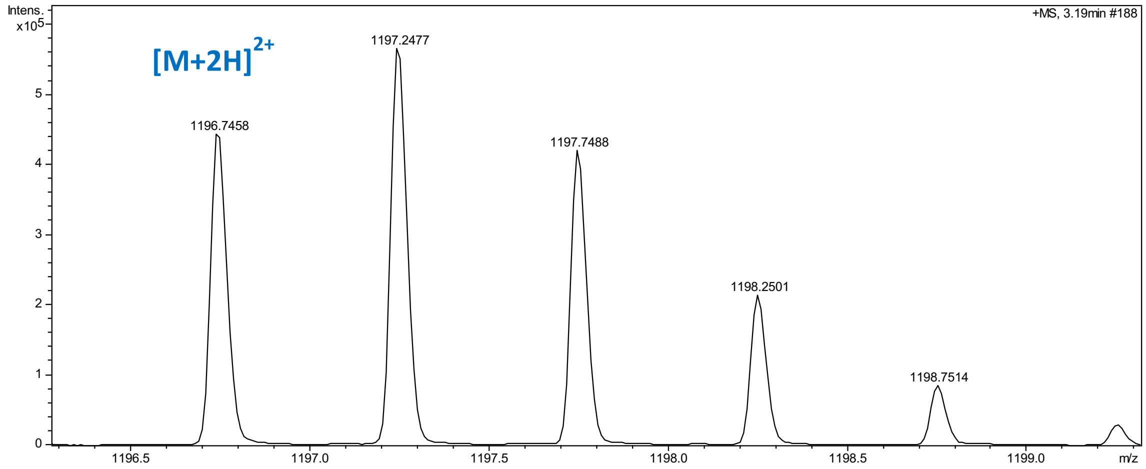


h) HRESIMS(+)-TOF spectra (ISCID 75 eV) of **1**. $[M+2H]^{2+}$ adduct (zoom in region)

**Figure S3.** UV (DAD) and ESI(+)-TOF spectrum for pure gargantulide C (**2**)

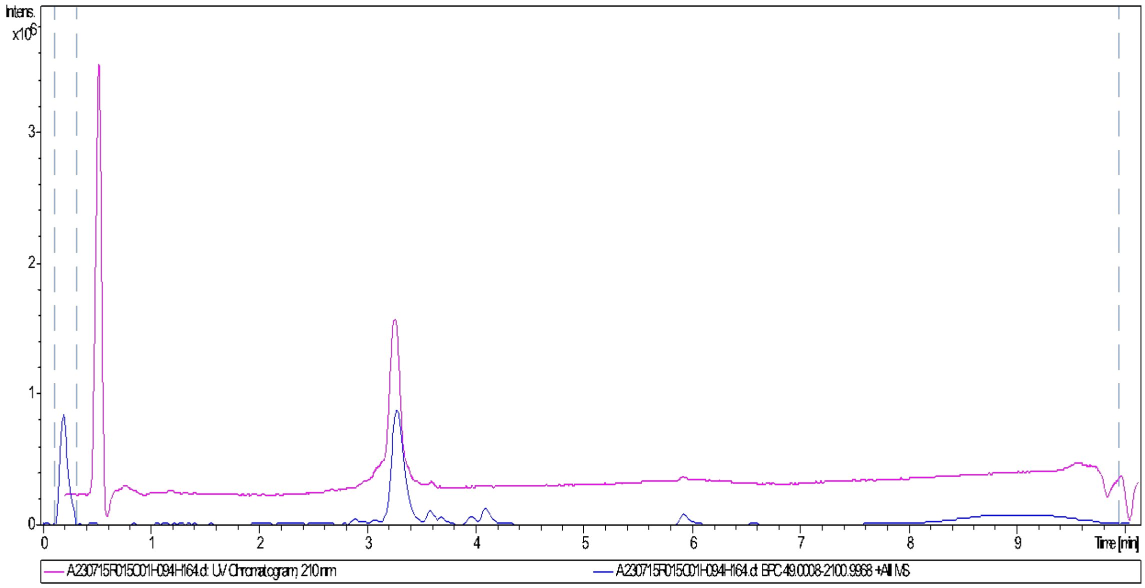


a) LC-UV-HRMS chromatogram (UV 210 nm: pink trace; MS+: blue trace) for **2**

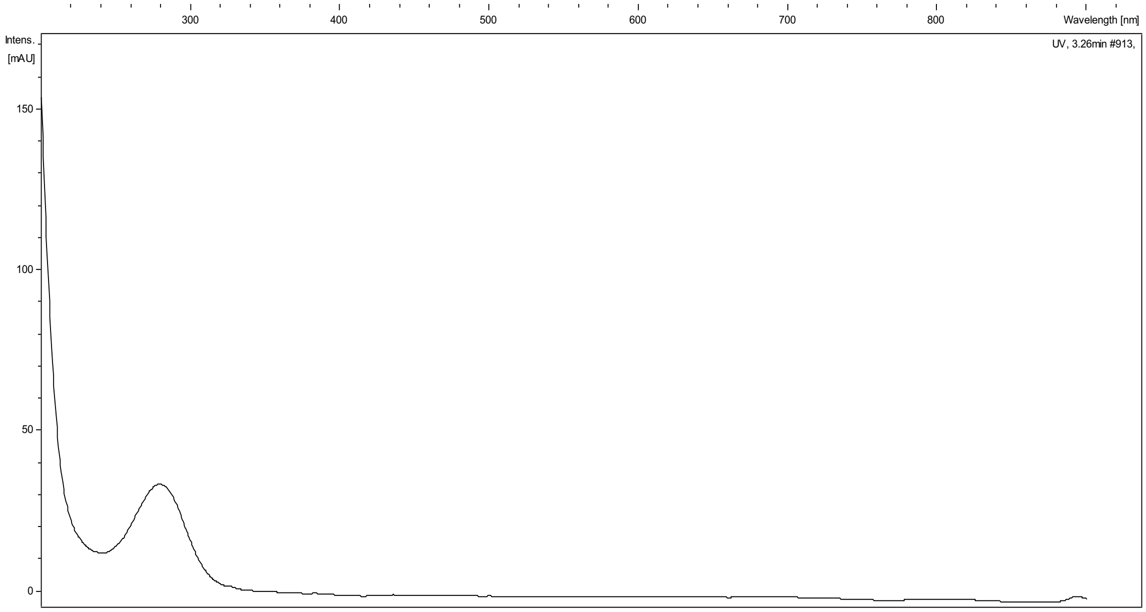


b) UV spectrum of **2**

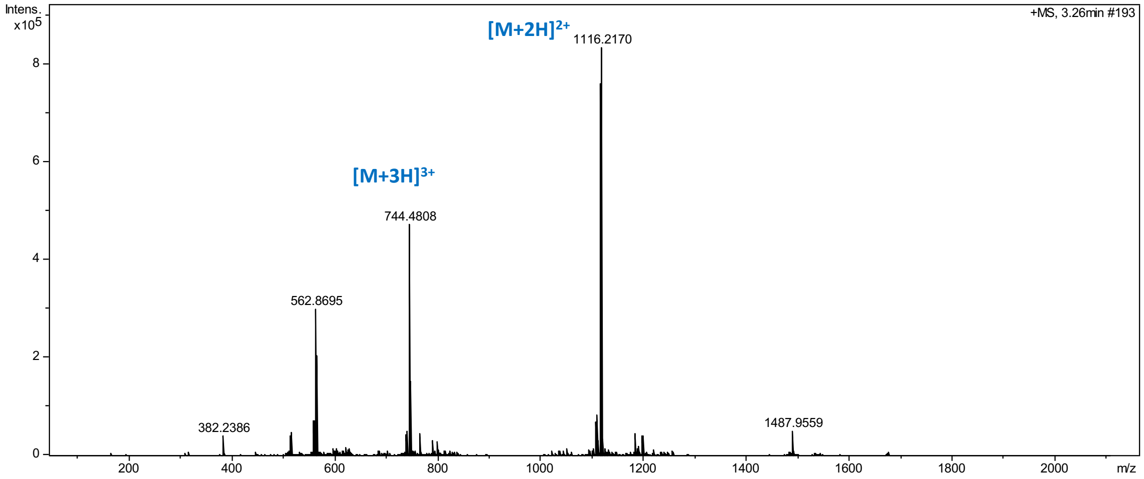


c) HRESIMS(+)-TOF spectra (ISCID 0 eV) of **2** (overview)

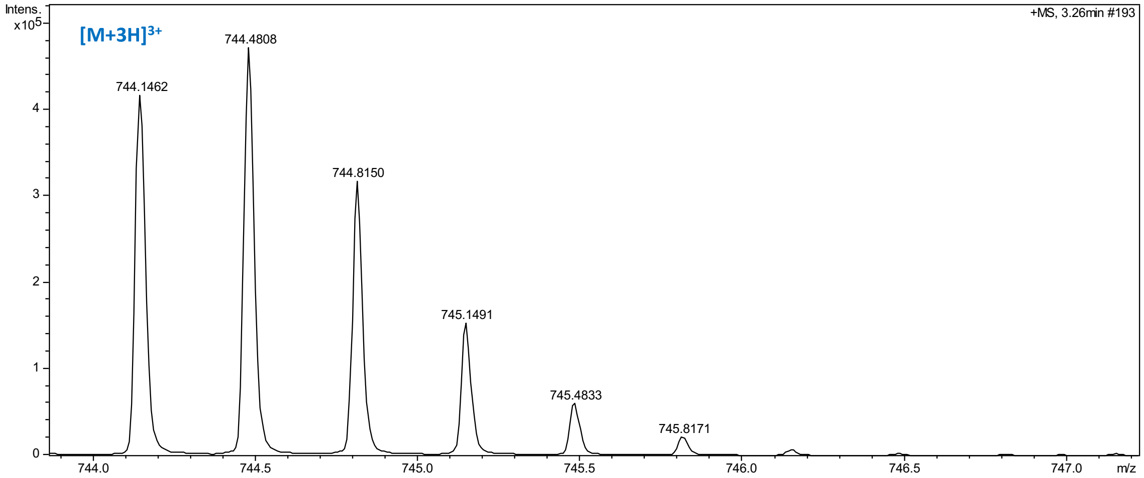


d) HRESIMS(+)-TOF spectra (ISCID 0 eV) of **1**. Triply-charged adducts (zoom in region)

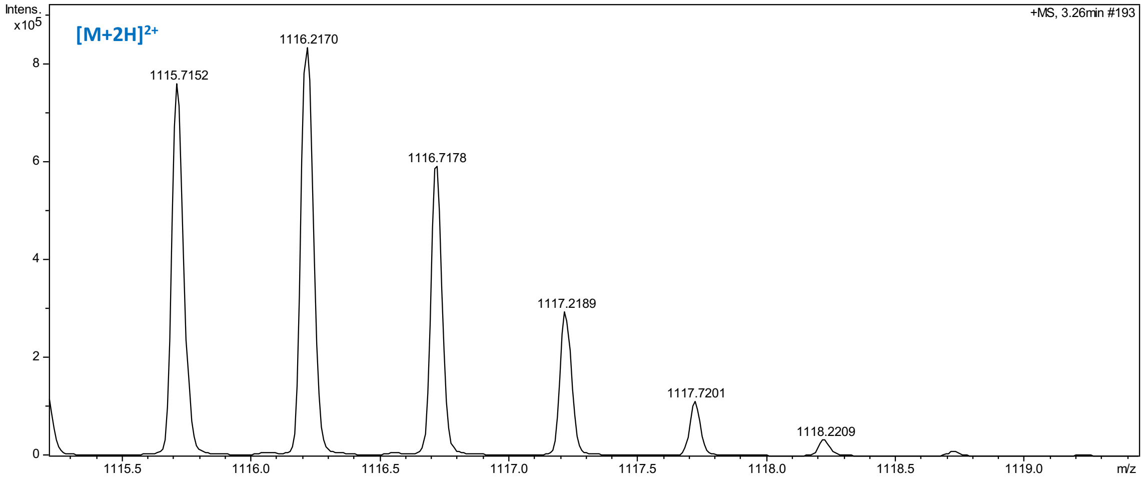


e) HRESIMS(+)-TOF spectra (ISCID 0 eV) of **1**. Doubly-charged adducts (zoom in region)

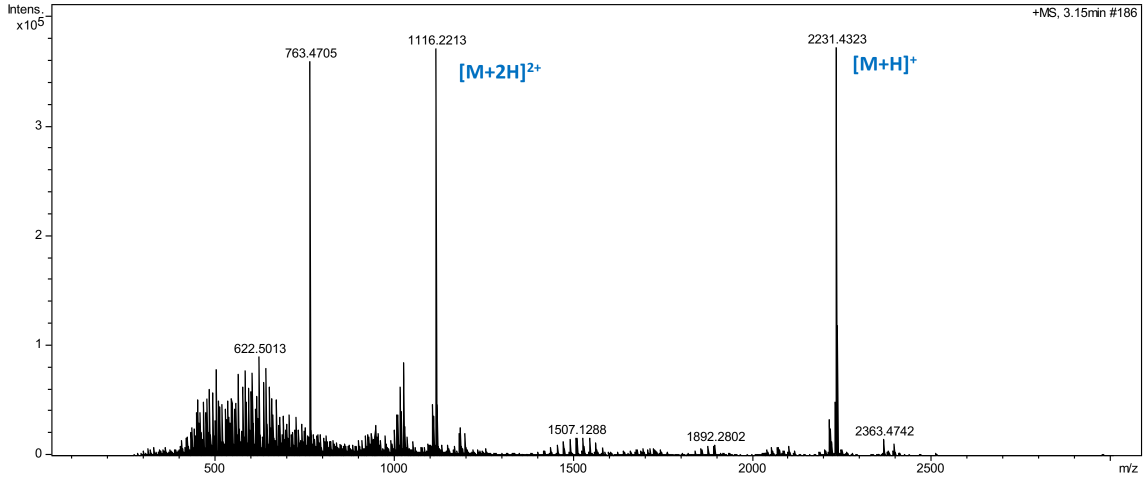


f) HRESIMS(+)-TOF spectra (ISCID 75 eV) of **2** (overview)

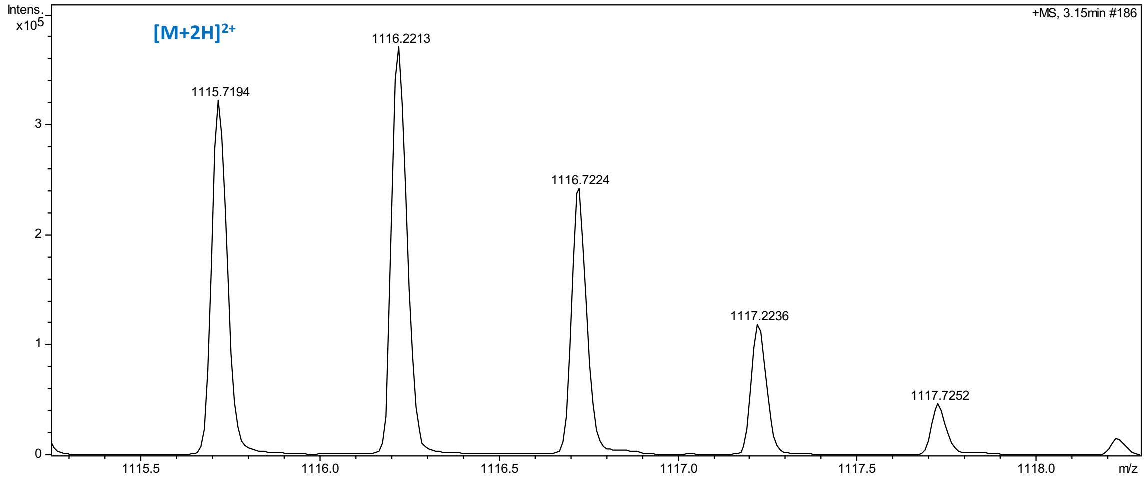


g) HRESIMS(+)-TOF spectra (ISCID 75 eV) of **2**. $[M+H]^+$ adduct (zoom in region)

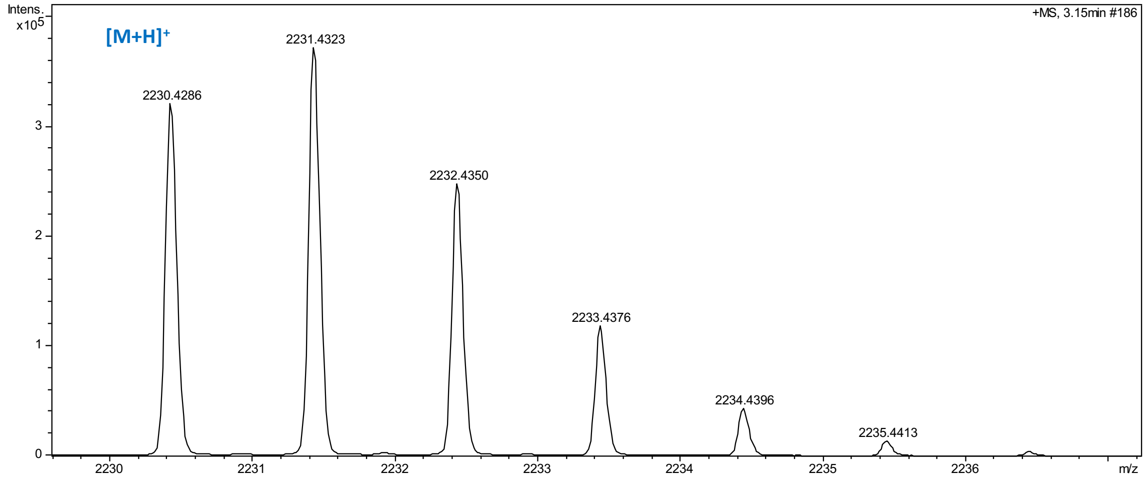


h) HRESIMS(+)-TOF spectra (ISCID 75 eV) of **2**. $[M+2H]^{2+}$ adduct (zoom in region)

**Table S1.** NMR spectroscopic data ($CD_3OD$, 500 MHz) for compounds **1** and **2**.

| Gargantulide B (**1**) | | | Gargantulide C (**2**) | | |
|---|---|---|---|---|---|
| **nº** | **δ $^{13}$C** | **δ $^{1}$H (mult, *J*, Hz)** | **nº** | **δ $^{13}$C** | **δ $^{1}$H (mult, *J*, Hz)** |
| 1 | 175.3, C | | 1 | 175.3, C | |
| 2 | 35.5, $CH_2$ | 2.36, m | 2 | 35.6, $CH_2$ | 2.36, m |
| 3 | 26.9, $CH_2$ | 1.61 [a], m | 3 | 26.9, $CH_2$ | 1.62 [a], m |
| 4 | 28.5 [a], $CH_2$ | 1.32 [a], m | 4 | 28.5 [a], $CH_2$ | 1.33 [a], m |
| | | 1.43 [a], m | | | 1.43 [a], m |
| 5 | 34.4 [b], $CH_2$ | 1.19, m | 5 | 34.4 [a], $CH_2$ | 1.19, m |
| | | 1.50 [a], m | | | 1.50 [a], m |
| 6 | 39.9, CH | 1.48 [a], m | 6 | 39.9, CH | 1.48 [a], m |
| Me-6 | 14.6, $CH_3$ | 0.88 [a], d (6.7) | Me-6 | 14.6, $CH_3$ | 0.89 [a], d (6.7) |
| 7 | 76.2, CH | 3.42, m | 7 | 76.2, CH | 3.42, m |
| 8 | 32.9, $CH_2$ | 1.46 [a], m | 8 | 32.9, $CH_2$ | 1.47 [a], m |
| 9 | 32.5, $CH_2$ | 1.38 [a], m | 9 | 32.5, $CH_2$ | 1.38 [a], m |
| | | 1.46 [a], m | | | 1.47 [a], m |
| 10 | 36.2, CH | 1.65 [a], m | 10 | 36.2, CH | 1.65 [a], m |
| Me-10 | 13.4, $CH_3$ | 0.86 [a], d (6.7) | Me-10 | 13.3, $CH_3$ | 0.86 [a], d (6.7) |
| 11 | 79.9, CH | 3.12, dd (8.2, 3.2) | 11 | 79.9, CH | 3.12, dd (8.3, 3.2) |
| 12 | 37.6, CH | 1.55 [a], m | 12 | 37.6, CH | 1.55, m |
| Me-12 | 16.8, $CH_3$ | 0.86 [a], d (6.7) | Me-12 | 16.8, $CH_3$ | 0.86 [a], d (6.8) |
| 13 | 34.0, $CH_2$ | 1.10 [a], m | 13 | 34.1, $CH_2$ | 1.10 [a], m |
| | | 1.75, m | | | 1.75, m |
| 14 | 28.5 [a], $CH_2$ | 1.48 [a], m | 14 | 28.5 [a], $CH_2$ | 1.48 [a], m |
| | | 1.22, m | | | 1.22, m |
| 15 | 27.2, $CH_2$ | 1.33 [a], m | 15 | 27.1, $CH_2$ | 1.32 [a], m |
| | | 1.57 [a], m | | | 1.57 [a], m |
| 16 | 35.6, $CH_2$ | 1.35 [a], m | 16 | 35.9, $CH_2$ | 1.35 [a], m |
| | | 1.57 [a], m | | | 1.57 [a], m |
| 17 | 74.8, CH | 3.67 [a], m | 17 | 74.8, CH | 3.60 [a], m |
| 18 | 54.1, CH | 2.69, dq (8.6, 6.9) | 18 | 53.9, CH | 2.68, dq (8.6, 7.0) |
| Me-18 | 13.5, $CH_3$ | 1.00, d (6.9) | Me-18 | 13.7, $CH_3$ | 0.99, d (6.8) |
| 19 | 214.6, C | | 19 | 214.6, C | |
| 20 | 49.9, $CH_2$ | 2.72, dd (17.2, 5.1) | 20 | 49.9, $CH_2$ | 2.80, dd (17.6, 6.9) |
| | | 2.89, dd (17.2, 7.8) | | | 2.87, dd (17.6, 6.8) |
| 21 | 74.1, CH | 4.49, dd (7.8, 5.1) | 21 | 74.1, CH | 4.47, dd (6.9, 6.8) |
| 22 | 146.8, C | | 22 | 146.7, C | |
| Me-22 | 12.6, $CH_3$ | 1.81 [a], s | Me-22 | 11.9, $CH_3$ | 1.78 [a], s |
| 23 | 123.2, CH | 5.21, d (10.3) | 23 | 123.5, CH | 5.22, d (10.3) |
| 24 | 76.1, CH | 4.76, dd (10.3, 8.1) | 24 | 75.8, CH | 4.81, dd (10.3, 8.0) |
| 25 | 80.6, CH | 3.56, dd (8.1, 3.5) | 25 | 80.4, CH | 3.64, dd (8.0, 2.8) |
| 26 | 39.3, CH | 1.69 [a], m | 26 | 39.4, CH | 1.74 [a], m |
| Me-26 | 11.9, $CH_3$ | 1.06, d (7.0) | Me-26 | 11.8, $CH_3$ | 1.06, d (7.2) |
| 27 | 68.6, CH | 4.22 [a], m | 27 | 68.8 [a], CH | 4.20 [a], m |
| 28 | 43.9, $CH_2$ | 1.27, m | 28 | 43.9, $CH_2$ | 1.27, m |
| | | 1.59 [a], m | | | 1.59 [a], m |
| 29 | 68.6 [a], CH | 3.64 [a], m | 29 | 68.8 [a], CH | 3.61 [a], m |
| 30 | 39.2 [a], $CH_2$ | 1.40 [a], m | 30 | 39.2 [a], $CH_2$ | 1.41 [a], m |
| | | 1.44 [a], m | | | 1.44 [a], m |
| 31 | 23.1, $CH_2$ | 1.53 [a], m | 31 | 23.2, $CH_2$ | 1.51 [a], m |
| 32 | 39.2 [a], $CH_2$ | 1.48 [a], m | 32 | 39.2 [a], $CH_2$ | 1.48 [a], m |

| | | | | | |
|---|---|---|---|---|---|
| | | 1.52 [a], m | | | 1.52 [a], m |
| 33 | 69.4, CH | 3.84 [a], m | 33 | 69.4, CH | 3.84 [a], m |
| 34 | 46.0 [a], $CH_2$ | 1.51 [a], m | 34 | 46.3 [a], $CH_2$ | 1.51 [a], m |
| | | 1.59 [a], m | | | 1.59 [a], m |
| 35 | 66.6, CH | 4.10 [a], m | 35 | 66.6, CH | 4.10 [a], m |
| 36 | 46.9 [a], $CH_2$ | 1.57 [a], m | 36 | 46.9 [a], $CH_2$ | 1.58 [a], m |
| 37 | 66.6 [a], CH | 4.10 [a], m | 37 | 66.6, CH | 4.10 [a], m |
| 38 | 46.9 [a], $CH_2$ | 1.58 [a], m | 38 | 46.9 [a], $CH_2$ | 1.58 [a], m |
| 39 | 66.6 [a], CH | 4.08 [a], m | 39 | 66.6 [a], CH | 4.08 [a], m |
| 40 | 44.1, $CH_2$ | 1.57 [a], m | 40 | 44.2, $CH_2$ | 1.57 [a], m |
| | | 1.61 [a], m | | | 1.61 [a], m |
| 41 | 73.8, CH | 4.04, m | 41 | 73.8, CH | 4.03, m |
| 42 | 41.1, CH | 1.63 [a], m | 42 | 40.9, CH | 1.63 [a], m |
| Me-42 | 6.9, $CH_3$ | 0.95 [a], d (6.8) | Me-42 | 6.8, $CH_3$ | 0.95 [a], d (6.9) |
| 43 | 77.5, CH | 3.80 [a], m | 43 | 77.5, CH | 3.80 [a], m |
| 44 | 41.9, CH | 1.67 [a], m | 44 | 41.9, CH | 1.67 [a], m |
| Me-44 | 10.8, $CH_3$ | 0.84, d (6.9) | Me-44 | 10.8, $CH_3$ | 0.84, d (6.8) |
| 45 | 72.4, CH | 4.23 [a], m | 45 | 72.4, CH | 4.22 [a], m |
| 46 | 40.8, $CH_2$ | 1.58 [a], m | 46 | 40.7, $CH_2$ | 1.57 [a], m |
| 47 | 71.6, CH | 4.06, m | 47 | 71.6, CH | 4.05, m |
| 48 | 38.2, $CH_2$ | 1.58 [a], m | 48 | 38.3, $CH_2$ | 1.57 [a], m |
| | | 1.67 [a], m | | | 1.66 [a], m |
| 49 | 77.1, CH | 3.69 [a], m | 49 | 77.1, CH | 3.69 [y], m |
| 50 | 38.5, CH | 2.30, m | 50 | 38.5, CH | 2.30, m |
| Me-50 | 10.8, $CH_3$ | 0.93 [a], d (6.9) | Me-50 | 10.7, $CH_3$ | 0.93 [a], d (6.9) |
| 51 | 76.9, CH | 4.88, m | 51 | 77.0, CH | 4.88, m |
| 52 | 41.1, CH | 1.52 [a], m | 52 | 41.1, CH | 1.52 [a], m |
| 53 | 24.7, $CH_2$ | 1.20 [a], m | 53 | 24.6, $CH_2$ | 1.20 [a], m |
| | | 1.78 [a], m | | | 1.78 [a], m |
| 54 | 31.9, $CH_2$ | 1.44 [a], m | 54 | 31.8, $CH_2$ | 1.45 [a], m |
| | | 1.89, m | | | 1.89, m |
| 55 | 83.9, CH | 3.68 [a], m | 55 | 83.8, CH | 3.67 [a], m |
| 56 | 43.4, CH | 1.73, m | 56 | 43.3, CH | 1.73, m |
| Me-56 | 10.7, $CH_3$ | 0.94 [a], d (6.9) | Me-56 | 10.8, $CH_3$ | 0.94 [a], d (7.0) |
| 57 | 68.8, CH | 4.19 [a], m | 57 | 68.8, CH | 4.19 [a], m |
| 58 | 44.2, $CH_2$ | 1.39 [a], m | 58 | 44.6, $CH_2$ | 1.41 [a], m |
| | | 1.67 [a], m | | | 1.67 [a], m |
| 59 | 77.0 [a], CH | 3.84 [a], m | 59 | 68.8, CH | 3.61, m |
| 60 | 46.0 [a], $CH_2$ | 1.58 [a], m | 60 | 46.0 [a], $CH_2$ | 1.58 [a], m |
| | | 1.63 [a], m | | | 1.63 [a], m |
| 61 | 77.0 [a], CH | 3.84 [a], m | 61 | 76.9, CH | 3.83, m |
| 62 | 38.7, $CH_2$ | 1.53, m | 62 | 38.7, $CH_2$ | 1.53, m |
| | | 1.41 [a], m | | | 1.41 [a], m |
| 63 | 22.6, $CH_2$ | 1.41 [a], m | 63 | 22.5, $CH_2$ | 1.41 [a], m |
| | | 1.61 [a], m | | | 1.61 [a], m |
| 64 | 38.6, $CH_2$ | 1.40 [a], m | 64 | 38.5, $CH_2$ | 1.40 [a], m |
| | | 1.49 [a], m | | | 1.49 [a], m |
| 65 | 73.1 [a], CH | 3.50 [a], m | 65 | 72.9 [a], CH | 3.50 [a], m |
| 66 | 36.1, $CH_2$ | 1.36, m | 66 | 36.1, $CH_2$ | 1.35 [a], m |
| | | 1.51 [a], m | | | 1.51 [a], m |
| 67 | 34.2 [a], $CH_2$ | 1.10 [a], m | 67 | 34.2 [a], $CH_2$ | 1.11 [a], m |
| | | 1.50 [a], m | | | 1.50 [a], m |
| 68 | 34.2 [a], CH | 1.42 [a], m | 68 | 34.2 [a], CH | 1.43 [a], m |
| Me-68 | 20.2, $CH_3$ | 0.92 [a], d (6.5) | Me-68 | 20.2, $CH_3$ | 0.92 [a], d (6.5) |
| 69 | 37.7, $CH_2$ | 1.20 [a], m | 69 | 37.7, $CH_2$ | 1.19 [a], m |

| | | | | | |
|---|---|---|---|---|---|
| | | 1.39 [a], m | | | 1.38 [a], m |
| 70 | 25.1, $CH_2$ | 1.39 [a], m | 70 | 25.1, $CH_2$ | 1.39 [a], m |
| | | 1.45 [a], m | | | 1.44 [a], m |
| 71 | 29.1, $CH_2$ | 1.64 [a], m | 71 | 29.1, $CH_2$ | 1.64 [a], m |
| 72 | 40.9, $CH_2$ | 2.92, t (7.7) | 72 | 40.9, $CH_2$ | 2.92, t (7.7) |
| 1’ | 34.4 [b], $CH_2$ | 1.09, m | 1’ | 34.4 [a], $CH_2$ | 1.10, m |
| | | 1.36 [a], m | | | 1.35 [a], m |
| 2’ | 21.6, $CH_2$ | 1.29, m | 2’ | 21.6, $CH_2$ | 1.28, m |
| | | 1.48 [a], m | | | 1.48 [a], m |
| 3’ | 14.9, $CH_3$ | 0.89 [a], t (7.2) | 3’ | 14.9, $CH_3$ | 0.89 [a], t (7.2) |
| Man-1 | 97.1, CH | 4.58, br s | Man-1 | 96.9, CH | 4.58, br s |
| Man-2 | 73.1 [a], CH | 3.81 [a], m | Man-2 | 73.1 [a], CH | 3.81 [a], m |
| Man-3 | 75.4, CH | 3.47, dd (9.5, 3.2) | Man-3 | 75.4, CH | 3.47, dd (9.5, 3.2) |
| Man-4 | 68.6 [a], CH | 3.67 [a], t (9.5) | Man-4 | 68.8 [a], CH | 3.61 [a], t (9.5) |
| Man-5 | 78.5, CH | 3.17, m | Man-5 | 78.6, CH | 3.15, m |
| Man-6 | 62.8 [a], $CH_2$ | 3.75 [m´], m | Man-6 | 62.9 [a], $CH_2$ | 3.74 [m´], m |
| | | 3.87 [a], m | | | 3.87 [a], m |
| Glc-1 | 102.4, CH | 4.20 [a], d (7.8) | Glc-1 | 102.4, CH | 4.22 [a], d (7.8) |
| Glc-2 | 74.9, CH | 3.20 [a], m | Glc-2 | 74.9, CH | 3.20 [a], m |
| Glc-3 | 78.2, CH | 3.33 [a], m | Glc-3 | 78.2, CH | 3.33 [a], m |
| Glc-4 | 71.8, CH | 3.32 [a], m | Glc-4 | 71.7, CH | 3.33 [a], m |
| Glc-5 | 78.3 [a], CH | 3.21 [a], m | Glc-5 | 78.3 [a], CH | 3.21 [a], m |
| Glc-6 | 62.8 [a], $CH_2$ | 3.68 [a], m | Glc-6 | 62.9 [a], $CH_2$ | 3.68 [a], m |
| | | 3.84 [a], m | | | 3.85 [a], m |
| maG-1 | 104.4, CH | 4.46, d (7.5) | maG-1 | 104.7, CH | 4.45, d (7.4) |
| maG-2 | 70.5, CH | 3.50 [a], dd (10.4, 7.5) | maG-2 | 70.6, CH | 3.49 [a], dd (10.6, 7.4) |
| maG-3 | 66.1, CH | 3.06, t (10.4) | maG-3 | 66.0, CH | 3.05, t (10.6) |
| maG-4 | 71.5, CH | 3.39 [a], dd (10.4, 8.7) | maG-4 | 71.5, CH | 3.38 [a], dd (10.6, 8.8) |
| maG-5 | 74.4, CH | 3.40, dq (8.7, 6.2) | maG-5 | 74.3, CH | 3.41, dq (8.8, 6.1) |
| maG-6 | 18.4, $CH_3$ | 1.34 [a], d (6.2) | maG-6 | 18.4, $CH_3$ | 1.33 [a], d (6.1) |
| N-Me | 31.1, $CH_3$ | 2.81, br s | N-Me | 31.1, $CH_3$ | 2.81, br s |
| Ara-1 | 108.3, CH | 5.01, d (1.6) | Ara-1 | 108.2, CH | 5.01, d (1.5) |
| Ara-2 | 83.9, CH | 3.96, dd (3.9, 1.6) | Ara-2 | 83.9, CH | 3.96, dd (3.9, 1.5) |
| Ara-3 | 78.7, CH | 3.86 [a], m | Ara-3 | 78.6, CH | 3.85 [a], m |
| Ara-4 | 85.4, CH | 3.98, m | Ara-4 | 85.4, CH | 3.98, m |
| Ara-5 | 63.2, $CH_2$ | 3.64 [a], m | Ara-5 | 63.2, $CH_2$ | 3.64 [a], m |
| | | 3.74 [a], m | | | 3.75 [a], m |
| Glc´-1 | 103.9, CH | 4.33, d (7.6) | | | |
| Glc´-2 | 75.2, CH | 3.21 [a], m | | | |
| Glc´-3 | 78.3 [a], CH | 3.37 [a], m | | | |
| Glc´-4 | 72.5, CH | 3.23 [a], m | | | |
| Glc´-5 | 78.3 [a], CH | 3.21 [a], m | | | |
| Glc´-6 | 63.4, $CH_2$ | 3.93, m | | | |
| | | 3.70 [a], m | | | |

[a] Overlapping with other isochronous $^{13}$C-NMR / $^{1}$H-NMR signals.

* No multiplicity is given (m) for $^{1}$H-NMR signals for which coupling constants (*Js*) could not be measured directly in $^{1}$H or JRES spectra nor determined from HSQC traces. The assignments were supported by HSQC, HMBC, TOCSY and HSQC-TOCSY.

**Figure S4.** NMR spectra of gargantulide B (**1**)

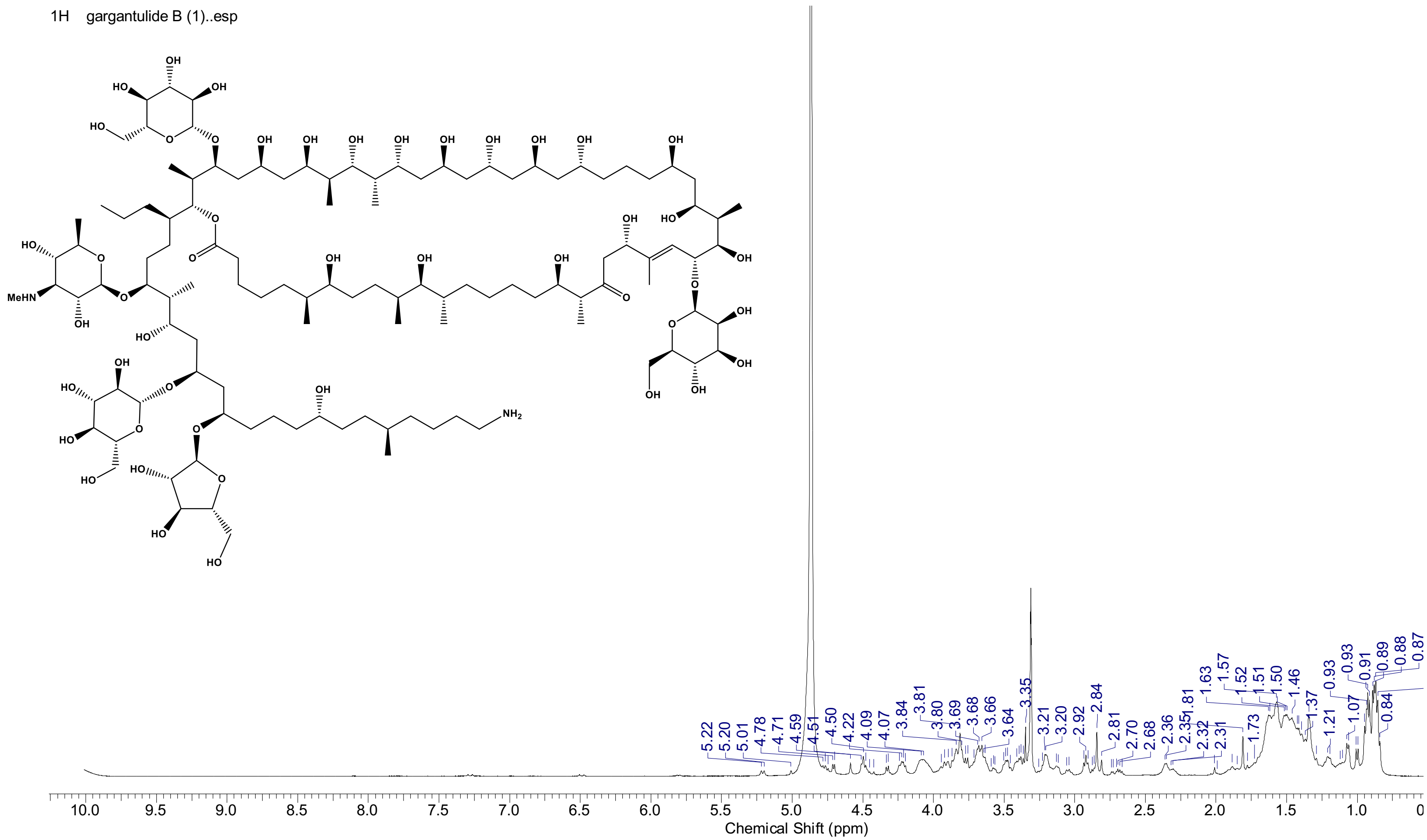

a) $^{1}$H NMR spectrum ($CD_3OD$, 500 MHz) of **1**. Full scale (0-10 ppm)

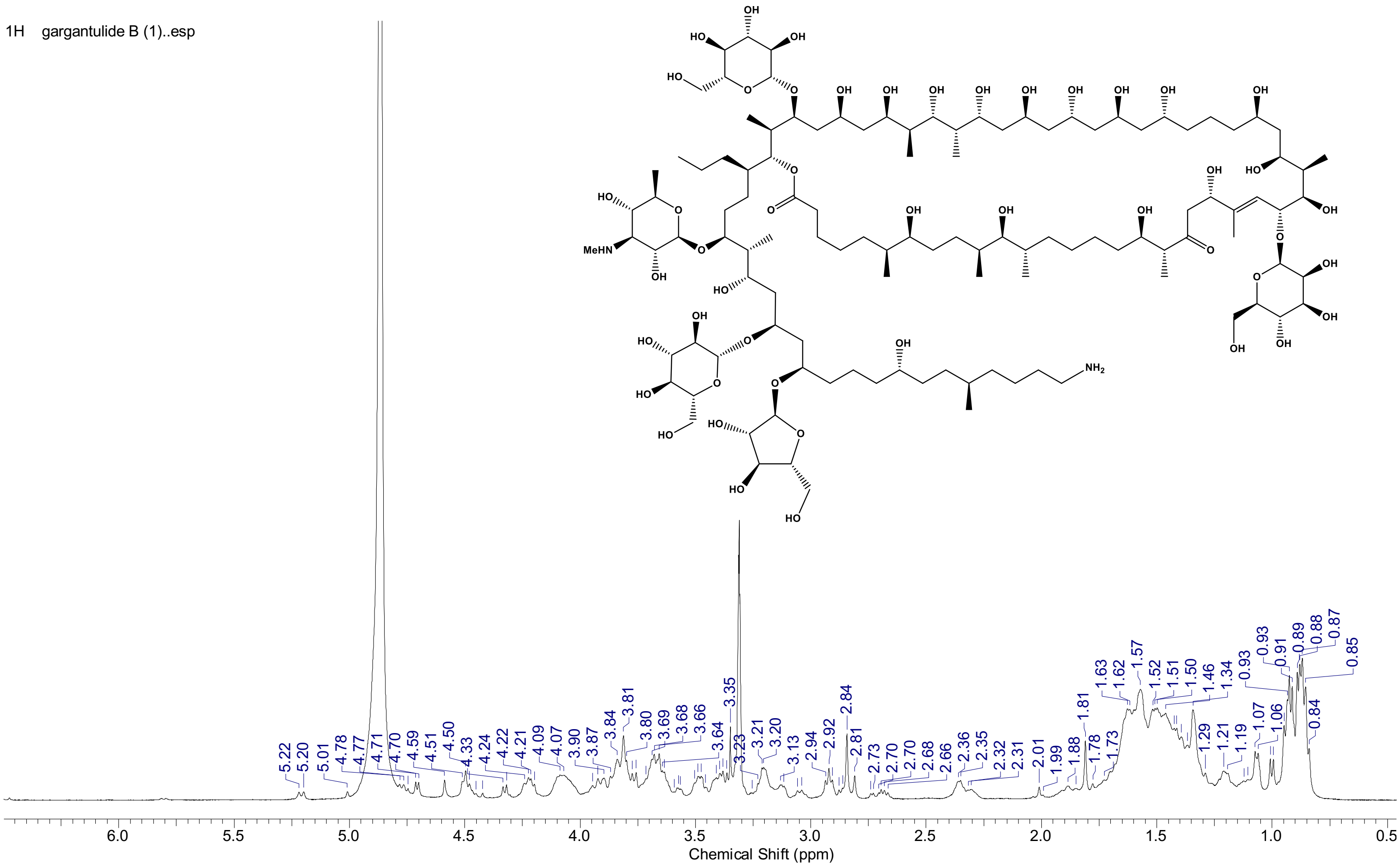


b) ¹H NMR spectrum ($CD_3OD$, 500 MHz) of **1**. Expansion between 0 and 7 ppm

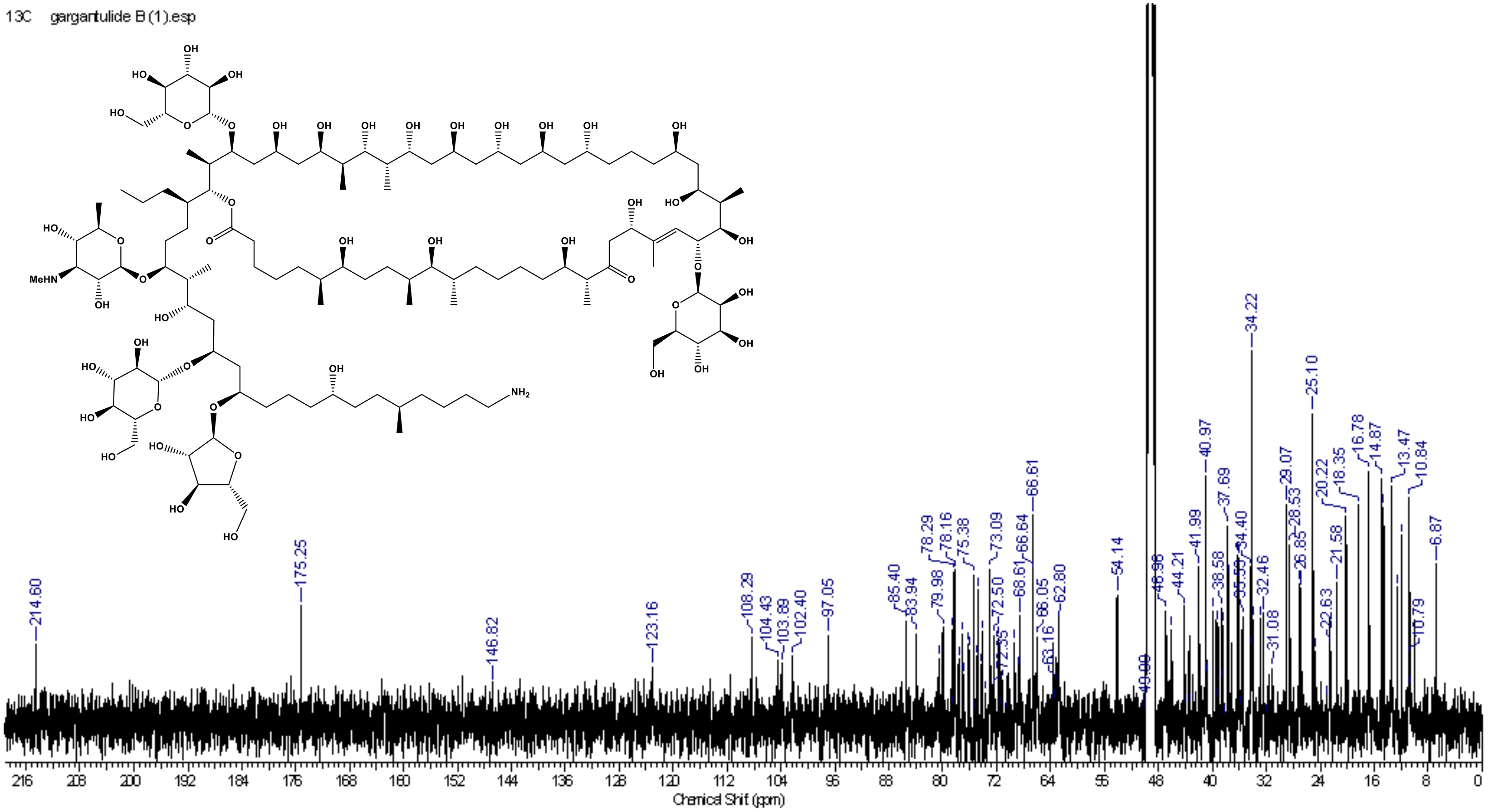


c) $^{13}C$ NMR spectrum ($CD_3OD$, 125 MHz) of **1**

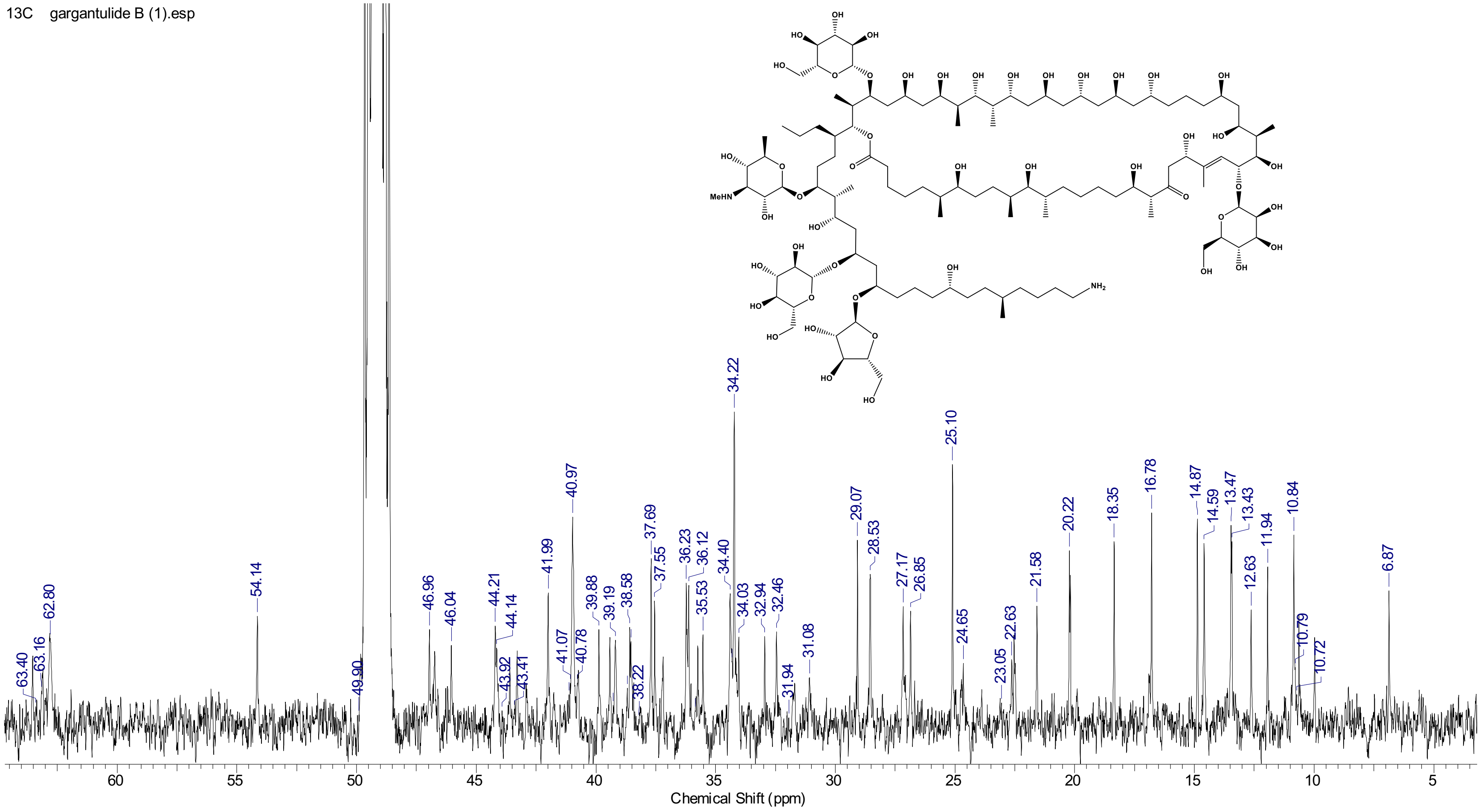


d) $^{13}C$ NMR spectrum ($CD_3OD$, 125 MHz) of **1**. Expansion between 0 and 65 ppm

13C gargantulide B (1).esp

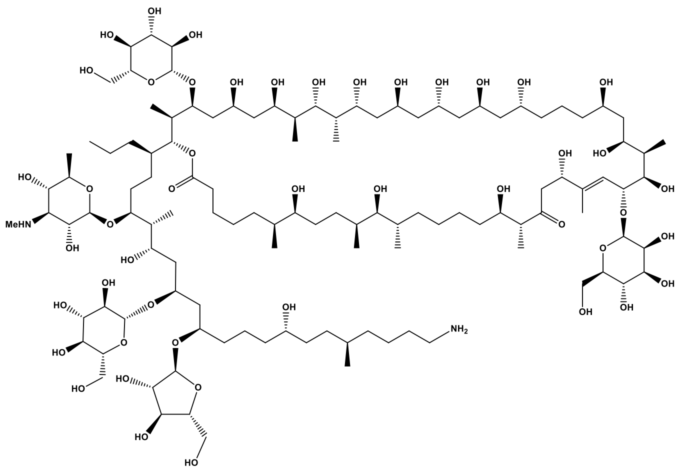

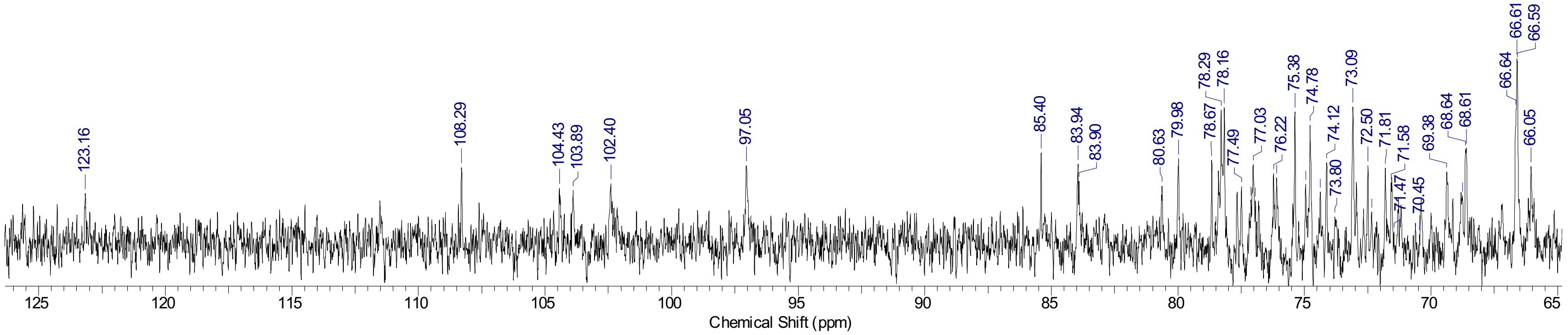


e) $^{13}$C NMR spectrum ($CD_3OD$, 125 MHz) of **1**. Expansion between 65 and 125 ppm

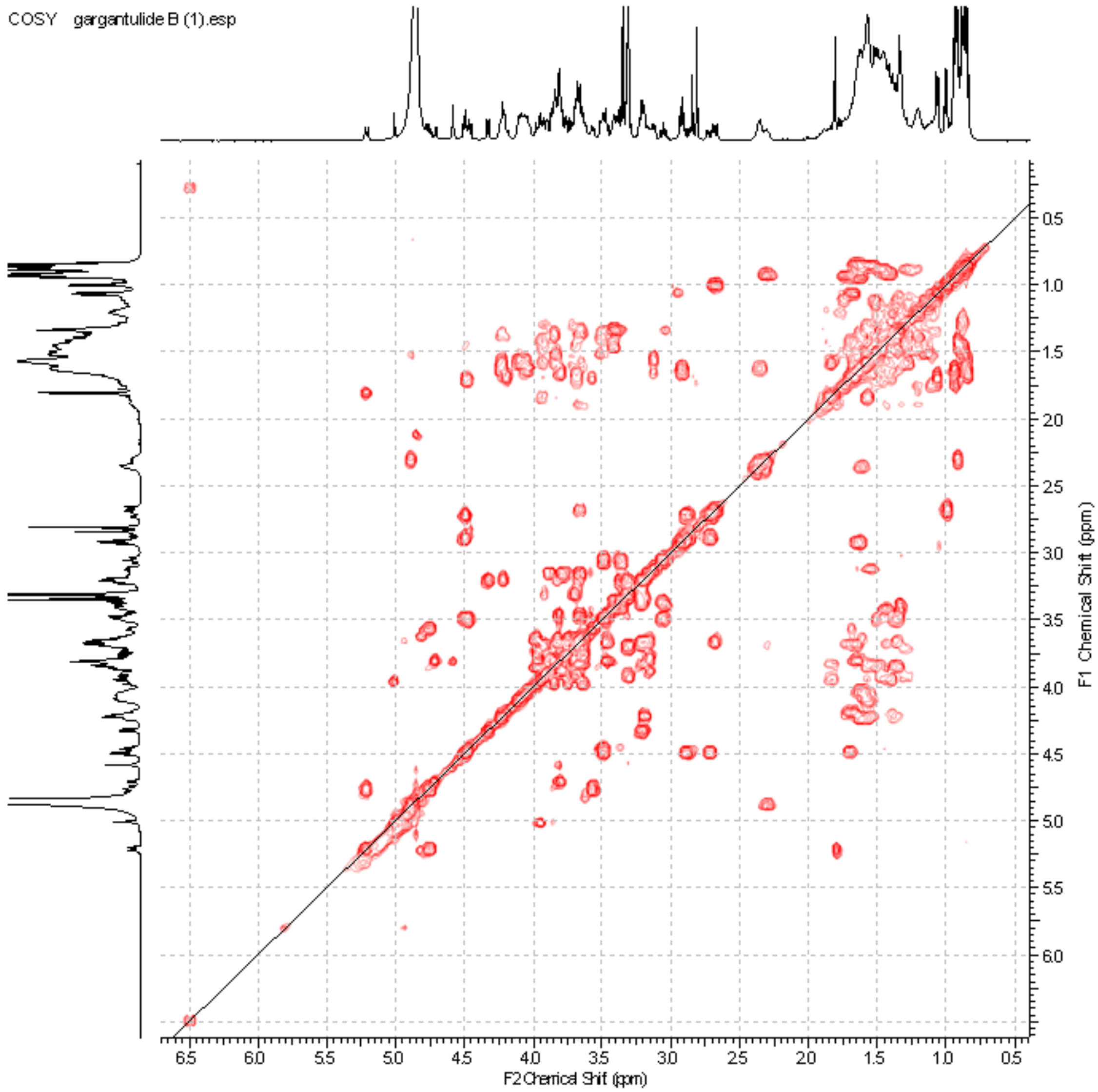


f) COSY spectrum of **1**

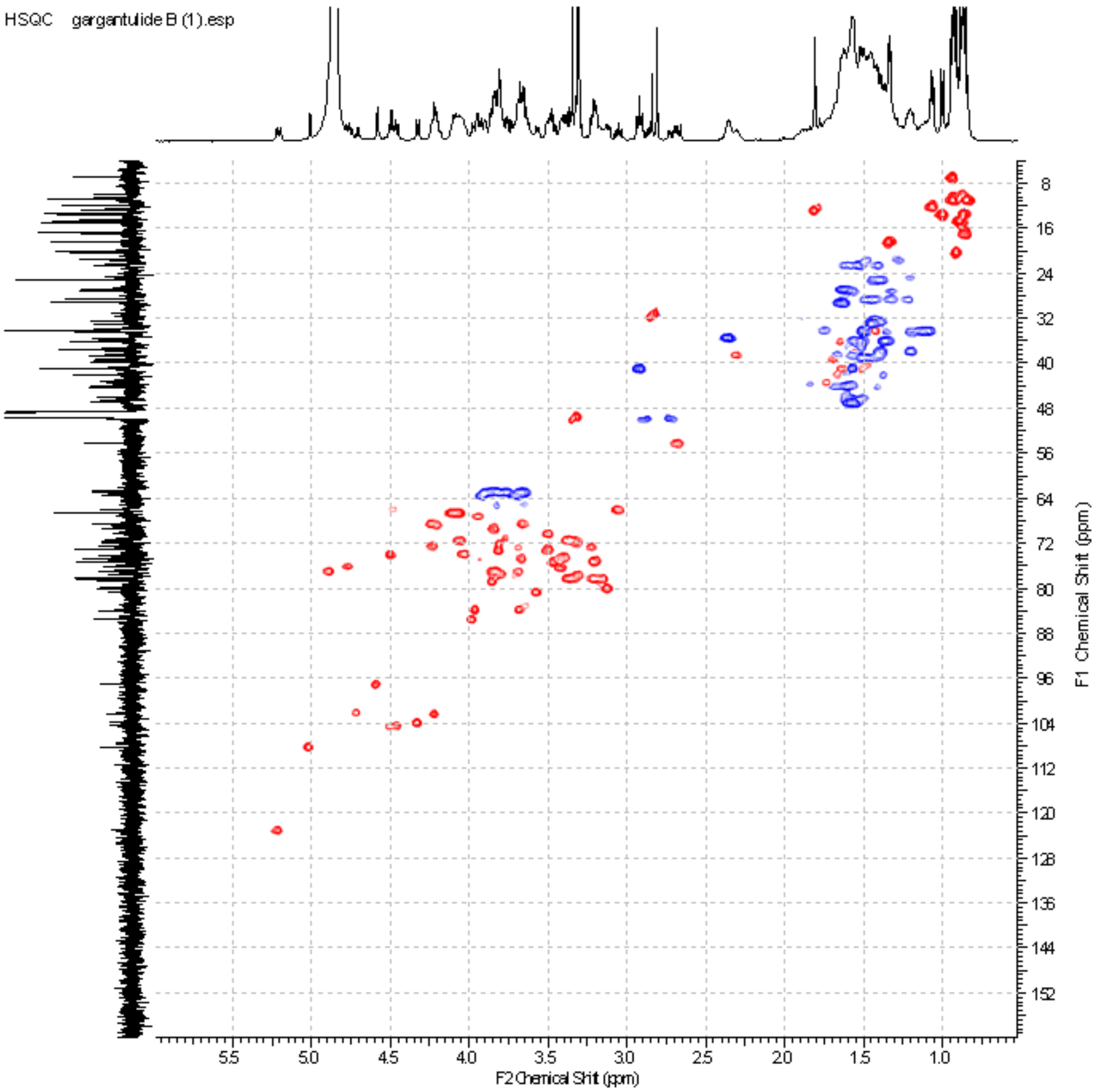


g) HSQC spectrum of **1**

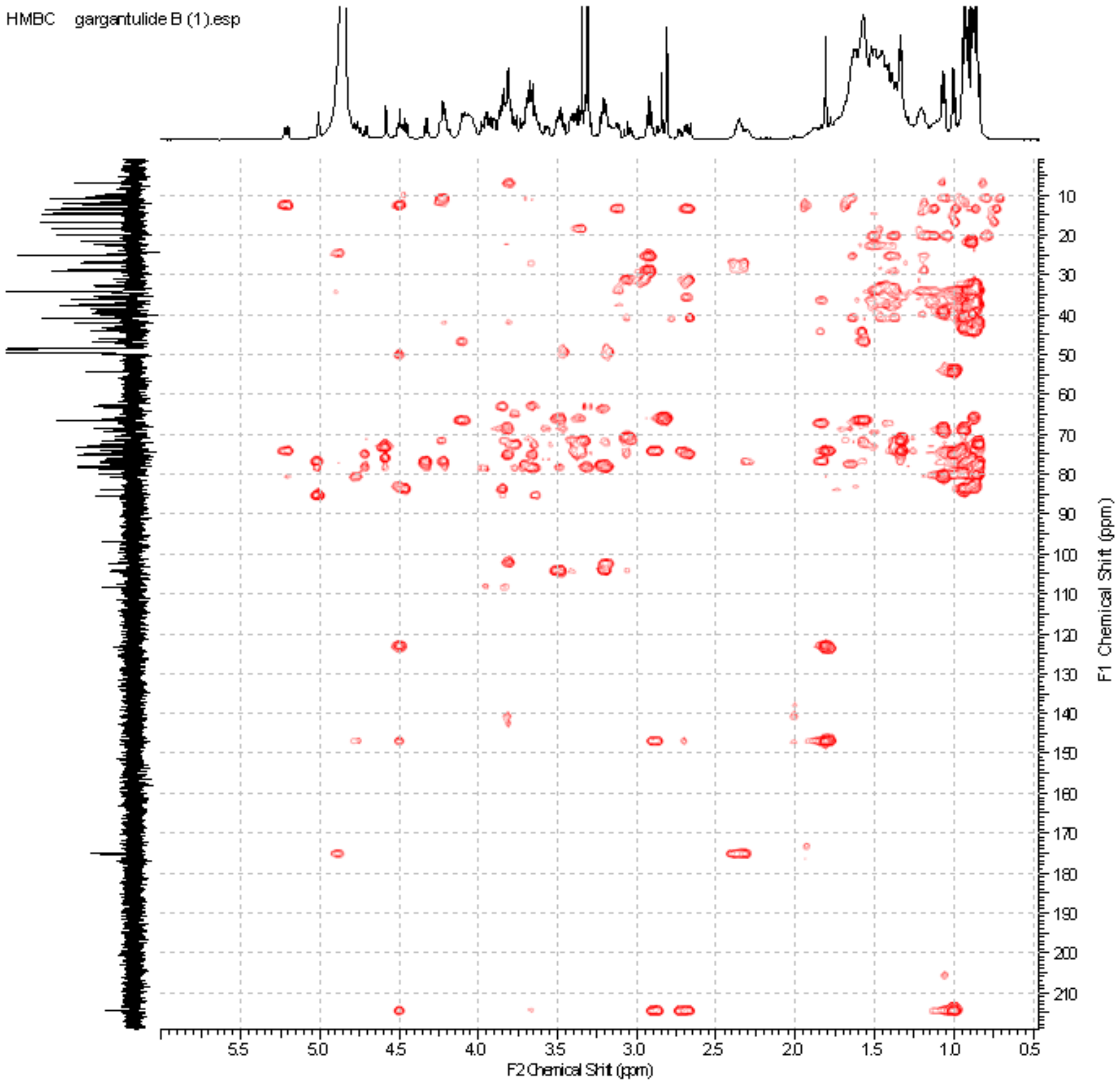


h) HMBC spectrum of **1**

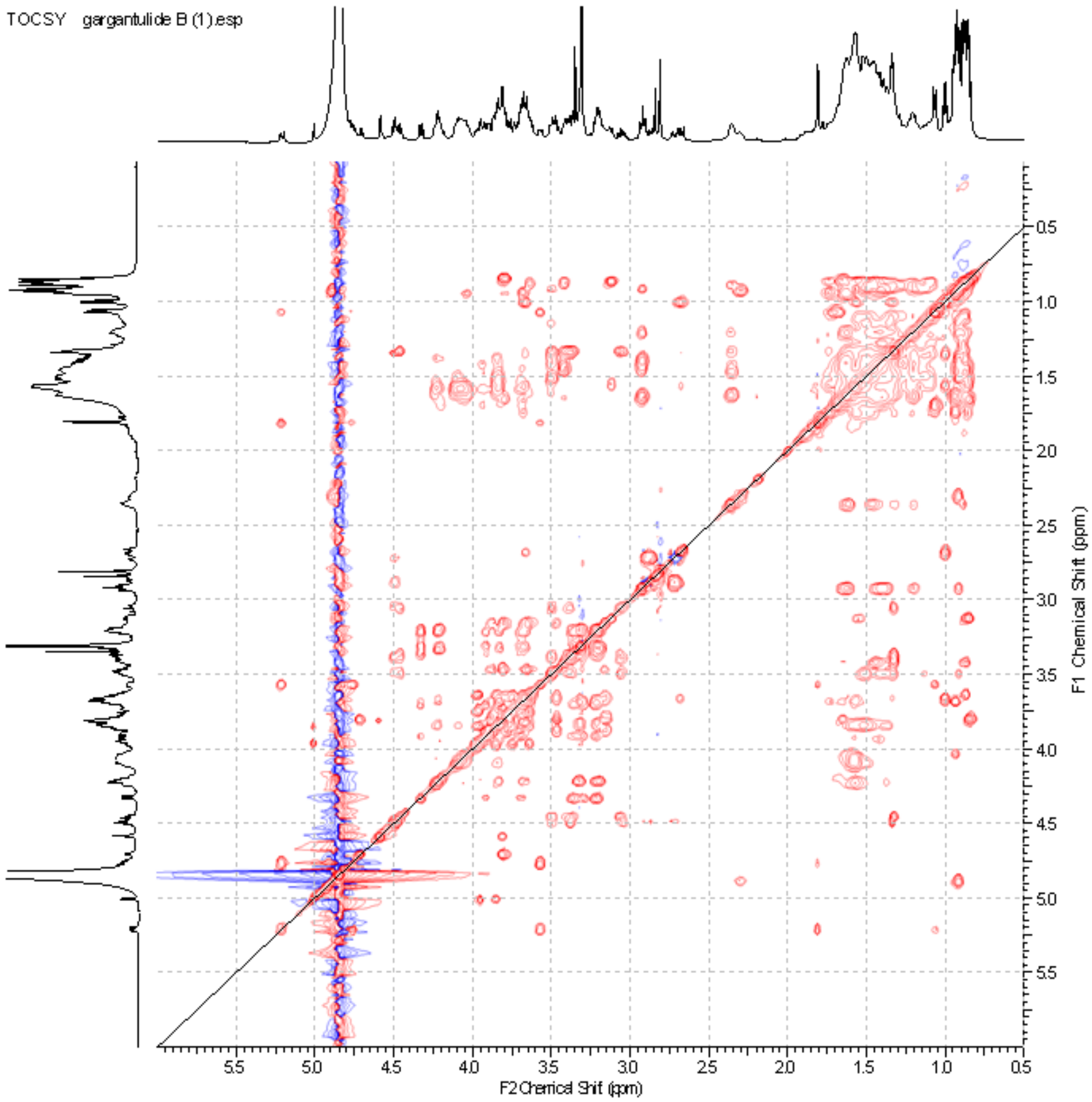


i) TOCSY spectrum of **1**

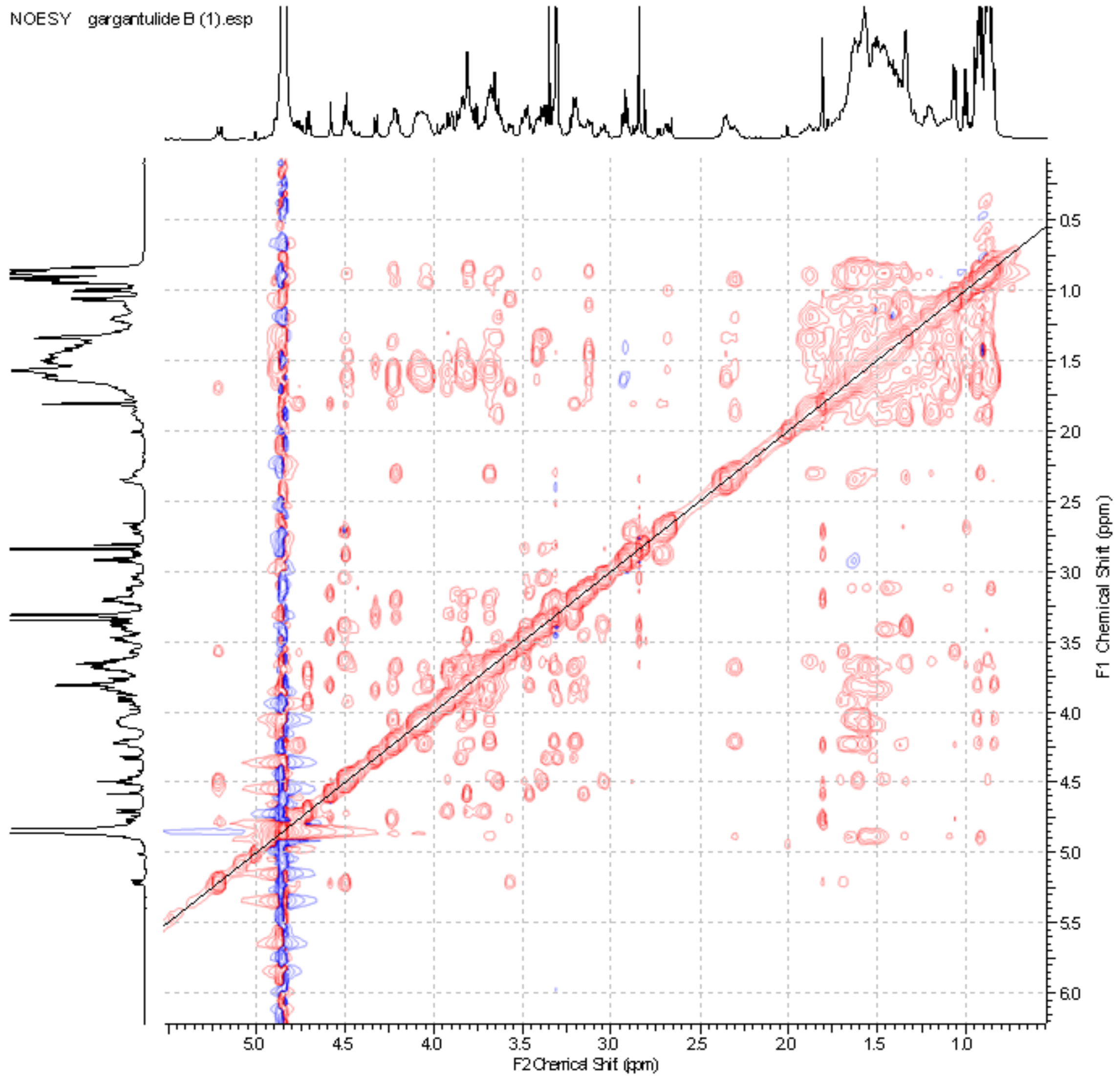


j) NOESY spectrum of **1**

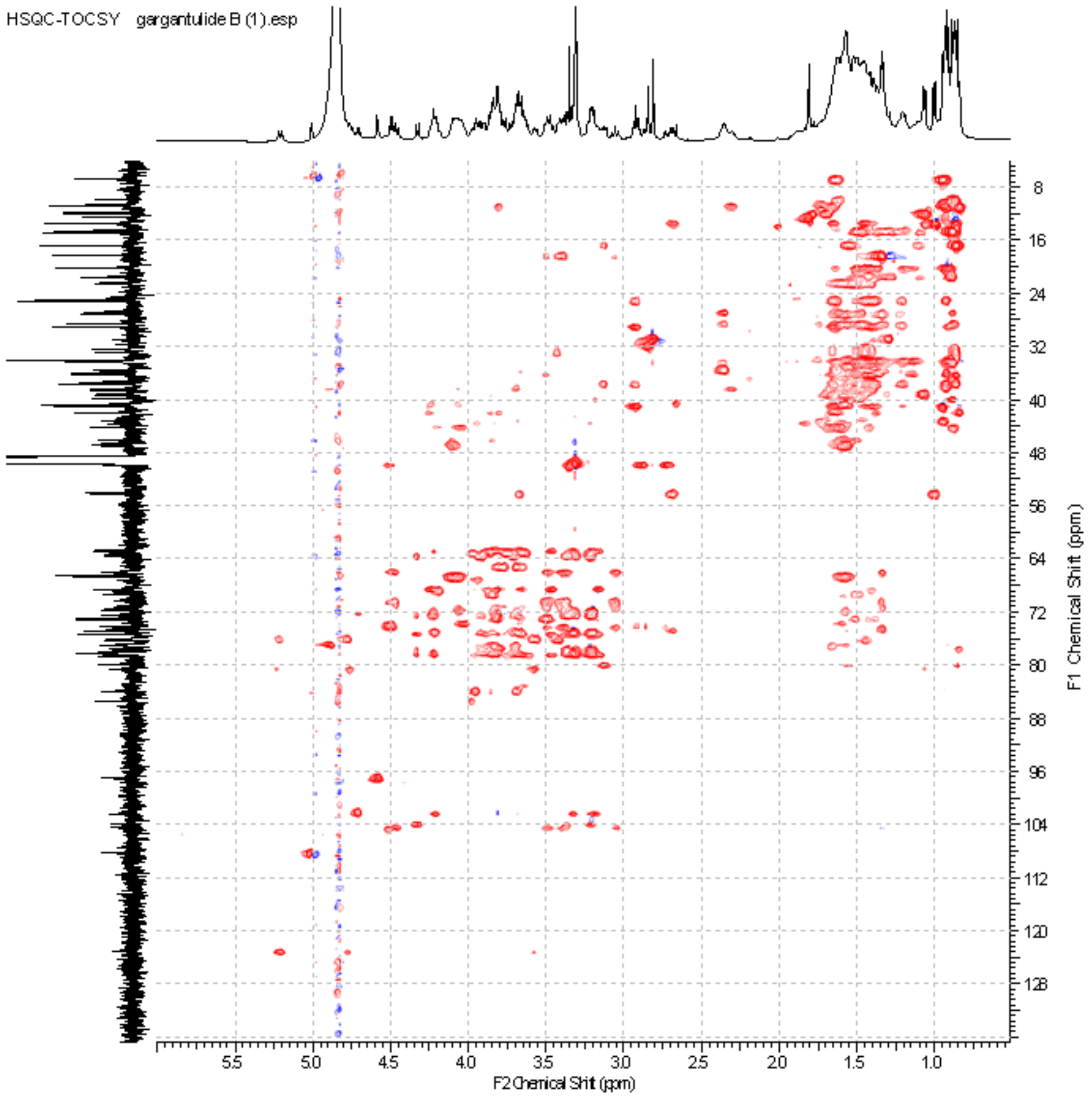


k) HSQC-TOCSY spectrum of **1**

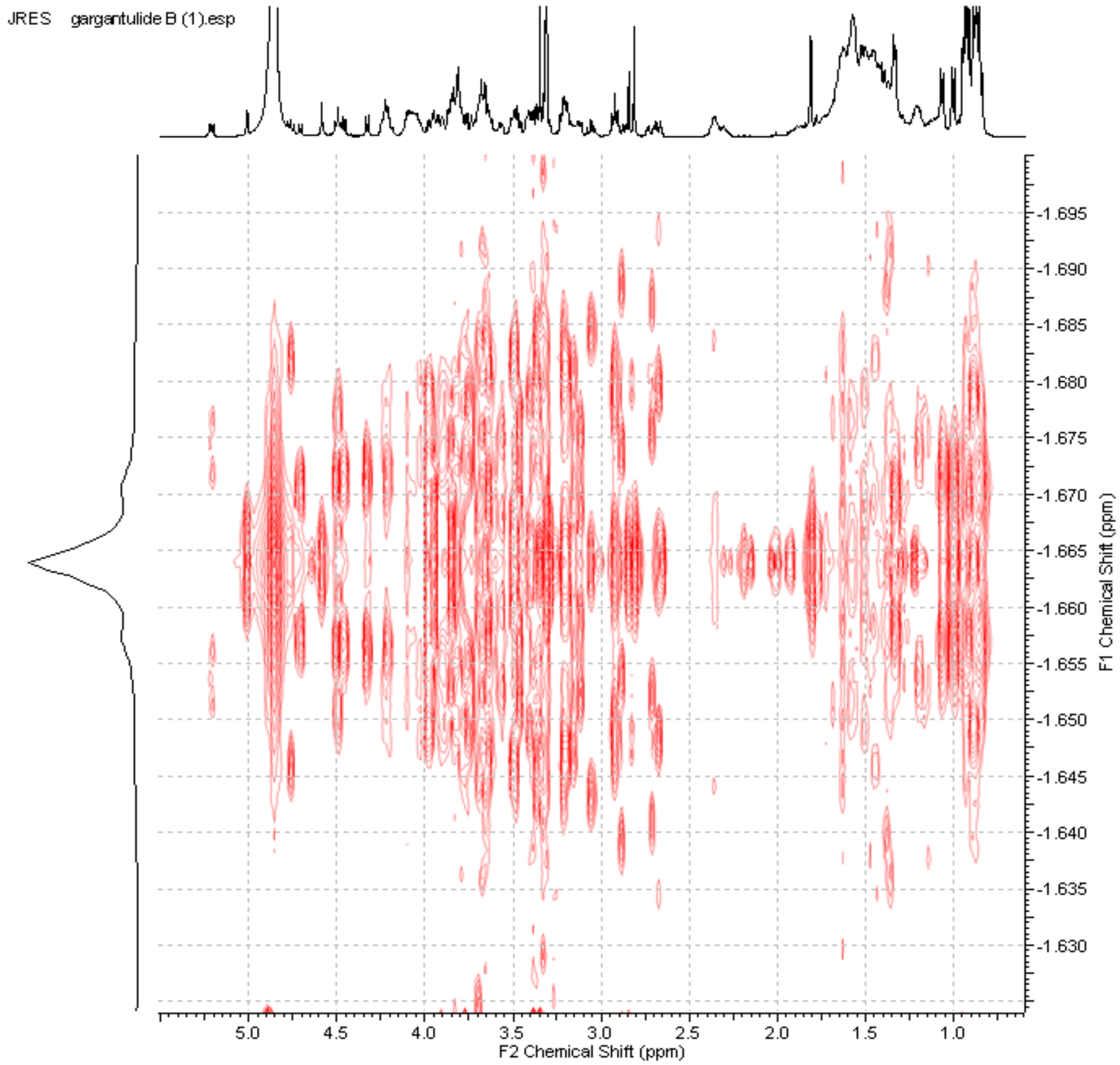


l) JRES spectrum of **1**

**Figure S5.** NMR spectra of gargantulide C (**2**)

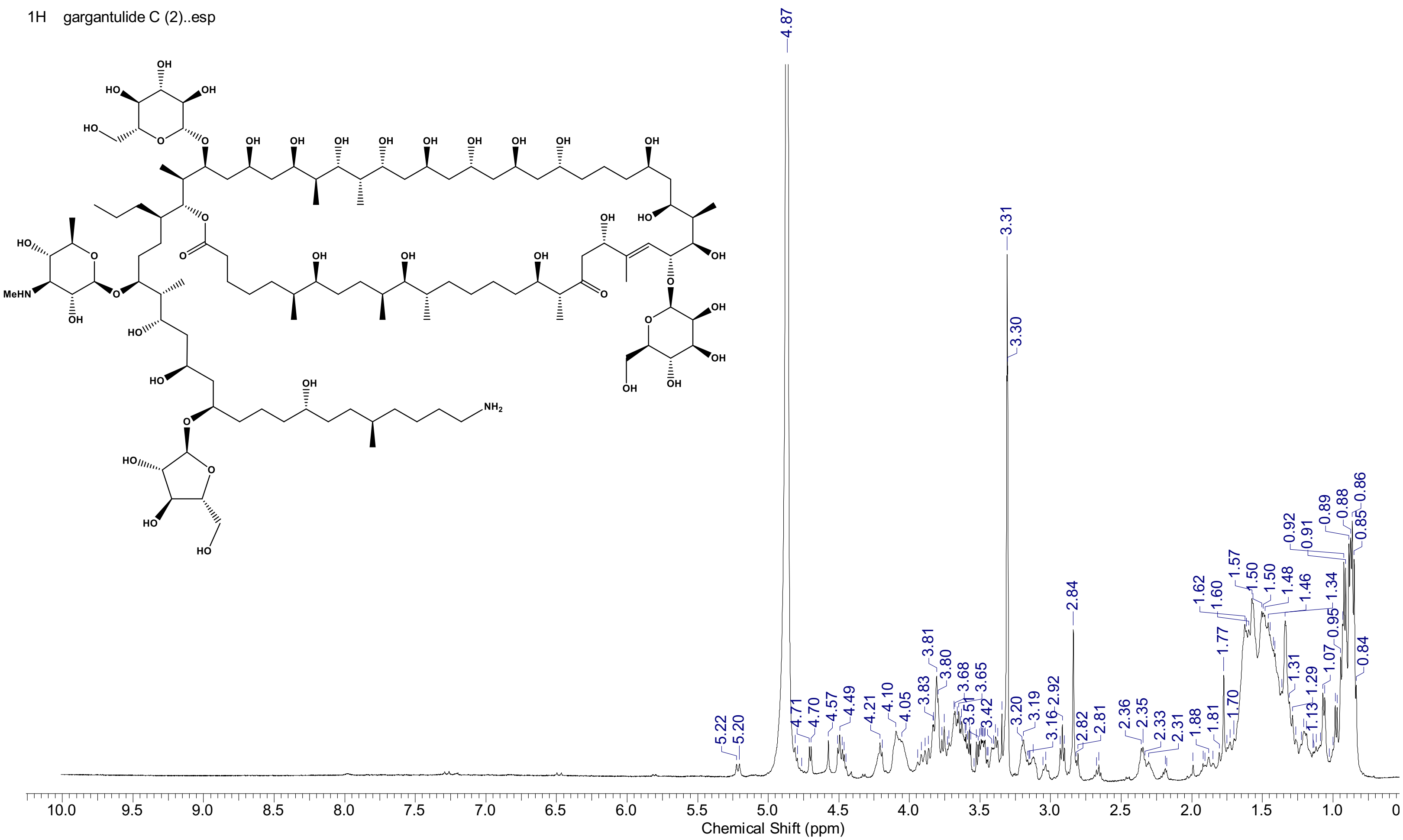


a) $^{1}H$ NMR spectrum ($CD_3OD$, 500 MHz) of **2**. Full scale (0-10 ppm)

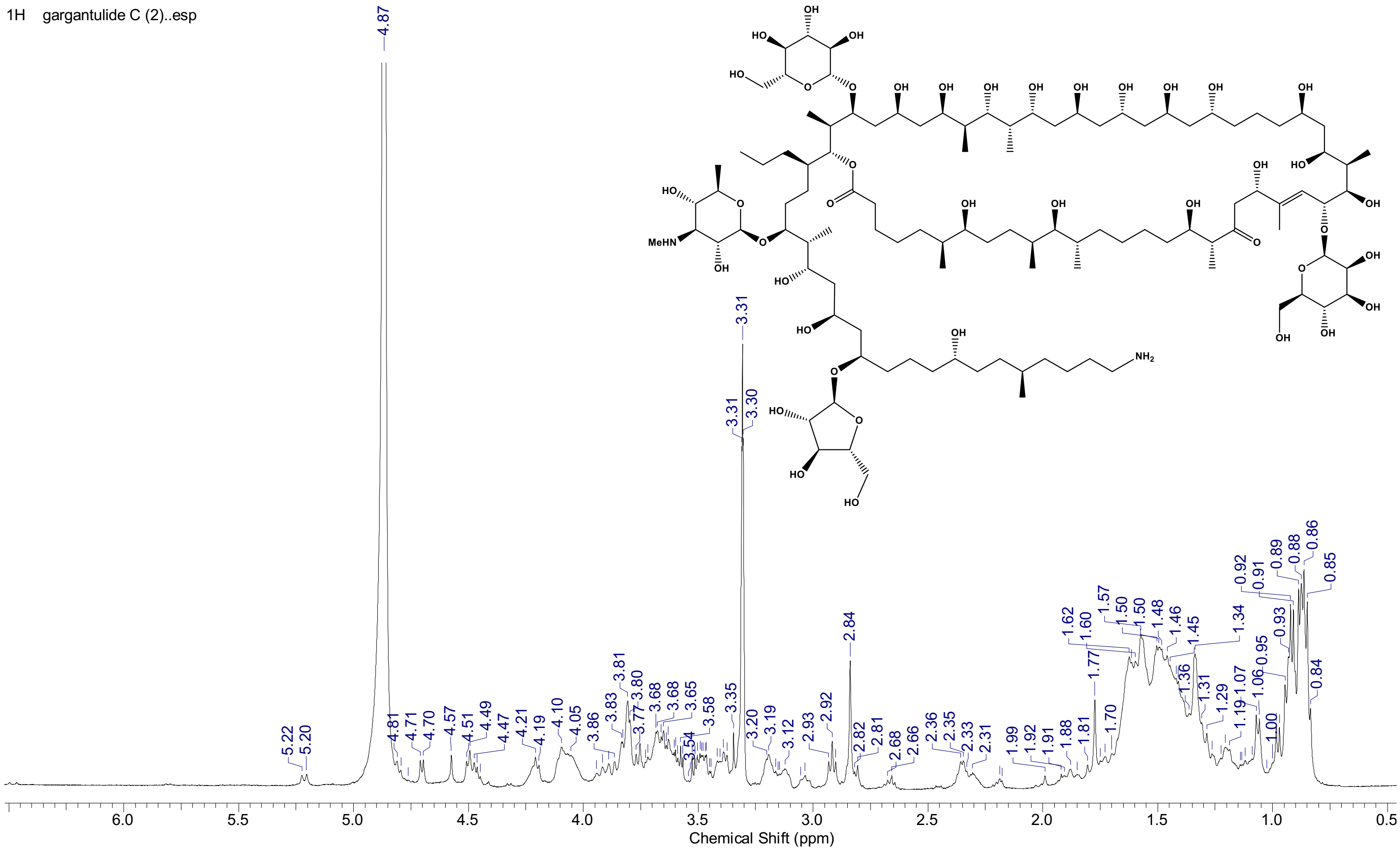


b) $^{1}H$ NMR spectrum ($CD_3OD$, 500 MHz) of **2**. Expansion between 0 and 7 ppm

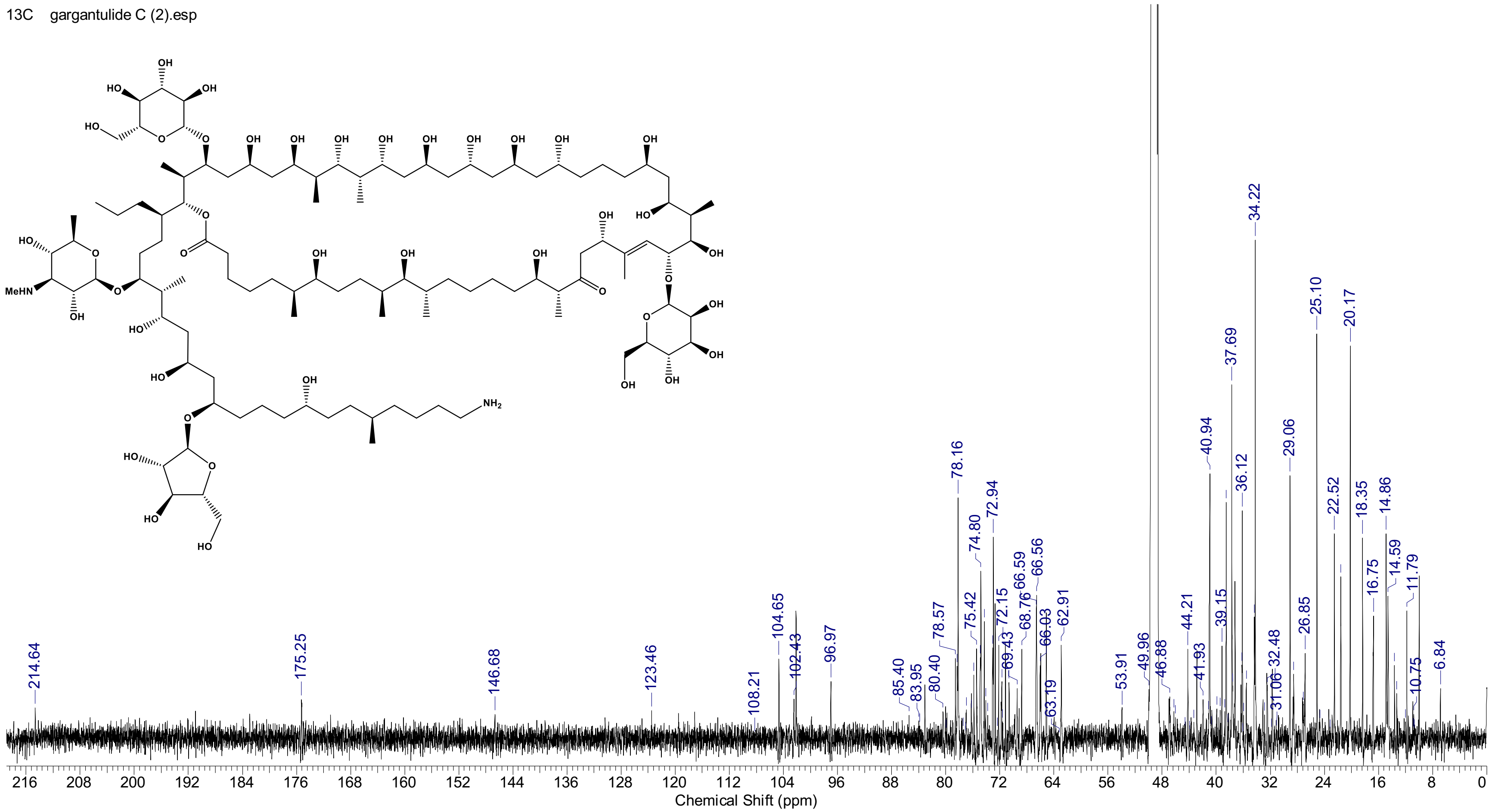


c) $^{13}$C NMR spectrum ($CD_3OD$, 125 MHz) of **2**

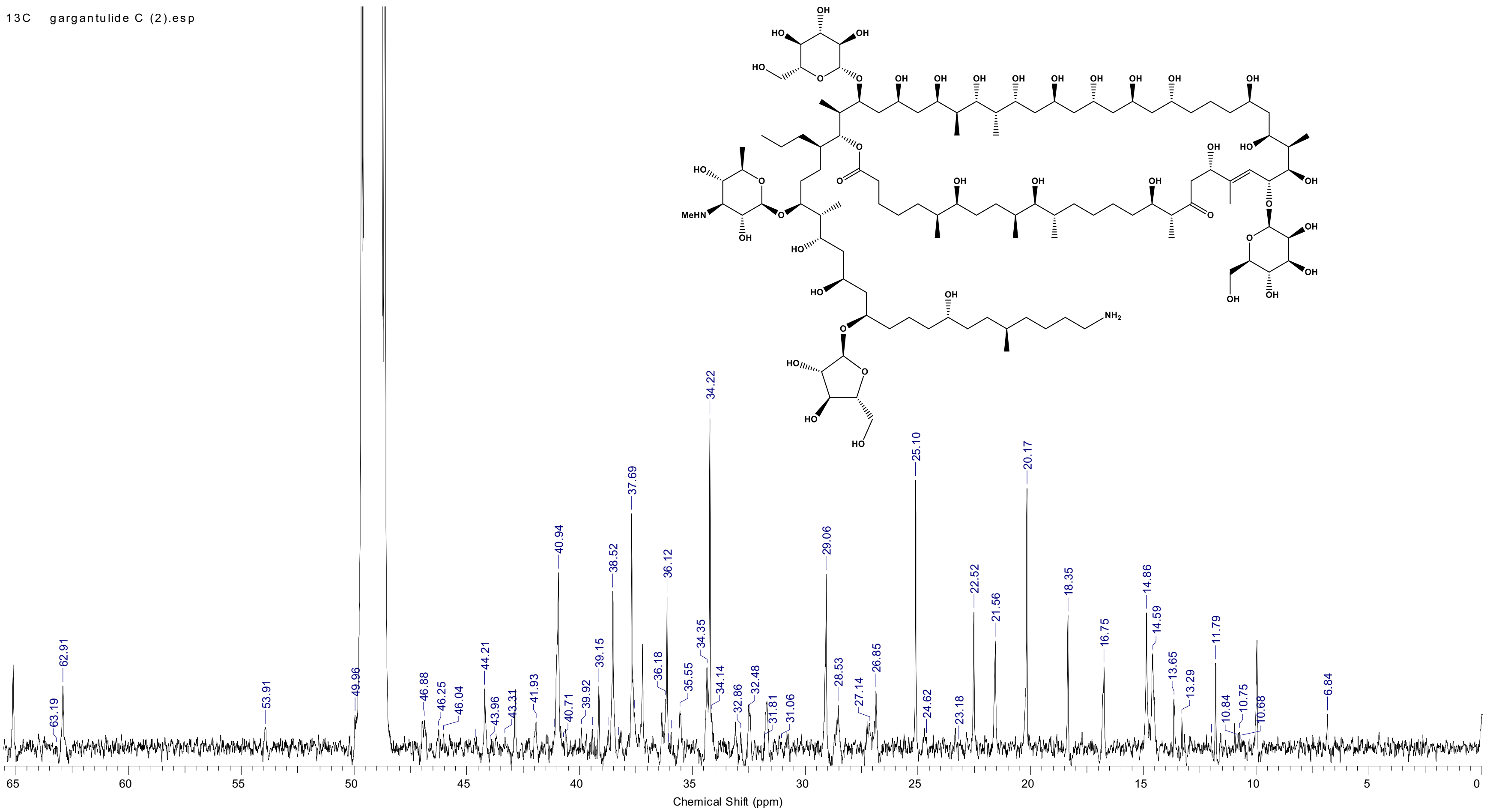


d) $^{13}$C NMR spectrum ($CD_3OD$, 125 MHz) of **2**. Expansion between 0 and 65 ppm

13C gargantulide C (2).esp

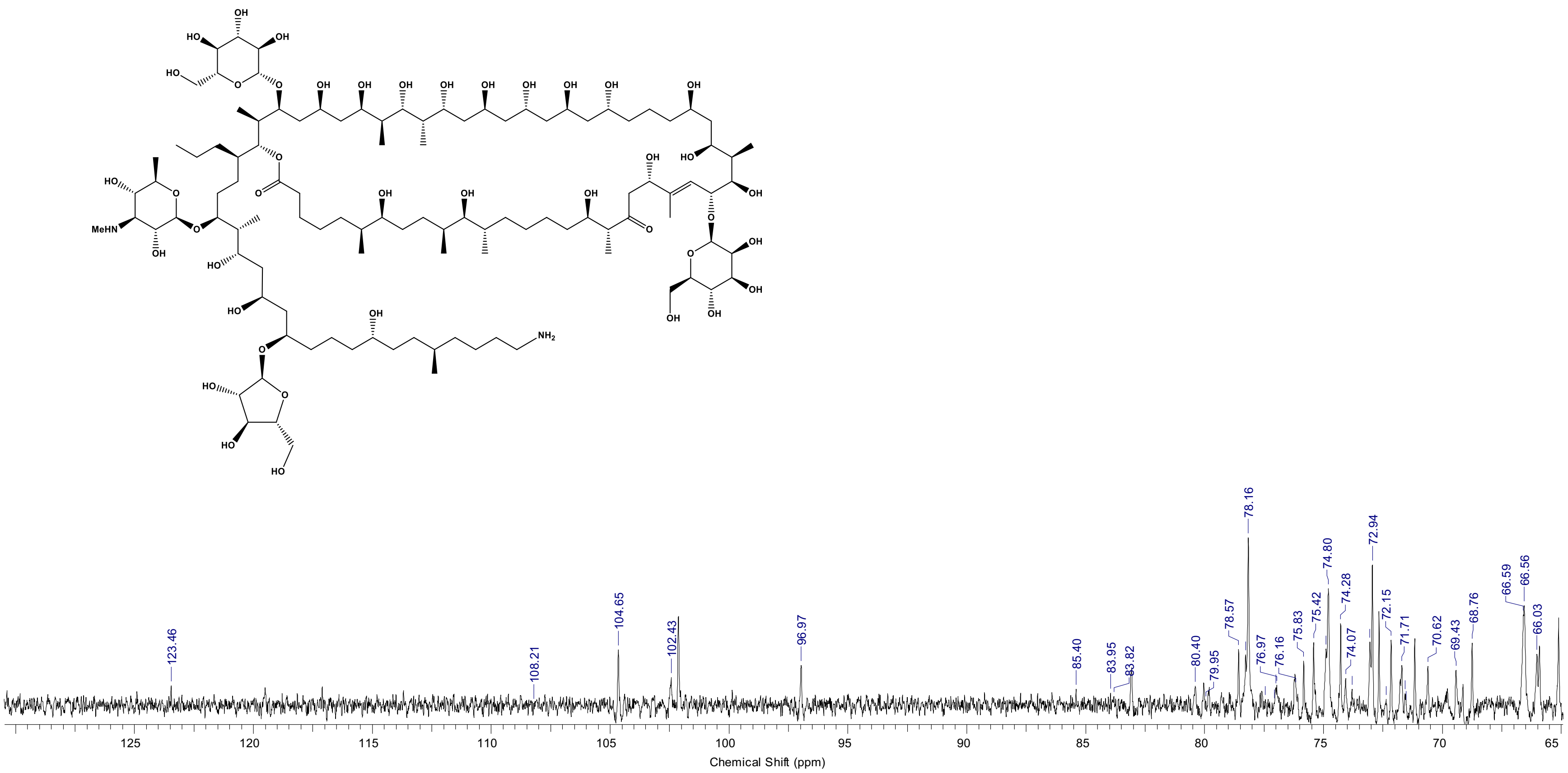


e) $^{13}C$ NMR spectrum ($CD_3OD$, 125 MHz) of **2**. Expansion between 65 and 130 ppm

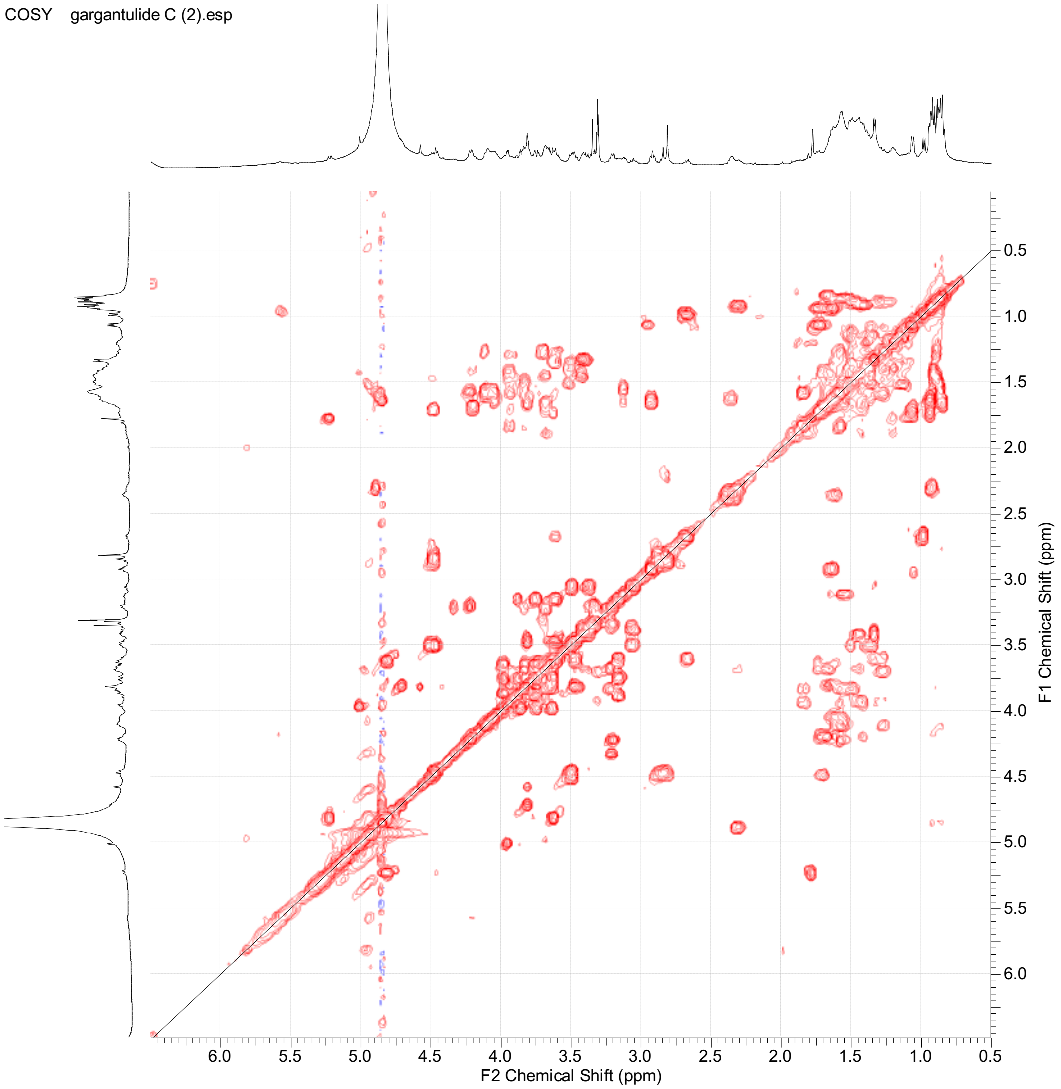


f) COSY spectrum of **2**

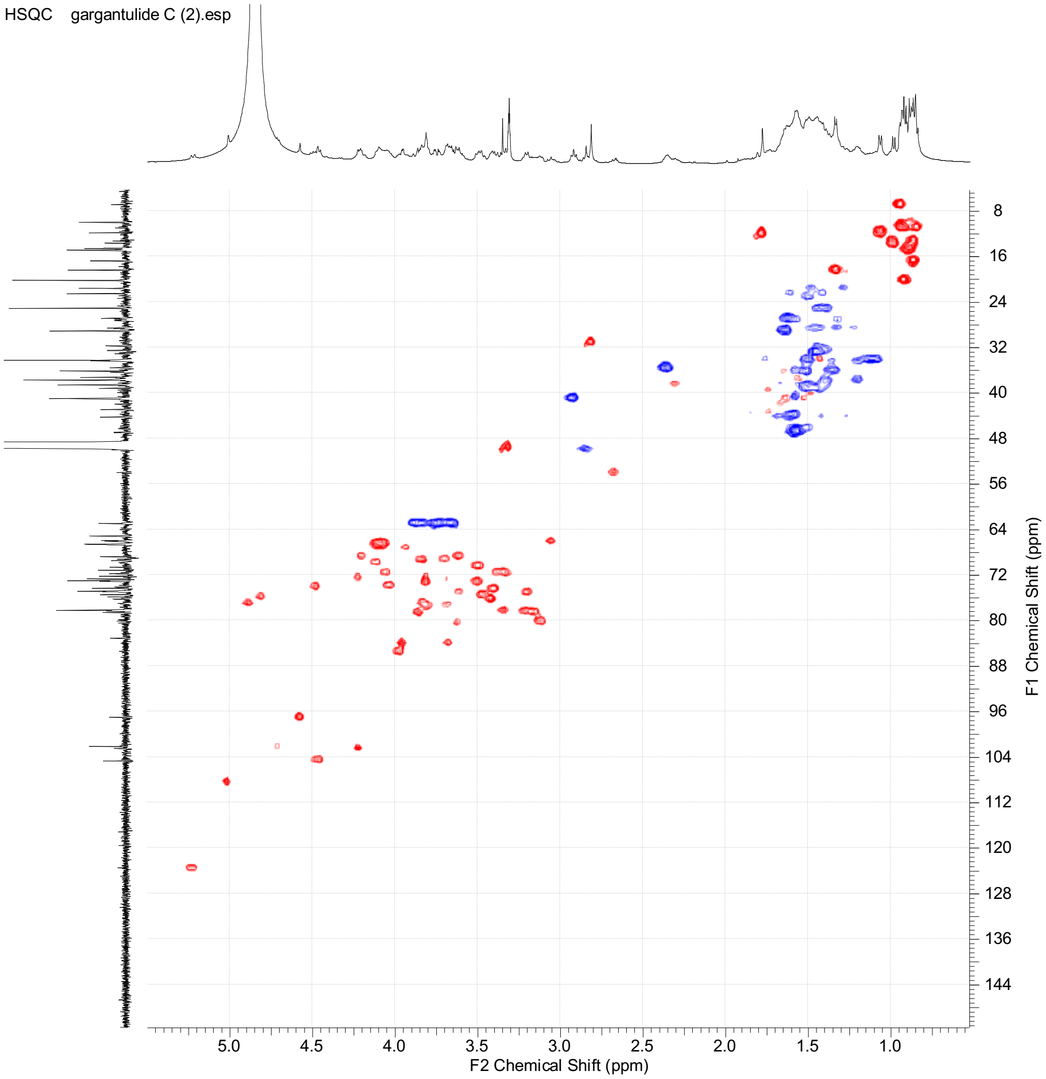


g) HSQC spectrum of **2**

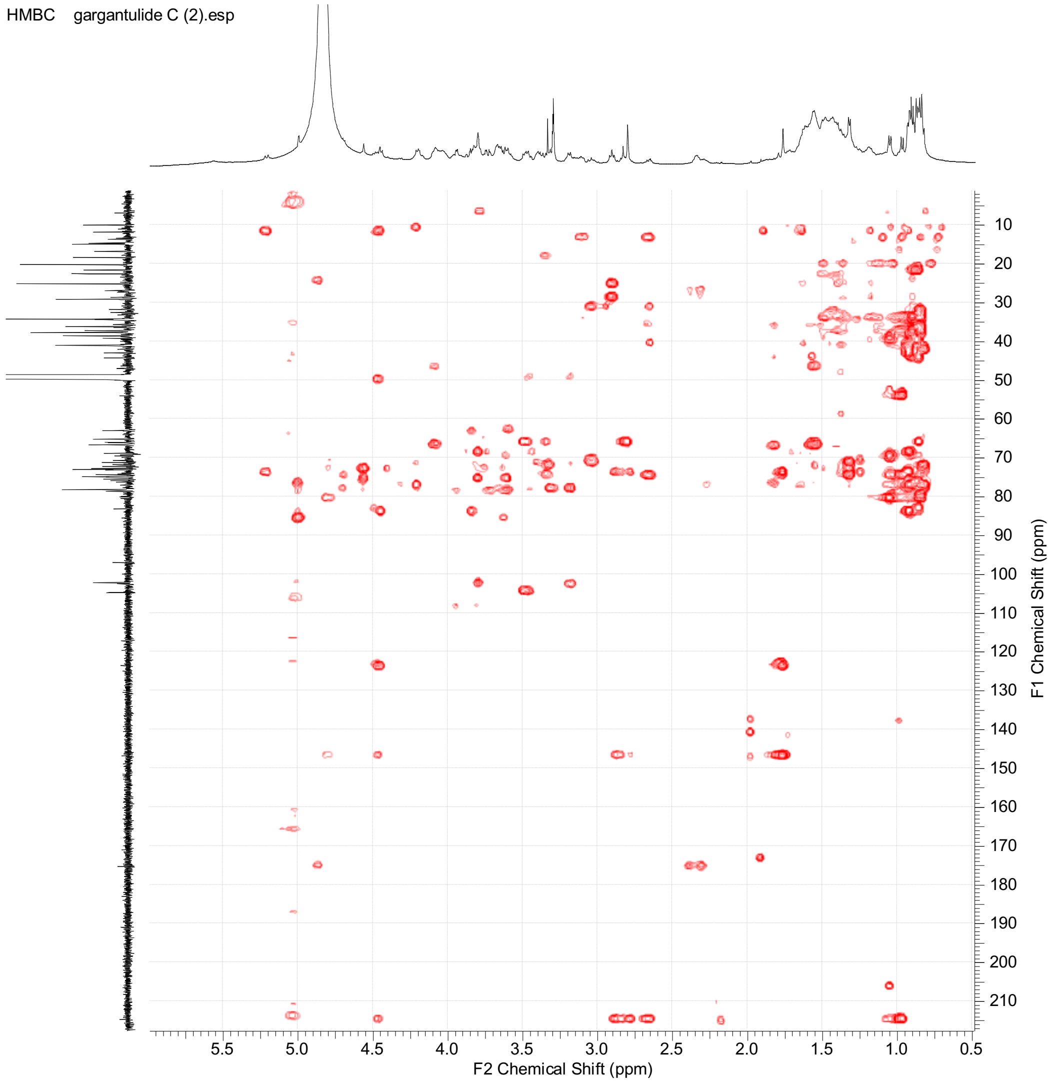


h) HMBC spectrum of **2**

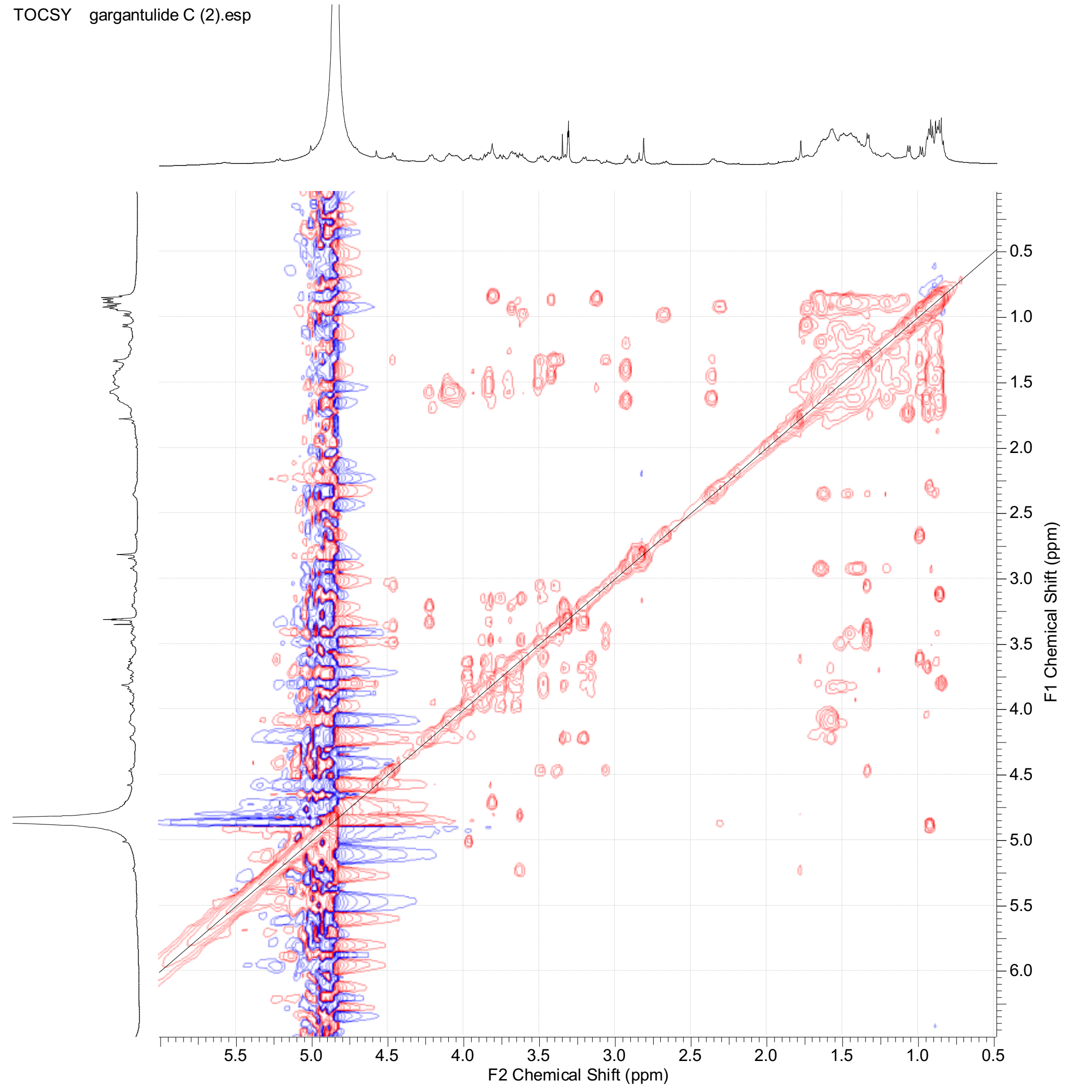


i) TOCSY spectrum of **2**

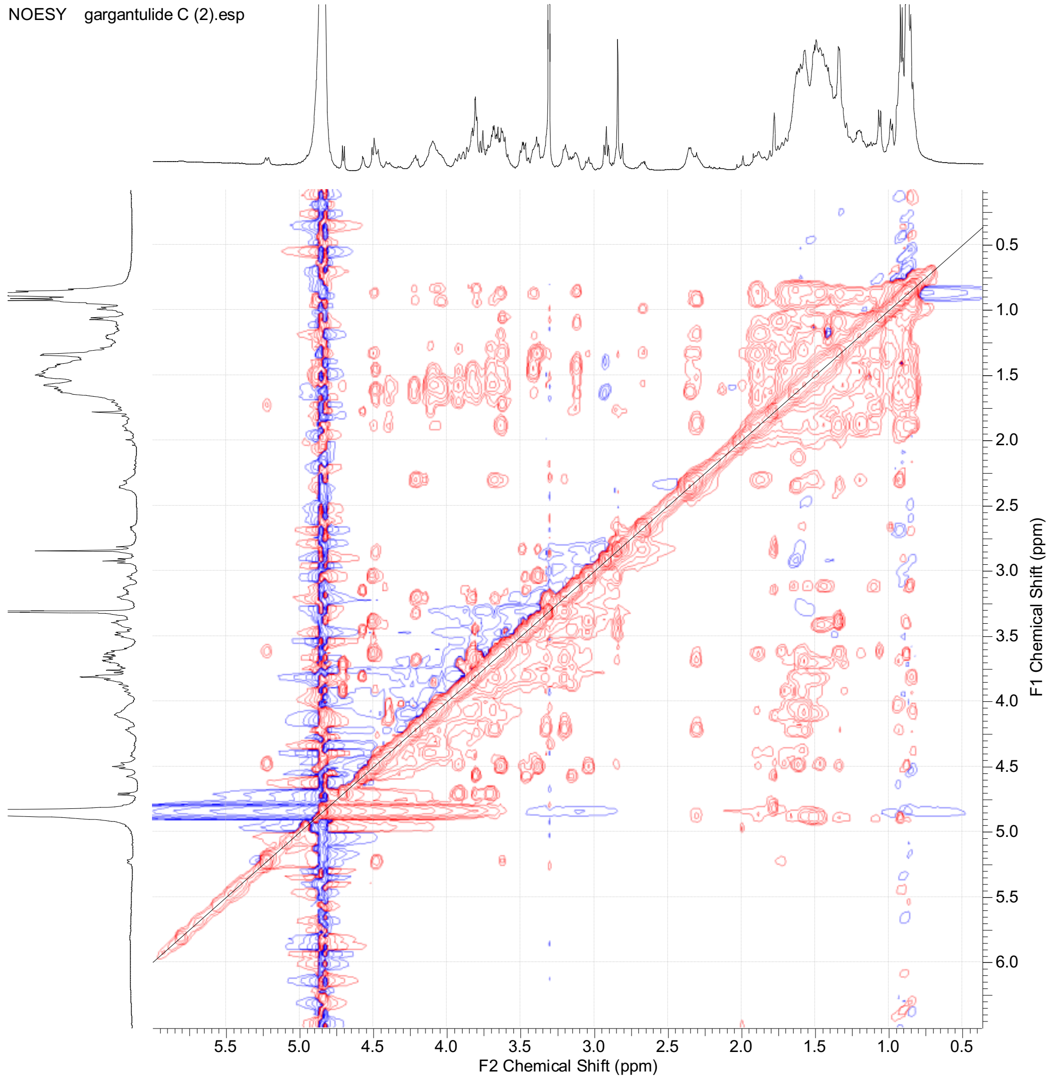


j) NOESY spectrum of **2**

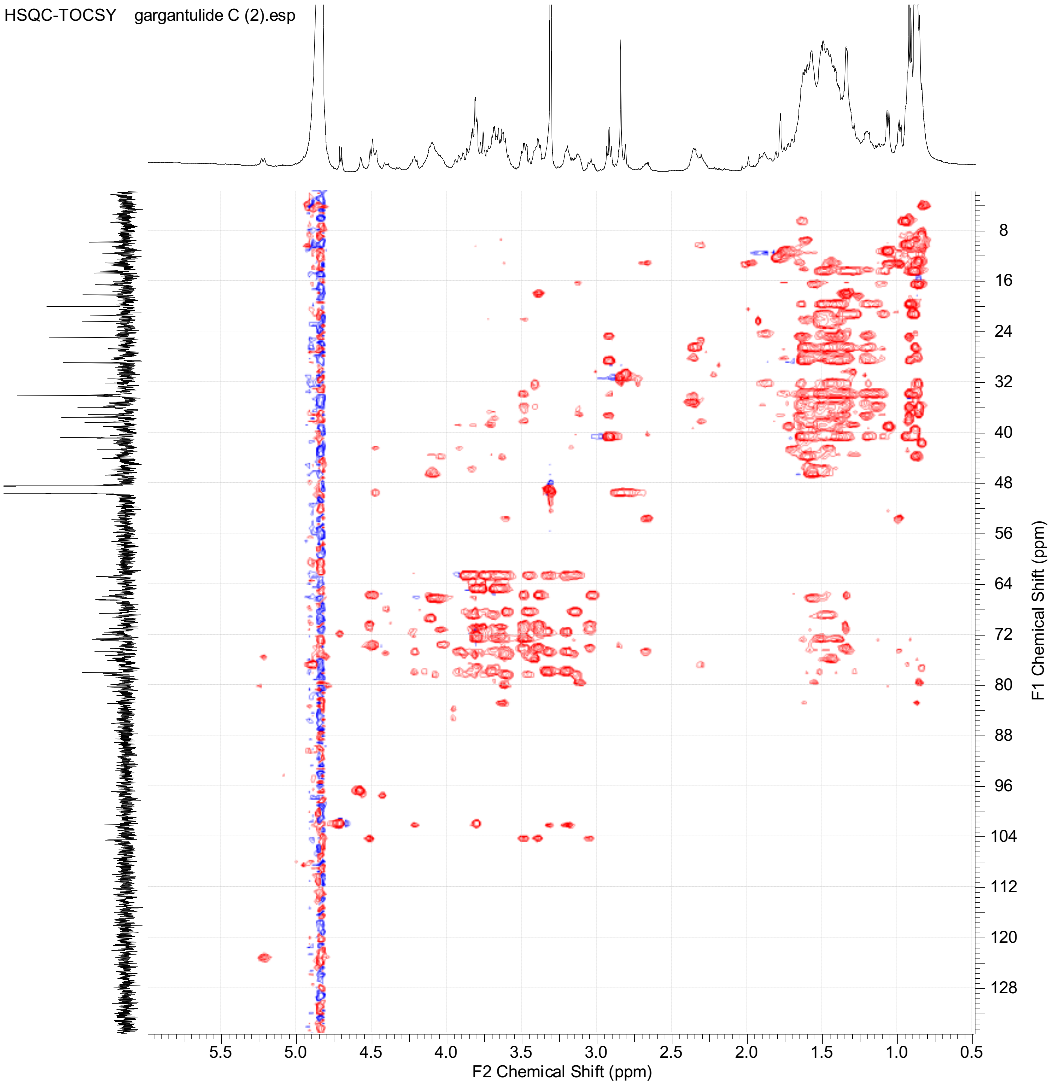


k) HSQC-TOCSY spectrum of **2**

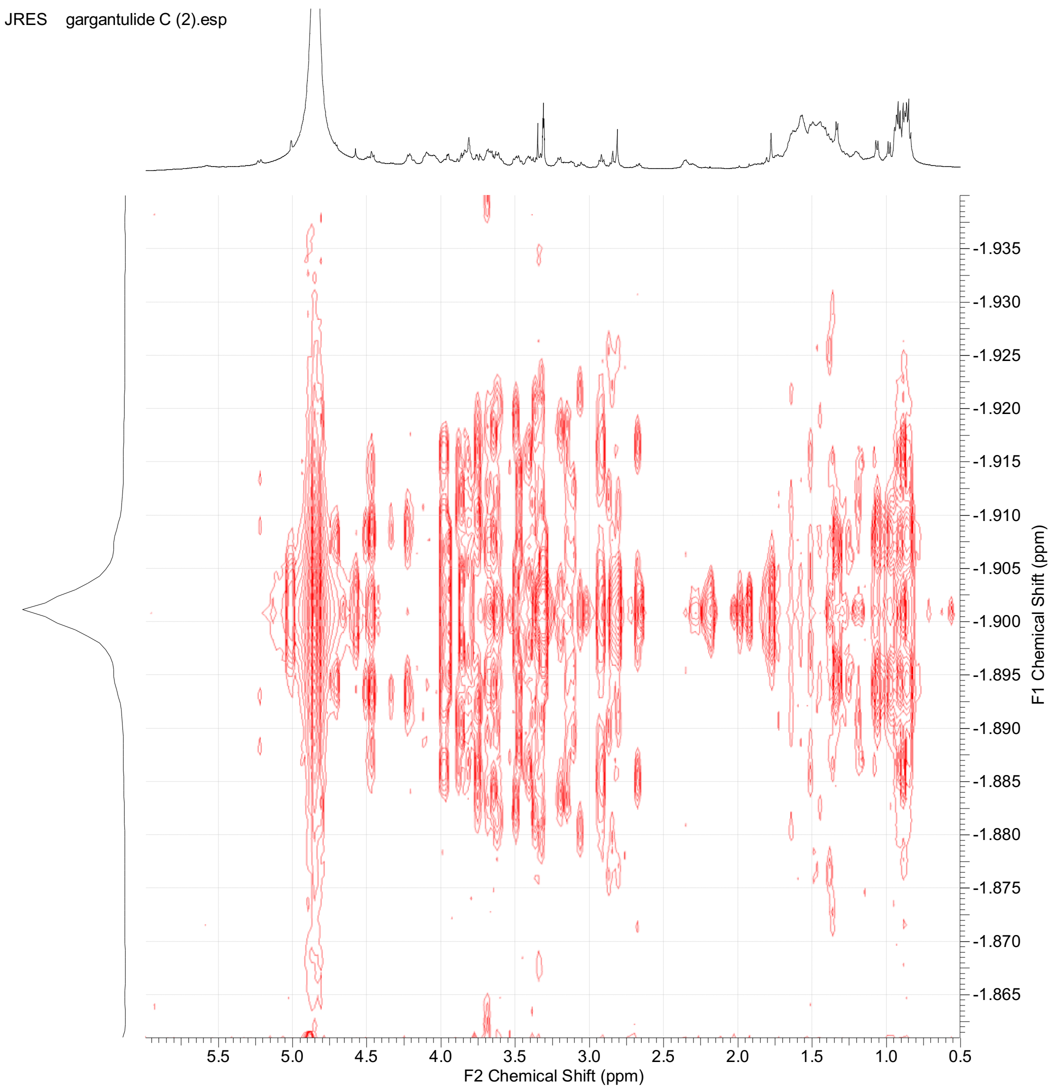


l) JRES spectrum of **2**

**Figure S6.** Key COSY, TOCSY, HSQC-TOCSY and HMBC correlations observed for **2**

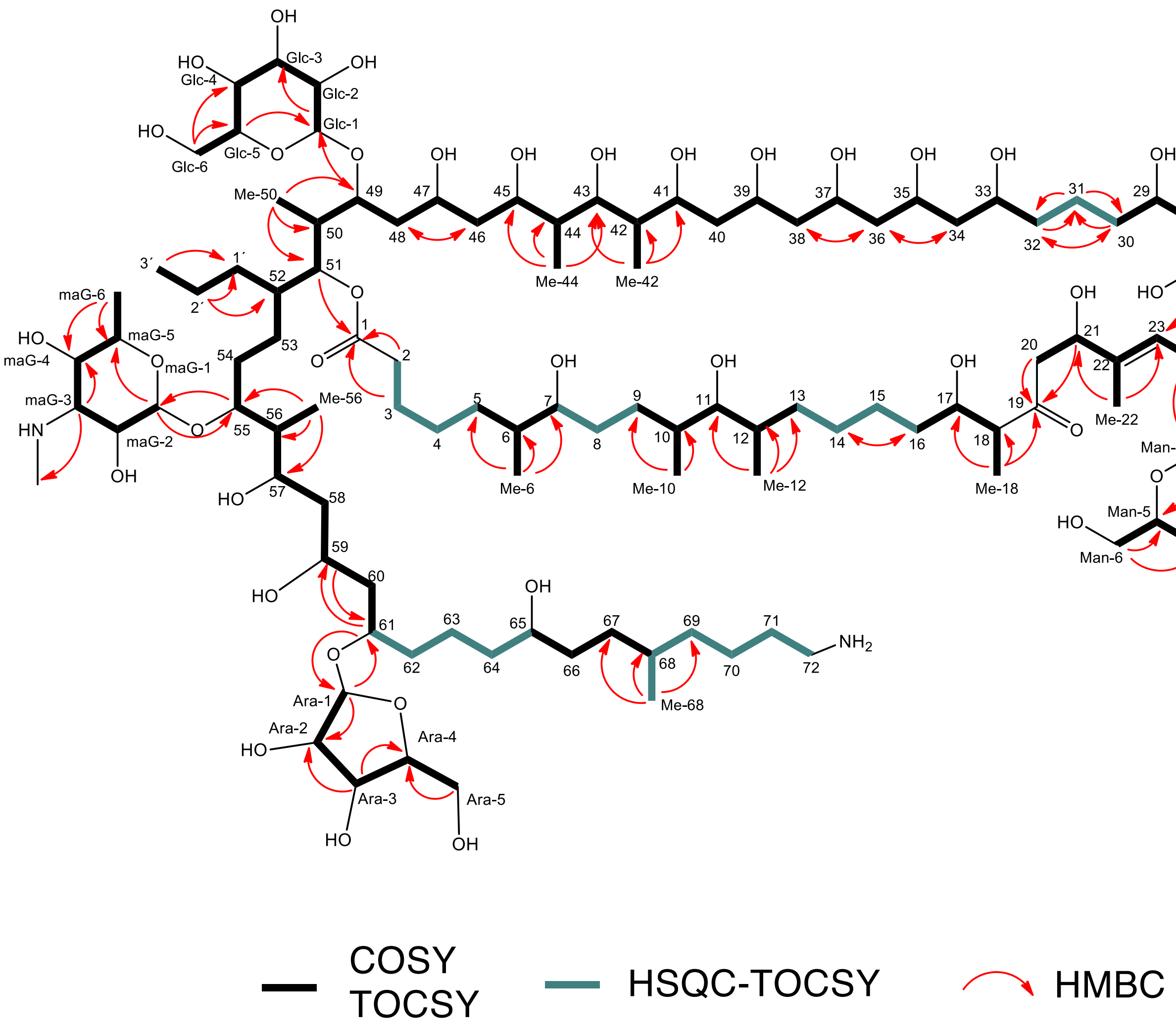

**Figure S7.** LC-UV-HRMS chromatogram (UV 210 nm: pink trace; MS+: blue trace) of the acetone crude extract of a parallel micro-scale fermentation showing the co-detection of gargantulides A, B and C. HRESIMS(+)-TOF spectra (ISCID 0 eV) of gargantulides A (triply and doubly-charged adducts) B and C (doubly-charged ions)

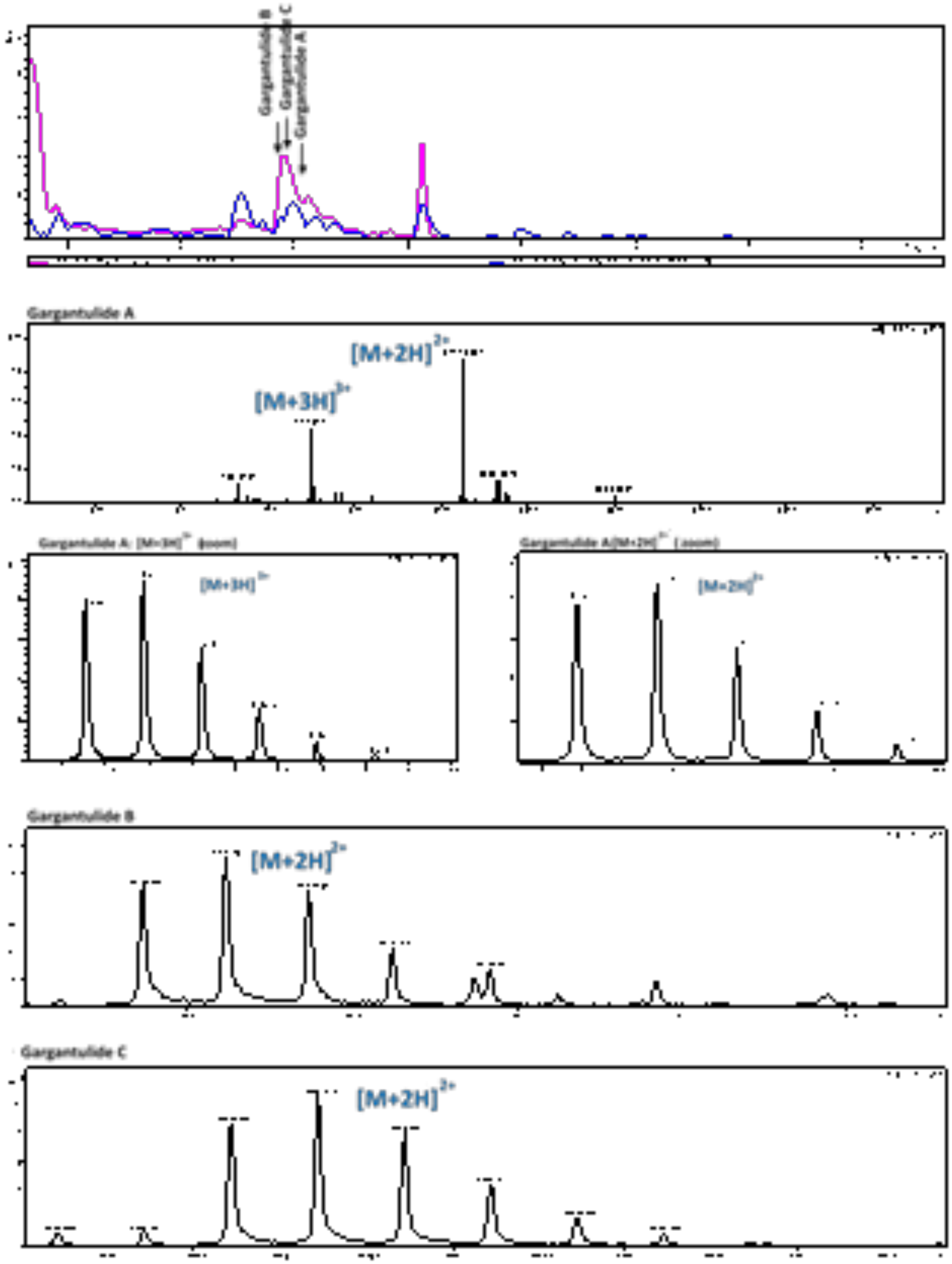

**Table S2.** antiSMASH results, bacterial version, relaxed mode. All types of BGCs are listed as detected by antiSMASH

| Region number | Type | Approx. size, kbp | Most similar known cluster | Remarks |
|---|---|---|---|---|
| **1** | PKS-like | 40,855 | No hits | |
| **2** | NRPS | 44,211 | Daptomycin (4%) | |
| **3** | Siderophore | 11,832 | Macrotetrolide (33%) | |
| **4** | RIPP-like | 9,971 | Enduracidin (4%) | |
| **5** | LAP, thiopeptide | 27,977 | No hits | |
| **6** | Terpene | 21,678 | Geosmin (100%) | |
| **7** | NRPS-like, terpene | 42,088 | Isorenieratene (42%) | Possibly two clusters |
| **8** | T1PKS | 45,457 | EDHA (33%) | |
| **9** | Terpene | 19,827 | Lipopetide 8D1-1 (6%) | |
| **10** | T1PKS | 45,036 | Sch-47554/Sch-47555 (3%) | |
| **11** | T1PKS, oligosaccharide | 216,516 | Funisamine (30%) | Putative cluster, coding for gargantulide compounds |
| **12** | LAP, ladderane, thioamide-NRP, NRPS | 76,620 | Ishigamide (72%) | Possibly several clusters |
| **13** | Other, thiopeptide, terpene, lantipeptide clas II | 72,243 | 2-methylisoborneol (100%) | |
| **14** | NAPAA | 31,640 | Himastatin (8%) | |
| **15** | NRPS | 62,803 | Albachelin (70%) | |
| **16** | CDPS | 20,683 | No hits | |
| **17** | PKS-like | 40,596 | No hits | |
| **18** | Lantipeptide class I | 25,334 | No hits | |
| **19** | Lantipeptide class III | 22,579 | Ery-9 / Ery-6 / Ery-8 / Ery-7 / Ery-5 / Ery-4 / Ery-3 (100%) | |
| **20** | Arylpolyene | 40,340 | Ibomycin (5%) | |
| **21** | Ectoine | 8,532 | Ectoine (100%) | |
| **22** | NRPS, T1PKS | 73,041 | Ecumicin (10%) | |
| **23** | thiopeptide, LAP | 30,258 | No hits | |
| **24** | redox-cofactor | 22,025 | Lankacidin C (20%) | |

| | | | | |
|---|---|---|---|---|
| **25** | cyanobactin, PKS-like, NRPS, T1PKS, lantipeptide class V, linaridin | 85,858 | prenylagaramide B / prenylagaramide C (8%) | |
| **26** | Lantipeptide class V | 40,587 | Arginomycin (13%) | |
| **27** | Linaridin | 20,557 | Legonaridin (25%) | |
| **28** | Lantipeptide class V | 41,924 | Ashimides (8%) | |
| **29** | Lantipeptide class II | 23,053 | No hits | |
| **30** | NAPAA | 32,569 | No hits | |
| **31** | hglE-KS, T1PKS | 49,901 | Rifamorpholine A/B/C/D/E (4%) | |
| **32** | NRPS | 57,957 | Telomycin (5%) | |
| **33** | NRPS, ectoine, T1PKS | 80,296 | Kosinostatin (16%) | |
| **34** | NRPS, NRPS-like | 74,191 | WS9326 (10%) | |
| **35** | NAPAA | 32,384 | No hits | |
| **36** | Terpene | 21,115 | SF2575 (6%) | |
| **37** | Ladderane, NRPS, T1PKS | 173,101 | Spiramycin (22%) | Possibly three clusters |
| **38** | NRPS, arylpolyene | 130,298 | Kedarcidin (14%) | |
| **39** | T1PKS | 67,639 | Meilingmycin (5%) | |
| **40** | Lassopeptide | 24,502 | No hits | |
| **41** | ladderane,NRPS,NRPS-like,transAT-PKS,T1PKS,NAPAA | 245,836 | Incednine (13%) | Possibly several clusters |
| **42** | NRPS, RIPP-like | 49,672 | Capreomycin IA/IB (6%) | |
| **43** | Furan | 20,998 | No hits | |
| **44** | NRPS, thioamitides | 80,729 | Mannopeptimycin (44%) | |
| **45** | Betalactone | 25,210 | No hits | |

**Table S3.** Putative functions of genes in *gar* BGC

| Protein (NCBI accession) | a.a. | Proposed function | BLAST hit protein | Query coverage/Per. identity/ E value | Accession number of BLAST hit |
|---|---|---|---|---|---|
| **GarR1 (HUW46_01962)** | 938 | Putative MalT family HTH-containing regulator | Helix-turn-helix transcriptional regulator (*Kutzneria albida*) | 99% / 49.58% / 0.0 | WP_025354824.1 |
| **GarR2 (HUW46_01961)** | 414 | Putative sensor histidine kinase | Sensor histidine kinase (*Asanoa ferruginea*) | 95% / 51.23% / 3e-110 | WP_116067603.1 |
| **GarT1 (HUW46_01960)** | 240 | Putative ABC-type multidrug transport system, ATPase component | ATP-binding cassette domain-containing protein (*Micromonospora pattaloongensis*) | 96% / 51.95% / 1e-62 | WP_091558576.1 |
| **GarA (HUW46_01959)** | 467 | Acyl-CoA ligase | Acyl-CoA ligase (*Kibdelosporangium aridum*) | 99% / 73.18% /0.0 | WP_084424151.1 |
| **Orf1 (HUW46_01958)** | 180 | PH domain-containing protein | PH domain-containing protein (Pseudonocardiaceae bacterium) | 91% /59.76%/ 3e-56 | MPZ66135.1 |
| **Orf2 (HUW46_01957)** | 524 | PH domain-containing protein | PH domain-containing protein (*Kutzneria albida*) | 95% /52.08 / 6e-169 | WP_025356057.1 |
| **GarR3 (HUW46_01956)** | 228 | TetR family transcriptional regulator | TetR/AcrR family transcriptional regulator [*Kutzneria albida*] | 81% / 48.92% / 4e- 55 | WP_025359812.1 |
| **GarT2 (HUW46_01955)** | 615 | Putative ABC-type multidrug transport system, ATPase component | ATP-binding cassette domain-containing protein [*Amycolatopsis* sp. EGI 650086] | 95% / 63.27 / 0.0 | WP_158888982.1 |
| **GarG1 (HUW46_01954)** | 396 | Putative glycosyltransferase of MGT family | glycosyltransferase MGT family protein [*Mycolicibacterium thermoresistibile*] | 98% / 57.11% / 6e-142 | WP_003926533.1 |
| **Orf3 (HUW46_01953)** | 84 | Hypothetical protein | hypothetical protein [*Sciscionella* sp. SE31] | 91% / 53.25% / 7e-17 | WP_031470089.1 |
| **GarG2 (HUW46_01952)** | 457 | Putative glycosyltransferase of MGT family | glycosyltransferase family 1 protein [*Amycolatopsis* sp. SID8362] | 92% / 43.88% / 5e-115 | WP_160698909.1 |

| | | | | | |
|---|---|---|---|---|---|
| **GarG3 (HUW46_01951)** | 421 | Putative glycosyltransferase of MGT family | glycosyltransferase family 1 protein [*Allokutzneria albata*] | 97% / 46.12% / 1e-115 | WP_030432743.1 |
| **GarG4 (HUW46_01950)** | 419 | Putative glycosyltransferase of MGT family | glycosyltransferase family 1 protein [*Allokutzneria albata*] | 98% / 43.06% / 1e-104 | WP_030432743.1 |
| **GarP1 (HUW46_01949)** | 4568 | Modular polyketide synthase | type I polyketide synthase [*Actinophytocola oryzae*] | 97% / 55.09% / 0.0 | WP_133901401.1 |
| **GarP2 (HUW46_01948)** | 9791 | Modular polyketide synthase | modular polyketide synthase [*Streptomyces* sp. RK95-74] | 96% / 56.64% /0.0 | BAW35608.1 |
| **GarP3 (HUW46_01947)** | 5411 | Modular polyketide synthase | PKS I [*Kutzneria albida* DSM 43870] | 99% / 56.34% / 0.0 | AHH99926.1 |
| **GarP4 (HUW46_01946)** | 6168 | Modular polyketide synthase | type I polyketide synthase [*Streptomyces* sp. LamerLS-31b] | 98% / 55.92% / 0.0 | WP_093893008.1 |
| **GarP5 (HUW46_01945)** | 8075 | Modular polyketide synthase | beta-ketoacyl synthase [*Streptomyces hygroscopicus*] | 99% / 54.13% / 0.0 | AQW47576.1 |
| **GarP6 (HUW46_01944)** | 4542 | Modular polyketide synthase | type I polyketide synthase [*Streptomyces cinnamoneus*] | 99% / 62.72% / 0.0 | WP_099199035.1 |
| **GarP7 (HUW46_01943)** | 5347 | Modular polyketide synthase | type I polyketide synthase [*Amycolatopsis orientalis*] | 100% / 54.65% / 0.0 | WP_051173827.1 |
| **GarP8 (HUW46_01942)** | 3617 | Modular polyketide synthase | type I polyketide synthase [*Streptomyces cinnamoneus*] | 99% / 59.56% / 0.0 | WP_099199033.1 |
| **GarP9 (HUW46_01941)** | 5662 | Modular polyketide synthase | PKS I [*Kutzneria albida* DSM 43870] | 99% / 58.65% / 0.0 | AHH99926.1 |
| **GarP10 (HUW46_01940)** | 5910 | Modular polyketide synthase | type I polyketide synthase [*Streptomyces cinnamoneus*] | 95% / 61.75% / 0.0 | WP_104650952.1 |
| **GarB (HUW46_01939)** | 314 | AT domain containing protein | ACP S-malonyltransferase [unclassified *Kitasatospora*] | 96% / 53.77% / 5e-111 | WP_057230119.1 |
| **GarC (HUW46_01938)** | 421 | Putative cytochrome p450 | cytochrome P450 [*Streptomyces cinnamoneus*] | 99% / 56.87 / 1e-165 | WP_099199030.1 |

| | | | | | |
|---|---|---|---|---|---|
| **GarR4 (HUW46_01937)** | 222 | Putative response regulator | response regulator [Propionibacteriales bacterium] | 92% / 69.42% / 1e-94 | MPZ63410.1 |
| **GarR5 (HUW46_01936)** | 452 | Sensor histidine kinase | sensor histidine kinase [Propionibacteriales bacterium] | 87% / 51.87% / 1e-103 | MPZ96507.1 |
| **GarD (HUW46_01935)** | 156 | Putative TDP-4-keto-6-deoxy-D-glucose 3,4-isomerase | WxcM-like domain-containing protein [*Amycolatopsis antarctica*] | 88% / 74.64% / 4e-71 | WP_094863117.1 |
| **GarE (HUW46_01934)** | 367 | Putative aminotransferase | DegT/DnrJ/EryC1/StrS family aminotransferase [Actinoalloteichus] | 100% / 68.39% / 0.0 | WP_075741202.1 |
| **GarG5 (HUW46_01933)** | 513 | Putative glycosyltransferase | glycosyltransferase, family 39 [*Streptomyces cinnamoneus*] | 100% / 60.23% / 0.0 | WP_099199029.1 |
| **GarF (HUW46_01932)** | 450 | Putative crotonyl-CoA reductase | crotonyl-CoA carboxylase/reductase [*Actinomadura macra*] | 99% / 79.15% / 0.0 | WP_067467652.1 |
| **GarH (HUW46_01931)** | 338 | Putative *fabH* | ketoacyl-ACP synthase III [*Micromonospora pisi*] | 97% / 77.2 %/ 2e-175 | WP_121159999.1 |
| **GarI (HUW46_01930)** | 291 | Putative 3-hydroxybutyryl-CoA dehydrogenase | 3-hydroxybutyryl-CoA dehydrogenase [*Micromonospora pisi*] | 92% / 74.72% / 3e-138 | WP_121160000.1 |
| **GarR6 (HUW46_01929)** | 956 | Putative HTH transcriptional regulator | helix-turn-helix transcriptional regulator [*Kutzneria albida*] | 97% / 49.89% / 0.0 | WP_025354824.1 |
| **Orf4 (HUW46_01928)** | 369 | Putative IS4-like element | IS4-like element ISMfl1 family transposase [Mycobacteriaceae] | 98% / 36.22% / 3e-52 | WP_011891513.1 |
| **Orf5 (HUW46_01927)** | 73 | Hypothetical protein | hypothetical protein [*Streptomyces kasugaensis*] | 100% / 60.27% / | WP_131126254.1 |
| **GarR7 (HUW46_01926)** | 222 | DNA-binding response regulator | DNA-binding response regulator, NarL/FixJ family, contains REC and HTH domains [*Sinosporangium album*] | 98% / 62.39 / 1e-86 | SDG85138.1 |
| **Orf6 (HUW46_01925)** | 370 | Putative IS256 family element | IS256 family transposase [*Actinopolymorpha alba*] | 66% / 30.26% / 5e-36 | WP_040420563.1 |

**Table S4.** Prediction of activity and stereochemistry of AT, KR, ER and DH domains from bioinformatics analysis. Stereochemical outcomes for gargantulides B and C.

| Protein | Module | Substrate of AT domain | KR domain | DH domain | ER domain | Predicted stereochemistry (Keatinge-Clay)[a] | Predicted stereochemistry (fully assembled linear polyketides)[b] | Stereochemical outcome for gargantulides B and C[c] |
|---|---|---|---|---|---|---|---|---|
| GarP1 | 1 | Mmal-CoA | B1 | Active | R | "*R*" (C-68) | *R* (C-68) | *R* (C-68) |
| | 2 | Mal-CoA | B1 | Active | R | - | - | - |
| GarP2 | 3 | Mal-CoA | B1 | - | - | "*R*" (C-65) | *S* (C-65) | *S* (C-65) |
| | 4 | Mal-CoA | B1 | Active | S | - | - | - |
| | 5 | Mal-CoA | B1 | - | - | "*R*" (C-61) | *R* (C-61) | *R* (C-61) |
| | 6 | Mal-CoA | B1 | - | - | "*R*" (C-59) | *R* (C-59) | *S* (C-59)[d] |
| | 7 | Mmal-coA | A1 | - | - | "*R*,*S*" (C-56,57) | *S*,*S* (C-56,57) | *R*,*S* (C-56,57)[e] |
| | 8 | Mal-CoA | B1 | n.d. | - | "*R*" (C-55) | *S* (C-55) | *S* (C-55) |
| GarP3 | 9 | Pmal-CoA[f] | B1 | Active | R | "*R*" (C-52) | *R* (C-52) | *R* (C-52) |
| | 10 | Mmal-CoA | B1 | - | - | "*R*,*R*" (C-50,51) | *S*,*R* (C-50,51) | *S*,*R* (C-50,51) |
| | 11 | Mal-CoA | B1 | Inactive | - | "*R*" (C-49) | *S* (C-49) | *S* (C-49) |
| GarP4 | 12 | Mal-CoA | B1 | - | - | "*R*" (C-47) | *R* (C-47) | *S* (C-47)[g] |
| | 13 | Mmal-CoA | B2 | - | - | "*S*,*R*" (C-44,45) | *S*,*R* (C-44,45) | *S*,*R* (C-44,45) |
| | 14 | Mmal-CoA | A1 | - | - | "*R*,*S*" (C-42,43) | *S*,*R* (C-42,43) | *S*,*R* (C-42,43) |
| | 15 | Mal-CoA | A1 | - | - | "*S*" (C-41) | *R* (C-41) | *R* (C-41) |
| GarP5 | 16 | Mal-CoA | B1 | - | - | "*R*" (C-39) | *S* (C-39) | *S* (C-39) |
| | 17 | Mal-CoA | A1 | - | - | "*S*" (C-37) | *R* (C-37) | *R* (C-37) |
| | 18 | Mal-CoA | B1 | - | - | "*R*" (C-35) | *S* (C-35) | *S* (C-35) |
| | 19 | Mal-CoA | A1 | - | - | "*S*" (C-33) | *R* (C-33) | *R* (C-33) |
| | 20 | Mal-CoA | B1 | Active | S | - | - | - |
| GarP6 | 21 | Mal-CoA | B1 | - | - | "*R*" (C-29) | *R* (C-29) | *R* (C-29) |
| | 22 | Mmal-CoA | A1 | - | - | "*R*,*S*" (C-26,27) | *S*,*S* (C-26,27) | *R*,*S* (C-26,27)[h] |
| | 23 | Mal-CoA | B1 | - | - | "*R*" (C-25) | *S* (C-25) | *R* (C-25)[h] |
| GarP7 | 24 | Mmal-CoA | B1 | Active | - | "*E*" double bond | *E* double bond | *E* double bond |
| | 25 | Mal-CoA | B1 | - | - | "*R*" (C-21) | *S* (C-21) | *S* (C-21) |
| | 26 | Mmal-CoA | C1[i] | n.d. | n.d. | "*R*" (C-18) | *S* (C-18) | *R* (C-18)[i] |
| GarP8 | 27 | Mal-CoA | A1 | - | - | "*S*" (C-17) | *R* (C-17) | *R* (C-17) |
| | 28 | Mal-CoA | B1 | Active | S | - | - | - |
| GarP9 | 29 | Mmal-CoA | B1 | Active | S | "*S*" (C-12) | *S* (C-12) | *S* (C-12) |
| | 30 | Mmal-CoA | A1 | - | - | "*R*,*S*" (C-10,11) | *S*,*S* (C-10,11) | *S*,*S* (C-10,11) |
| | 31 | Mal-CoA | B1 | Active | S | - | - | - |
| GarP10 | 32 | Mmal-CoA | A1 | - | - | "*R*,*S*" (C-6,7) | *S*,*S* (C-6,7) | *S*,*S* (C-6,7) |
| | 33 | Mal-CoA | B1 | Active | S | - | - | - |
| | 34 | Mal-CoA | B1 | Active | S | - | - | - |

[a] Bioinformatics prediction for each separate module
[b] See Fig SX (fully assembled linear polyketides)
[c] Stereochemistry determined by a combination of NMR and bioinformatics gene cluster analysis for gargantulides B and C (and extrapolated to gargantulide A, revised in this work)
[d] The Cahn-Ingold-Prelog descriptor in gargantulides B and C for C-59 is inverted to *S* due to the glycosylation at C-61
[e] The Cahn-Ingold-Prelog descriptor in gargantulides A-C for C-56 is inverted to *R* due to the glycosylation at C-55
[f] Propyl malonyl-CoA established by NMR analysis
[g] The Cahn-Ingold-Prelog descriptor in gargantulides A-C for C-47 is inverted to *S* due to macrolactone cyclization and glycosylation at C-49.
[h] The Cahn-Ingold-Prelog descriptor in gargantulides A-C for both C-25 and C-26 is inverted to *R* due to the post-PKS hydroxylation at C-24
[i] Redox-inactive, methyl-epimerizing KR; (module 26)

**Figure S8.** Extracted amino acid sequence alignments of AT, KR, DH and ER domains

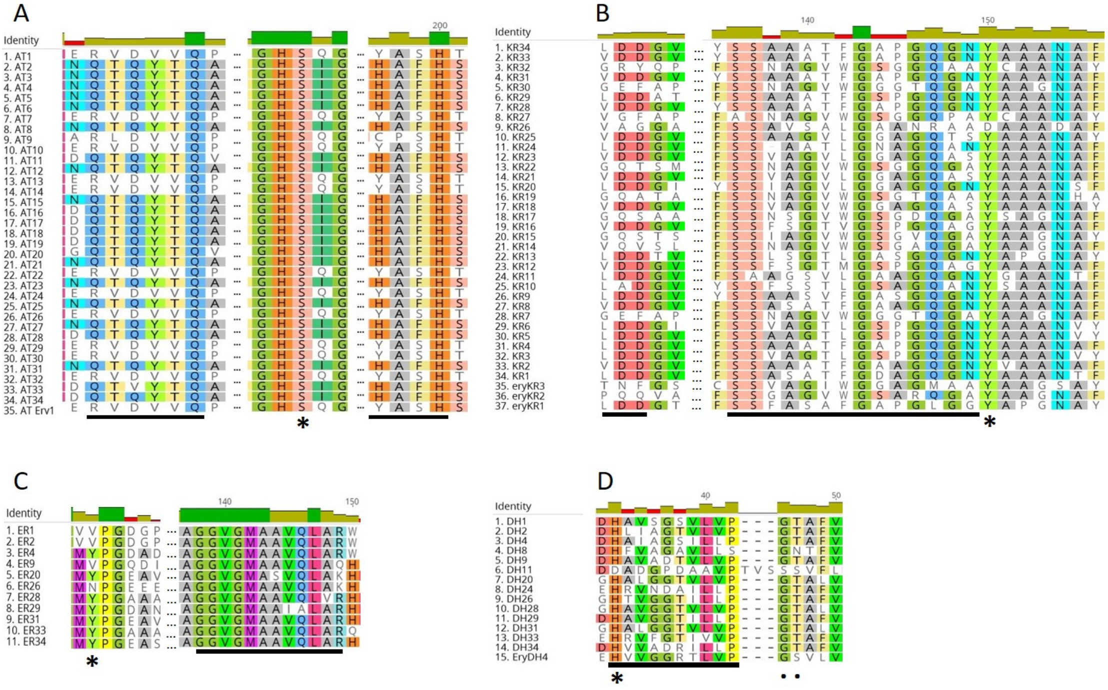


**A** - amino acid sequence alignment of AT domains. Characteristic residues for substrate recognition are underlined with black bars. The catalytic residue is identified with asterisk; **B** - amino acid sequence alignment of KR domains. Two motives, responsible for determination of A/B/C type of KRs are underlined with black bars; the catalytic tyrosine residue is identified with asterisk; **C** - amino acid sequence alignment of ER domains. The amino acid related to the stereochemistry of reduced residue is identified with an asterisk. NADPH binding site is identified with a black bar; **D** - amino acid sequence alignment of DH domains. The catalytic residue is identified with an asterisk. The conserved motif “HXXXGXXXXP” is identified with black bar. Two supporting catalytic residues are identified with filled black spots. The assignment of characteristic residues for substrate recognition and catalytic residues for ATs, KRs, DHs and ERs was done as described in previous works (3–6).

**Table S5.** Levels of identity and similarity of the putative *N*-Methyltransferase HUW46_03188 from CA-230715 with other known *N*-Methyltransferases.

| Protein | Microorganism | % Identity to HUW46_03188 | % Similarity to HUW46_03188 |
|---|---|---|---|
| TylM1 | *Streptomyces fradiae* | 43 | 61 |
| DesVI | *Streptomyces venezuelae* | 51 | 65 |
| SpnS | *Saccharopolyspora spinosa* | 42 | 58 |
| OssI | *Streptomyces ossamyceticus* | 43 | 59 |
| Orf1C | *Streptomyces ambofaciens* | 44 | 62 |
| Orf9c | *Streptomyces ambofaciens* | 44 | 57 |

**Table S6.** Identified genes encoding for putative glucose-1-phosphate thymidyltransferase and dTDP-glucose 4,6-dehydratase in the genome of CA-230715.

| CA-230715 ORF | Protein a. a. | Putative function | Best match accession number | % protein Identity / Similarity |
|---|---|---|---|---|
| HUW46_03193 | 293 | glucose-1-phosphate thymidylyltransferase | WP_020668504.1 | 87 / 93 |
| HUW46_02674 | 330 | dTDP-glucose 4,6-dehydratase | WP_144591396.1 | 88 / 92 |
| HUW46_09296 | 335 | dTDP-glucose 4,6-dehydratase | WP_125677294.1 | 72 / 83 |

**Figure S9.** Proposed biosynthetic pathway for the amino sugar 3,6-deoxy-3-methylamino D-glucose (maG)

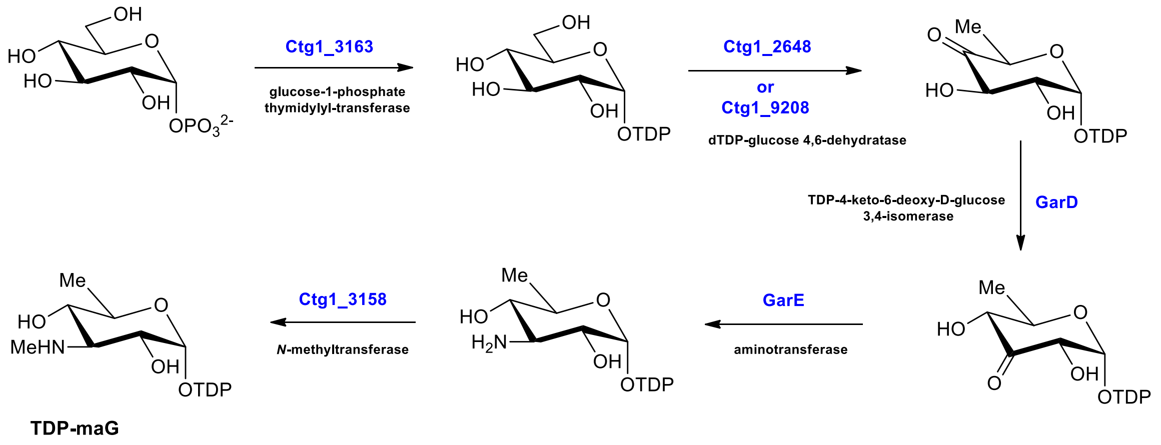

**Figure S10.** Bioinformatics prediction (*gar* BGC analysis) of the absolute configurations for the gargantulides polyketide aglycon (a). Comparison with the absolute configurations previously reported for gargantulide A (b) (13)

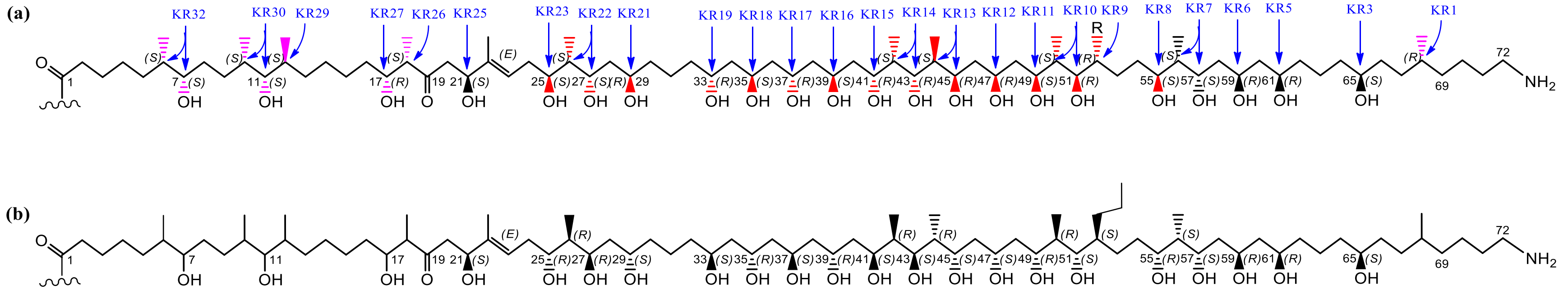


KR domains determining the stereochemical outcome are indicated in blue color. Chiral centers showing disagreement between the *in silico* prediction and the previously reported NMR-based structure for gargantulide A, are highlighted in red color. Undetermined chiral centers for gargantulide A in the original work, which have been now predicted by bioinformatic analysis, are highlighted in pink color.

**Figure S11.** Determination of the relative configuration of the C-55 to C-57 stereocluster for gargantulide B (**1**)

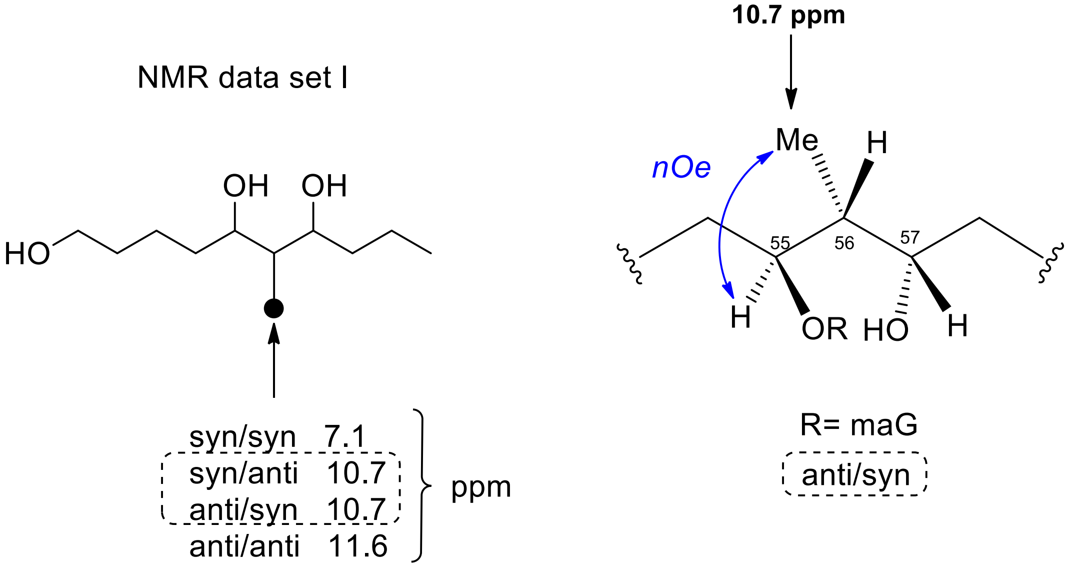


a) Application of Kishi´s NMR data set I for 2-methyl-1,3-diols (14) (left). Key NOESY correlations supporting the anti/syn configuration for C-55/C-56/C-57 (right)

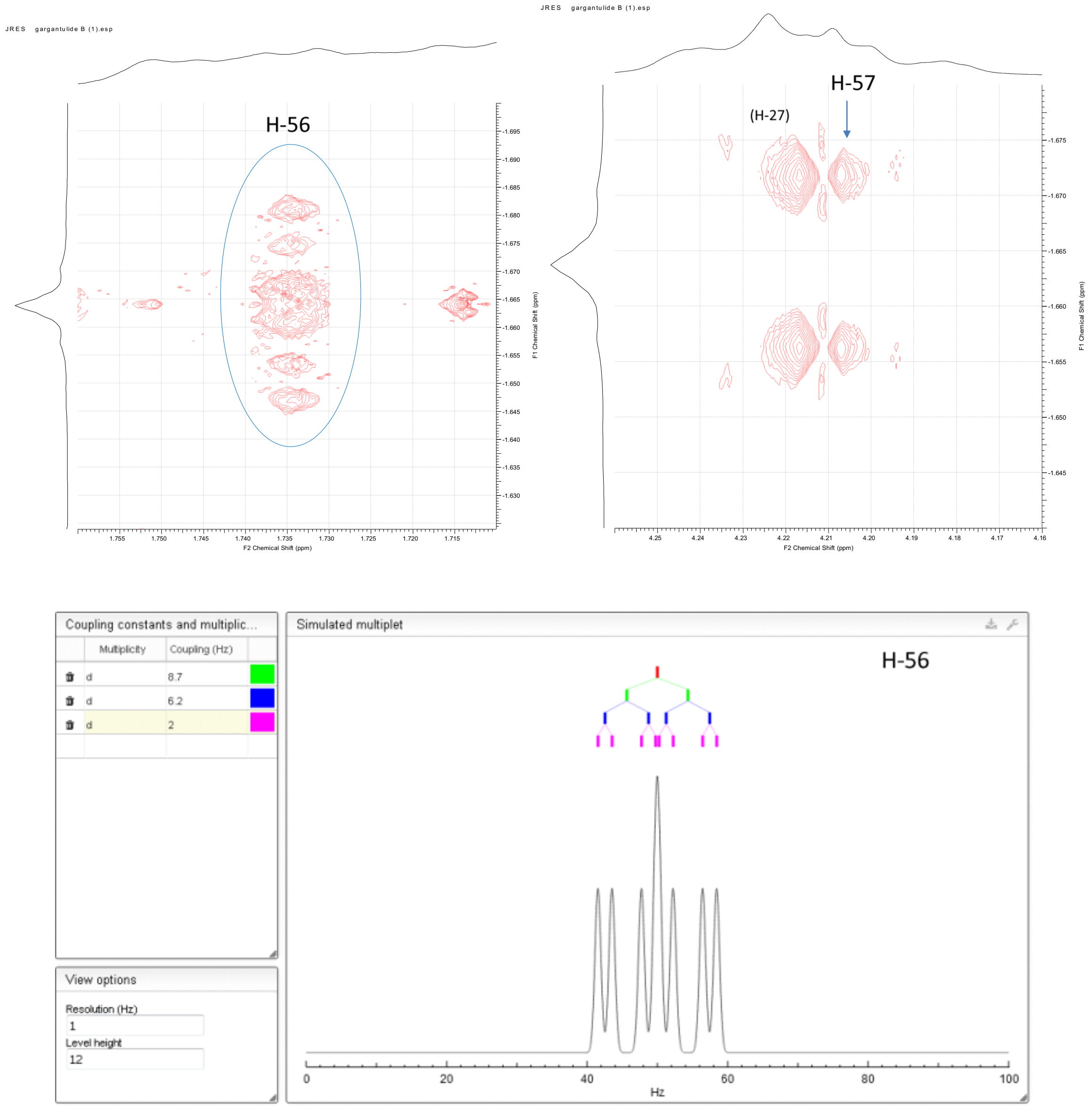


b) Expansions for the JRES spectrum of **1** showing the higher multiplicity of H-56 compared to that of H-57 (top). Multiplet simulation (15) for H-56 (applied values for $^1$H-$^1$H coupling constants are indicated) showing very good fitting with the multiplicity experimentally observed (bottom).

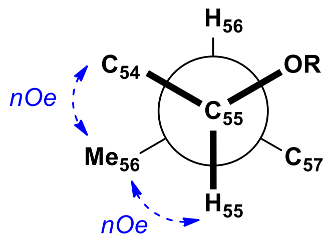


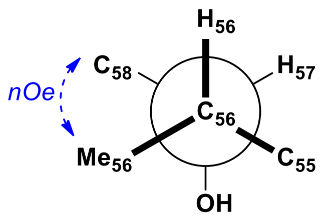


| | |
|---|---|
| $^3J_{H55\text{-}H56}$ = not measurable (overlappings) | $^3J_{H55\text{-}H56}$ = not measurable (overlappings) |
| $^3J_{C54\text{-}H56}$ = ca. small | $^3J_{C55\text{-}H57}$ = ca. small |
| $^3J_{C57\text{-}H55}$ = ca. small | $^3J_{C58\text{-}H56}$ = ca. small |
| $^3J_{Me56\text{-}H55}$ = ca. small | $^3J_{Me56\text{-}H57}$ = ca. large |

c) Qualitative $^3J_{C,H}$-based configuration analysis of the C-55 to C-57 stereocluster. Key NOESY correlations are also indicated (NOESY correlation Me-56/H-58 partially overlapped with other *nOe* cross-peaks).

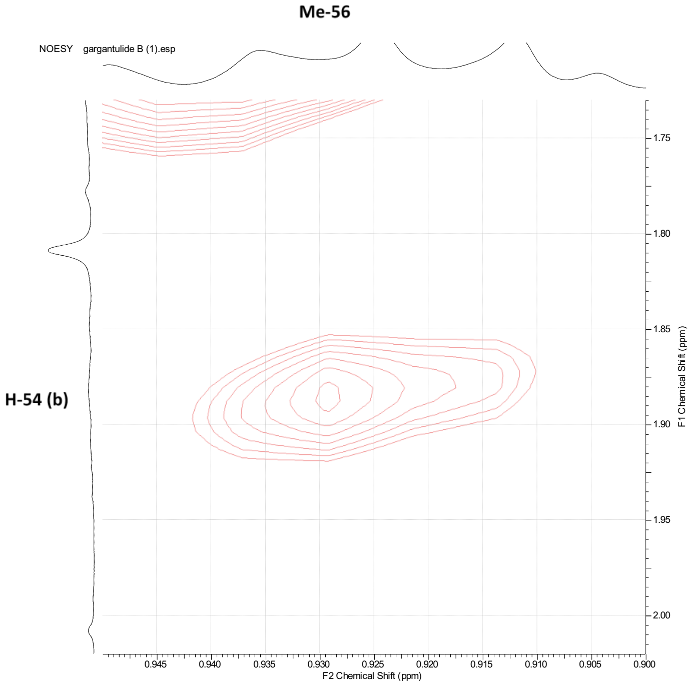


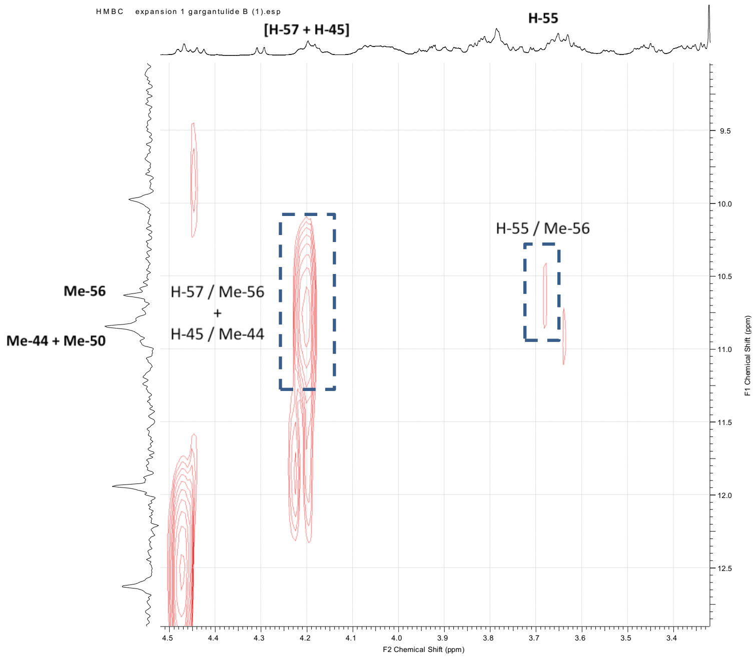


d) NOESY (left) and HMBC (right) expansion supporting the *anti/syn* configuration for C-55/C-56/C-57

**Figure S12.** Determination of the relative configuration of the C-6–C-7 stereocluster for gargantulide B (**1**)

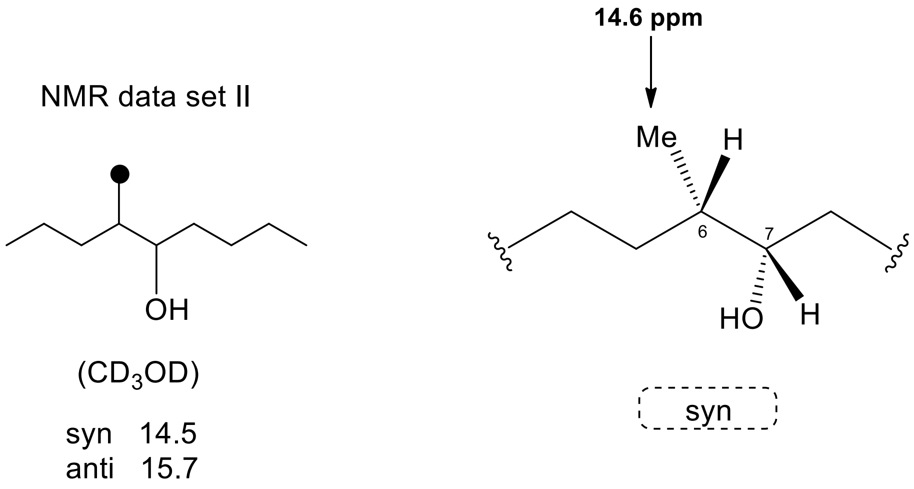


Application of Kishi´s NMR data set II for 4-methylnonan-5-ol isomers (16).

**Figure S13.** Determination of the relative configuration of the C-10–C-12 stereocluster for gargantulide B (**1**)

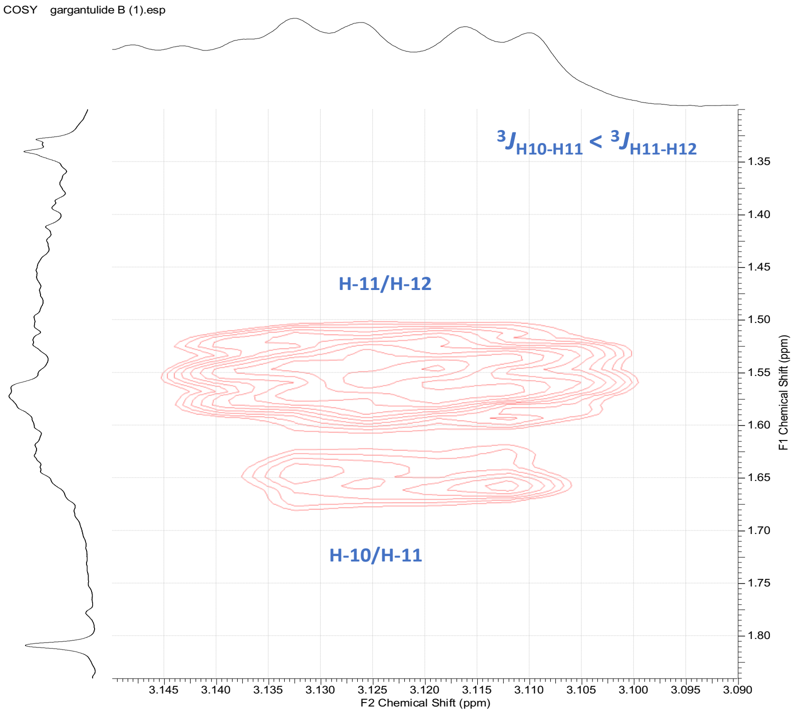


a) Key COSY correlations supporting the *syn/anti* configuration for C-10/C-11/C-12

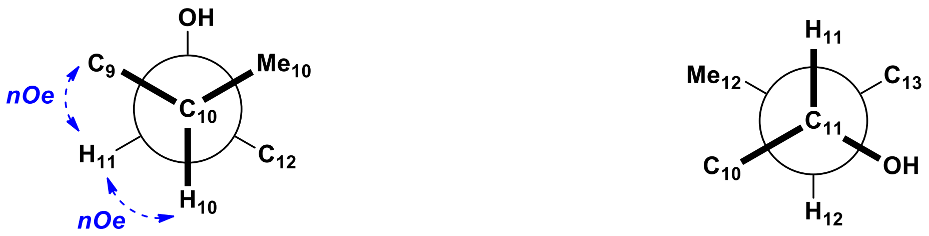


$^3J_{H10\text{-}H11}$ = 3.2 Hz

$^3J_{Me10\text{-}H11}$ = ca. large

$^3J_{C9\text{-}H11}$ = ca. small

$^3J_{C12\text{-}H10}$ = ca. small

$^3J_{H11\text{-}H12}$ = 8.2 Hz

$^3J_{Me12\text{-}H11}$ = ca. small

$^3J_{C10\text{-}H12}$ = ca. small

$^3J_{C13\text{-}H11}$ = ca. small

b) Qualitative $^3J_{C,H}$-based configuration analysis of the C-10 to C-12 stereocluster. Key NOESY correlations are also indicated (although critical overlapping of proton NMR signals corresponding to Me-10 and Me-12 make impossible to distinguish the key NOESY correlation H-10/Me-12 from the genuine cross-peak H-10/Me-10, the clear absence of NOESY correlations between H-10 and both H-13 further supports the configurational proposal)

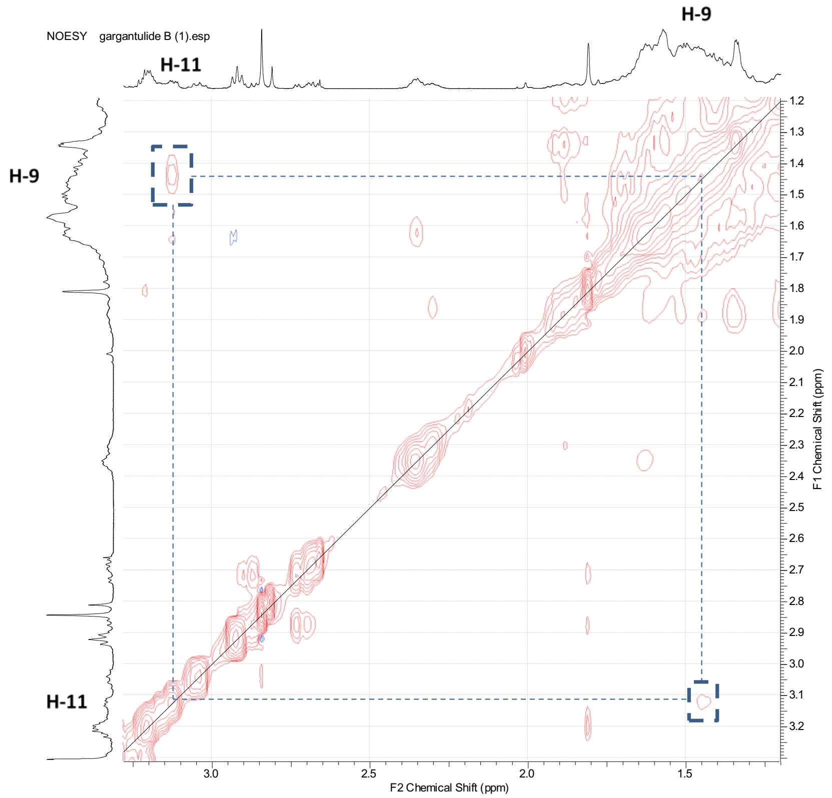


c) Key NOESY correlation supporting the *anti* configuration for C-10–C-12 stereocluster

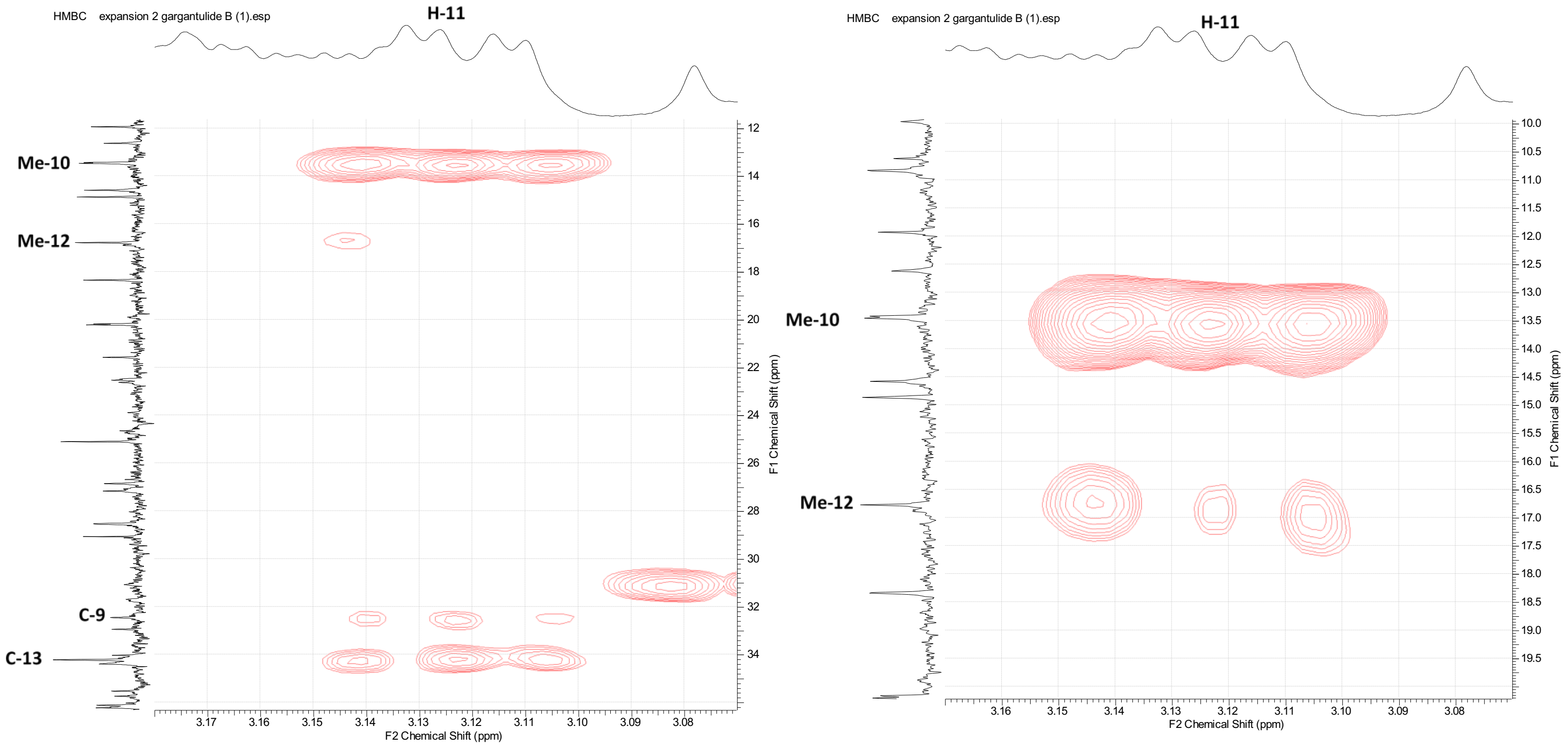


d) HMBC correlations supporting the *anti* configuration for C-10–C-12 stereocluster

**Figure S14.** Determination of the relative configuration of the C-17(R)–C-18(R) stereocluster for gargantulide B (**1**)

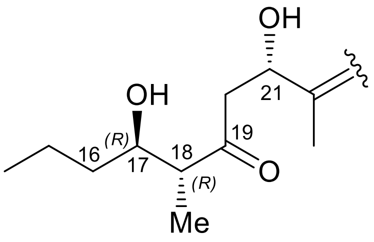


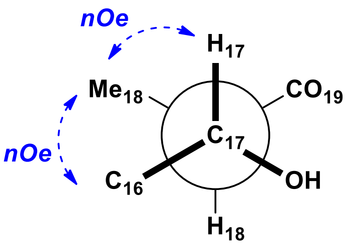


*anti* relative configuration

17(R), 18(R) abs. configuration

$^3J_{H17\text{-}H18}$ = 8.6 Hz

$^3J_{Me18\text{-}H17}$ = ca. small

$^3J_{C16\text{-}H18}$ = ca. small

$^3J_{C19\text{-}H17}$ = ca. small

a) Qualitative $^3J_{C,H}$-based configuration analysis of the C-17–C-18 stereocluster

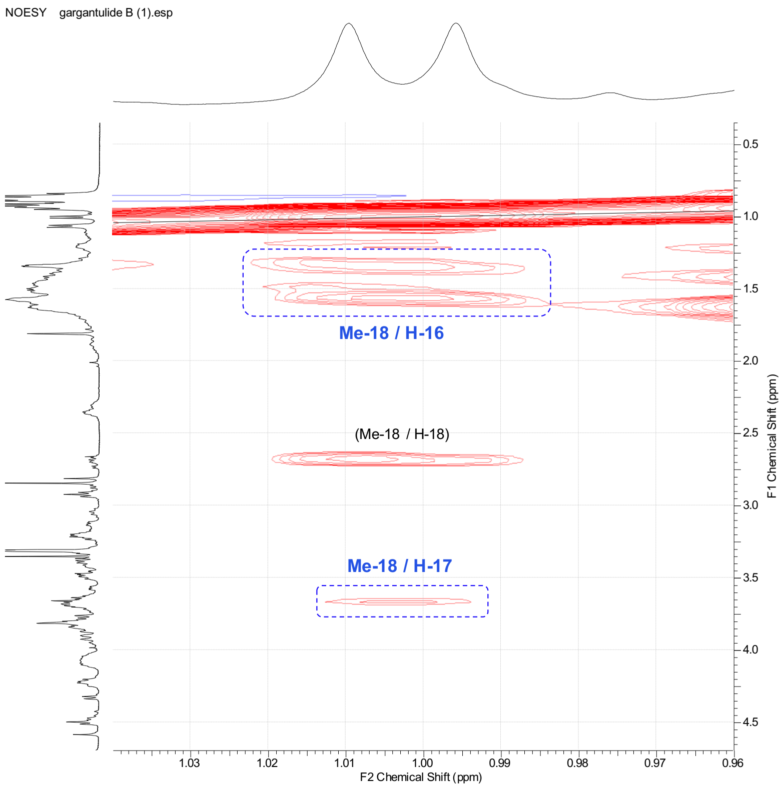


b) Key NOESY correlations supporting the anti configuration for C-17–C-18 stereocluster

**Figure S15.** Determination of the absolute configuration of the non-amino sugars present in gargantulides B (**1**) and C (**2**)

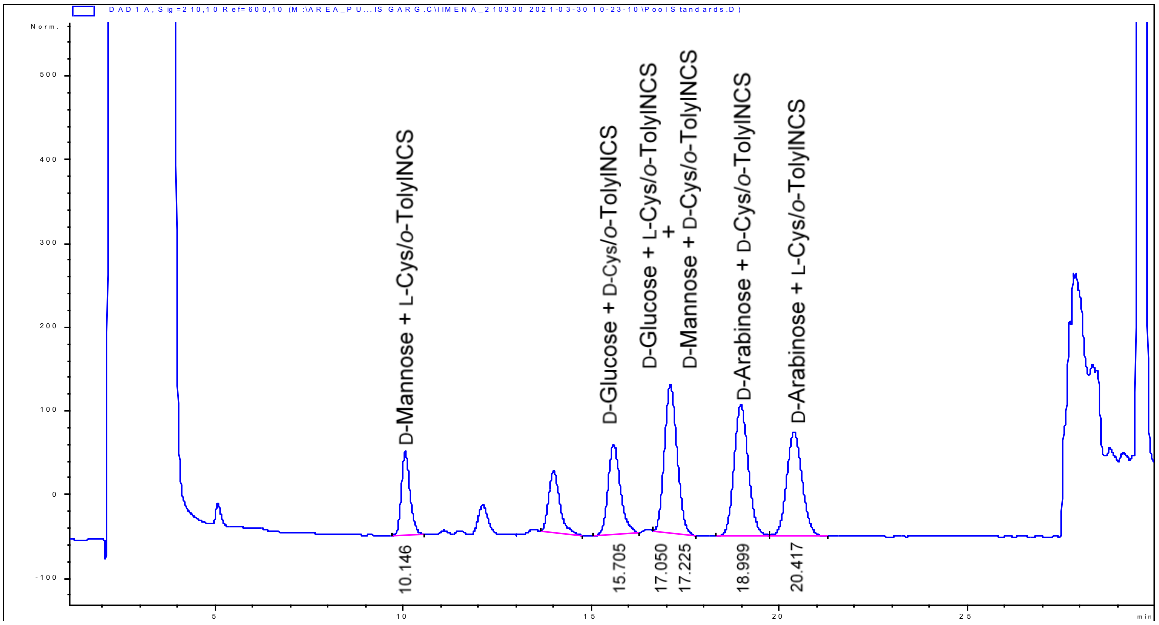


a) LC-UV chromatogram of a mixture of both L- and D-cysteine methyl ester hydrochloride / *o*-tolyl isothiocyanate derivatization reactions of D-Mannose / D-Glucose / D-arabinose standard monosaccharides.

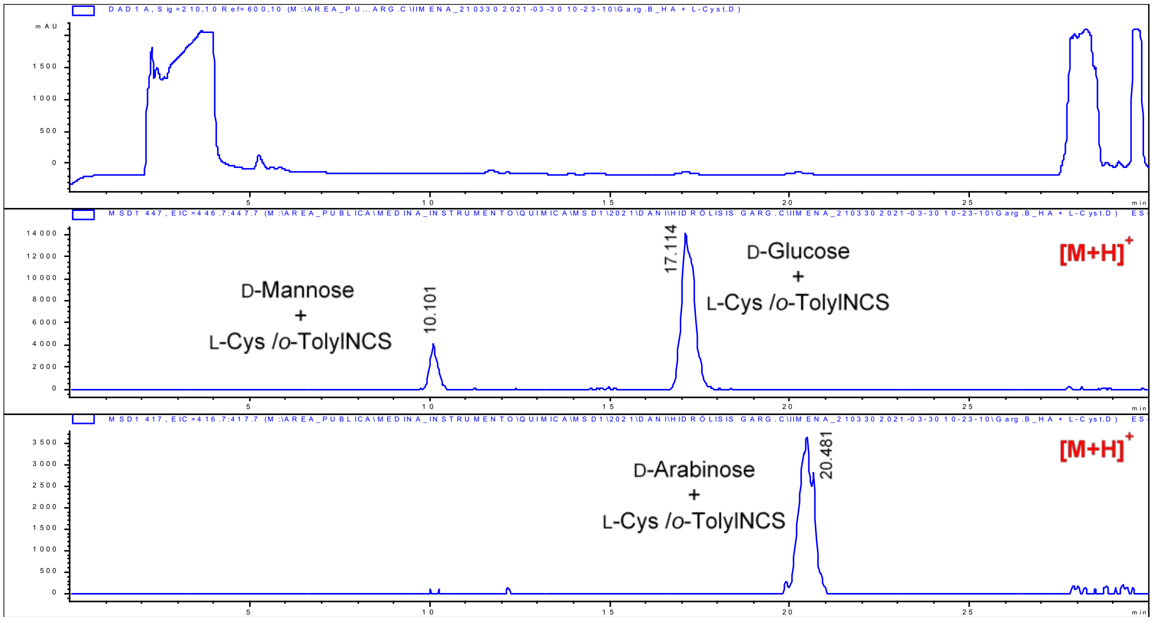


b) LC-UV and extracted-ion chromatograms (EIC) of L-cysteine methyl ester hydrochloride / *o*-tolyl isothiocyanate derivatized hydrolysis of **1** with HCl. D-Mannose, D-Glucose and D-Arabinose were found in **1**

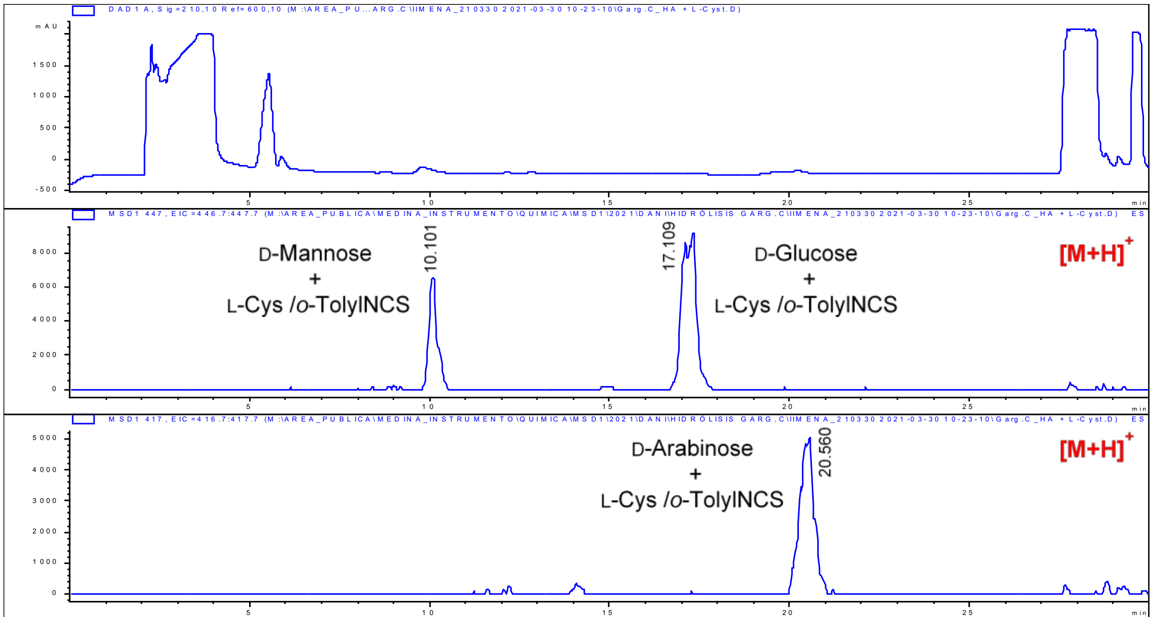


c) LC-UV and extracted-ion chromatograms (EIC) of L-cysteine methyl ester hydrochloride / *o*-tolyl isothiocyanate derivatized hydrolysis of **2** with HCl. D-Mannose, D-Glucose and D-Arabinose were found in **2**

**Table S9.** Antibacterial and antifungal activities of compounds **1** and **2**.

| | | MIC (µg/mL) | | | | | |
|---|---|---|---|---|---|---|---|
| **Microbial strain** | **Strain number** | **(1)** | **(2)** | **R** | **A** | **Am** | **V** |
| *A. baumannii* | MB5973 | 16-32 | 16-32 | 2-4 | | | |
| *P. aeruginosa* | MB5919 | >128 | >128 | | 1 | | |
| *E. coli* | ATCC 25922 | >128 | >128 | | 0.5-0.25 | | |
| *K. pneumoniae* | ATCC 700603 | >128 | >128 | | >16 | | |
| MRSA | | 4-8 | 2-4 | | | | 2-4 |
| MSSA | | 2-4 | 2-4 | | | | 1 |
| VRE | | 2-4 | 1-2 | | | | >128 |
| *A. fumigatus* | ATCC46645 | 32-64 | 32-64 | | | 4 | |
| *C. albicans* | ATCC64124 | >128 | >128 | | | 4 | |

*Positive controls: R Rifampicin, A Aztreonam, Am Amphotericin B, V Vancomycin
